# Gene duplication dynamics and regulatory evolution shape the diversification of Asteraceae

**DOI:** 10.1101/2025.10.29.685401

**Authors:** Erika R. Moore-Pollard, Paige A. Ellestad, Brannan R. Cliver, Jakub Baczyński, Matthew D. Pollard, Zachary Meharg, Andrew C. Willoughby, Samantha Drewry, Alex Harkess, J. Mauricio Bonifacino, Zachary L. Nimchuk, John M. Burke, Daniel S. Jones, Jennifer R. Mandel

## Abstract

The flowering plant order Asterales exhibits a striking disparity in species richness, with >30,000 species in Asteraceae compared to <50 in its sister family Calyceraceae. To investigate the genomic basis of this imbalance, we assembled three new chromosome-level genomes, including the first for Calyceraceae, and re-annotated five additional genomes. Comparative analyses revealed exceptionally high repeat content in both Asteraceae and Calyceraceae, pervasive chromosomal rearrangements, and evidence for shared and lineage-specific WGDs. In Asteraceae, tandem and dispersed duplications disproportionately drove expansions of gene families linked to secondary metabolism and stress response, while segmental duplicates bore signatures of adaptive selection for the regulation of biosynthetic and metabolic processes. Selective pressures on flowering time regulators suggest an evolved balance between regulatory flexibility and developmental constraint in floral diversification. These patterns reveal that, beyond ancient polyploidy, small-scale duplications and selective fine-tuning of regulatory networks underpinned the ecological versatility in Asteraceae, fueling its extraordinary diversification.

## Main

A recurring pattern in flowering plant evolution is the uneven distribution of species richness across the angiosperm tree of life, whereby species-poor lineages are often sister to highly diverse ones. While extinction and gaps in the fossil record may obscure some histories, the overall pattern remains clear^1^. Nowhere is this contrast more striking than in Asterales. The order includes ∼35,800 species across 11 families, with >30,000 species belonging to Asteraceae alone^2,3^. Accounting for ∼10% of all flowering plants and spanning nearly every continent and ecological niche (Fig. 1a), Asteraceae ranks among the most diverse and economically important plant families worldwide^4–10^. Over 2,000 of its members are used by humans for food, medicine, and ornamentals, yet it also ranks as the second most invasive plant family, exacting heavy ecological and economic costs^7,11^. In sharp contrast, its sister family, Calyceraceae, comprises fewer than 50 species confined to southern South America (Fig. 1a)^12–14^. Both Asteraceae and Calyceraceae originated in South America during the late Cretaceous^8,9^, but only Asteraceae subsequently diversified extensively and dispersed globally^7^. This extreme disparity raises a central question: what genomic innovations fueled Asteraceae’s extraordinary radiation?

**Fig. 1.**
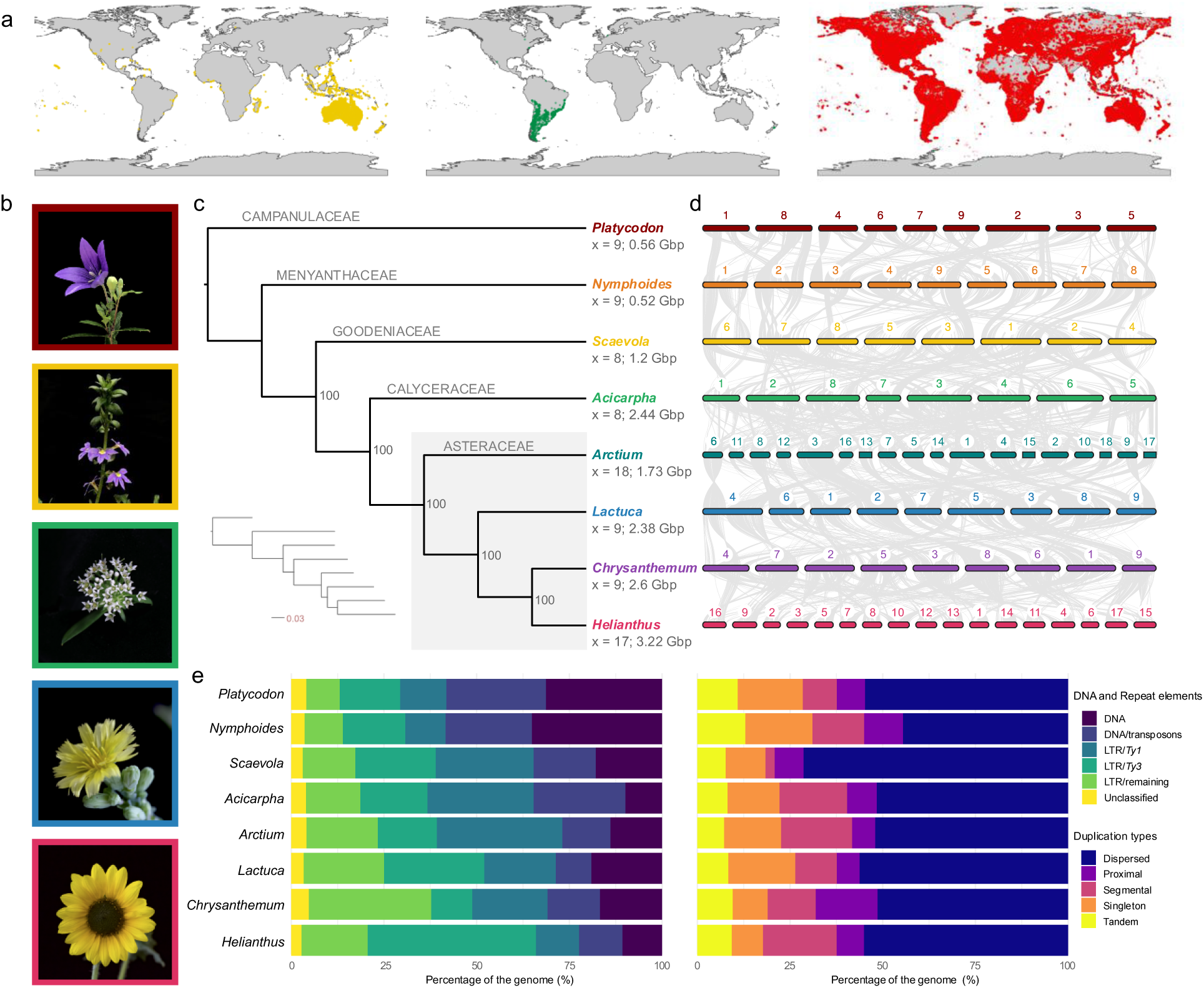
Genomic characteristics of the eight focal Asterales species. **a)** from left to right, geographic distributions of Goodeniaceae (yellow), Calyceraceae (green), and Asteraceae (red), based on occurrence points of preserved specimens accessed from the Global Biodiversity Information Facility (accessed October 8, 2025)^168–170^. **b)** Photos of some tested taxa. Colored box around image matches species colors in **c** and **d**. From top to bottom: *Platycodon grandiflorus* (Campanulaceae); *Scaevola aemula* (Goodeniaceae); *Acicarpha procumbens* (Calyceraceae); *Lactuca sativa* (Asteraceae); *Helianthus annuus* (Asteraceae). Image of *Acicarpha procumbens* by D.S. Jones, remaining images by A.C. Willoughby. **c)** Nuclear phylogenetic cladogram of the eight species generated via the multiple-species alignment from OrthoFinder and IQTree. Values at nodes are bootstrap support values. Tree to the bottom left shows branch lengths by nucleotide substitutions per site. The four Asteraceae taxa are shaded gray. Values under each species name at the tip label correspond to the estimated hapoloid chromosome number (x) and genome assembly size in giga-base pairs (Gbp), respectively. **d)** Synteny plots generated with JCVI ordered by phylogenetic placement of the eight target taxa. Each bar is representative of a single chromosome, with circled numbers above the bar indicating chromosome number. Gray lines between species’ chromosomes indicate a single syntenic gene block. **e)** Barplots indicating the percentage of DNA and repeat elements (left) and gene duplication type (right) across the genomes with percentage of the genome on the x-axis and species on the y-axis. Colors correspond to the legends to the right. Taxon order follows phylogenetic placement from **c.**

Asteraceae boasts a long evolutionary history shaped by rapid radiation, polyploidization, hybridization, widespread ecological adaptation, and morphological innovation^5–9^. Its extensive diversification has been largely attributed to enhanced reproductive success through pollinator attraction to its unique head-like inflorescence composed of many individual florets, the capitulum^6,15,16^. Although Calyceraceae inflorescences also resemble capitula, they did not spur analogous radiations. Key differences distinguishing the two families include the absence/presence of a terminal flower, un-fused/fused anthers, free/partially fused filaments attached to the corolla (Asteraceae versus Calyceraceae, respectively), and variation in secondary pollen presentation mechanisms^15,17,18^. Additionally, the production of diverse phytochemicals in Asteraceae facilitated ecological adaptation, particularly through defense responses^9^. These traits, coupled with a diversity of capitulum forms derived from distinct morphological and organizational modifications (Fig. 1b)^16^, likely contributed to the disparity in species richness between the two families, though the genomic mechanisms underlying their origin have yet to be elucidated.

Acting across multiple scales, from individual genes to entire genomes, duplications have been a major driver of plant diversification by providing the raw genetic material for functional innovation^19^. Among these processes, whole-genome duplications (WGDs) are particularly pervasive throughout the evolutionary history of angiosperms and are widely recognized as a central force contributing to both their expansive diversity and widespread distribution^20–22^. The triplicate genome structure of Asteraceae suggests that the family’s diversification was catalyzed by two successive WGD events (i.e., paleohexaploidy) at its core which was then followed by lineage-specific WGDs in some of its most species-rich groups, e.g., tribe Cichorieae and the Heliantheae alliance^23–26^. Although limited by available genomic data, evidence also suggests that the sister family Calyceraceae experienced the same paleohexaploidy event^4–6^. This finding raises further questions about the role of mechanisms beyond WGDs in plant diversification. For example, transposable elements (TEs) represent another dynamic force in plant diversification, driving genomic change through chromosomal rearrangements and altering gene expression via insertions or chromatin remodeling^27–29^. Asteraceae genomes, notable for their exceptionally high repeat content dominated by TEs, highlight the likely role of TE activity in promoting lineage-specific adaptations to diverse environments^30^. While both WGDs and TE-mediated small-scale duplications have profoundly shaped plant evolution, their relative contributions to diversification remain unresolved. Resolving the precise phylogenetic placement of these events within Asterales is key to understanding the enormous evolutionary success of Asteraceae.

Despite the evolutionary and economic importance of Asteraceae, our understanding of genome evolution in the family, and in the broader Asterales order, remains limited due to sparse genomic and transcriptomic resources, beyond a handful of well-studied species (e.g., *Helianthus*^31^ and *Lactuca*^26^). To date, assembled genomes are mostly concentrated to the highly derived clades of Asteraceae with no representation of the early South American lineages^32^. Similarly, the small sister family, Calyceraceae, lacks genomic data, leaving a major gap in our understanding of the origin and early evolution of Asteraceae.

To investigate the genomic drivers of diversification across Asterales, we generated three new chromosome-level assemblies of *Acicarpha procumbens* (the first Calyceraceae genome), *Scaevola aemula* (Goodeniaceae), and *Platycodon grandiflorus* (Campanulaceae), and re-annotated five publicly available genomes in Menyanthaceae (*Nymphoides indica*) and Asteraceae (*Arctium lappa*, *Lactuca sativa*, *Chrysanthemum lavandulifolium*, and *Helianthus annuus*) (Fig. 1c). In doing so, we address a fundamental evolutionary question: How have WGDs and small-scale gene duplications contributed to the origin and expansion of species diversity within Asterales? Our analyses highlight lineage-specific patterns shaped by genome duplications, chromosomal rearrangements, gene family dynamics, and signatures of selection to provide a comparative framework for understanding the genomic basis of Asteraceae’s remarkable diversification.

## Results

### Extensive genome rearrangements yet similar composition in Asteraceae and Calyceraceae

We assembled chromosome-level genomes of *Platycodon*, *Scaevola*, and *Acicarpha*, with *Scaevola* and *Acicarpha* assemblies being haplotype-resolved. The assemblies were highly contiguous and complete based on BUSCO analysis, and comparable with the re-annotated genomes (range = 96.6%–98%, mean = 97.6%; Supplementary Figs. 1 and 2, Supplementary Tables 1 and 2). Genome rearrangements were widespread, with variations in chromosome number indicating multiple chromosomal fission/fusion events across all Asterales taxa (Fig. 1c,d).

As expected^30^, repeat content was high in Asteraceae (range = 80.9%–89.3%)^28^, though similarly elevated in *Scaevola* (Goodeniaceae, 82.1%) and *Acicarpha* (Calyceraceae). Unexpectedly, *Acicarpha* exhibited the highest repeat proportion, reaching 90.1% (Fig. 1e, Supplementary Table 2), with *Ty1* being the most abundant repeat type, comprising 28.63% of the genome. *Arctium* had the highest and *Nymphoides* had the lowest *Ty1* content overall (33.82% and 10.84%, respectively). In contrast, *Helianthus* exhibited the highest proportion of *Ty3* elements (45.4%), while *Chrysanthemum* had the lowest (11.1%). Except for *Chrysanthemum*, the proportions of *Ty1*/*Ty3* retroelements show inverse patterns across the phylogeny, with earlier-diverging species having higher percentages of *Ty1* and lower *Ty3* (e.g., *Acicarpha*), while more recently derived taxa show the opposite (e.g., *Helianthus*; Fig. 1e), a pattern reported previously^30^.

The proportions of duplicated gene types varied across Asterales genomes (Fig. 1d). Segmental duplicates, likely derived from WGDs^33^, were most common in *Acicarpha* (18.2%), *Arctium* (19.1%), and *Helianthus* (19.8%) and least in *Scaevola* (2.5%). In contrast, small-scale duplicates (tandem, proximal, and dispersed), likely reflecting localized transposition or recombination events^33^, were most frequent in *Nymphoides* (12.9%) and showed modest variation across other genomes (range = 7.6–10.8%, mean = 9.2%). Overall, *Acicarpha*’s duplication frequencies fell within the range observed across Asteraceae (Fig. 1e).

### Shared and lineage-specific WGD in Asteraceae and Calyceraceae

Asteraceae is hypothesized to have experienced one to multiple paleopolyploidization events during its evolution^4–6,25,26^, though the phylogenetic position of WGDs within the family and related lineages remains uncertain. Syntenic gene depths corroborate the presence of ancient WGDs for all taxa assessed, ranging from two (*Platycodon, Nymphoides,* and *Scaevola*) to five (*Helianthus*), as compared to *Vitis vinifera* (an ideal reference due to its lack of WGD since the gamma [γ] event shared by all eudicots^34^; Fig. 2a, Supplementary Table 3). In Calyceraceae, we observed syntenic duplications and, to a lesser extent, triplications between *Acicarpha* and *Scaevola* (Calyceraceae versus Goodeniaceae), and between *Acicarpha* and *Arctium* (Calyceraceae versus Asteraceae), indicating lineage-specific duplications within Asteraceae and Calyceraceae (Fig. 2b, Supplementary Table 4).

**Fig. 2.**
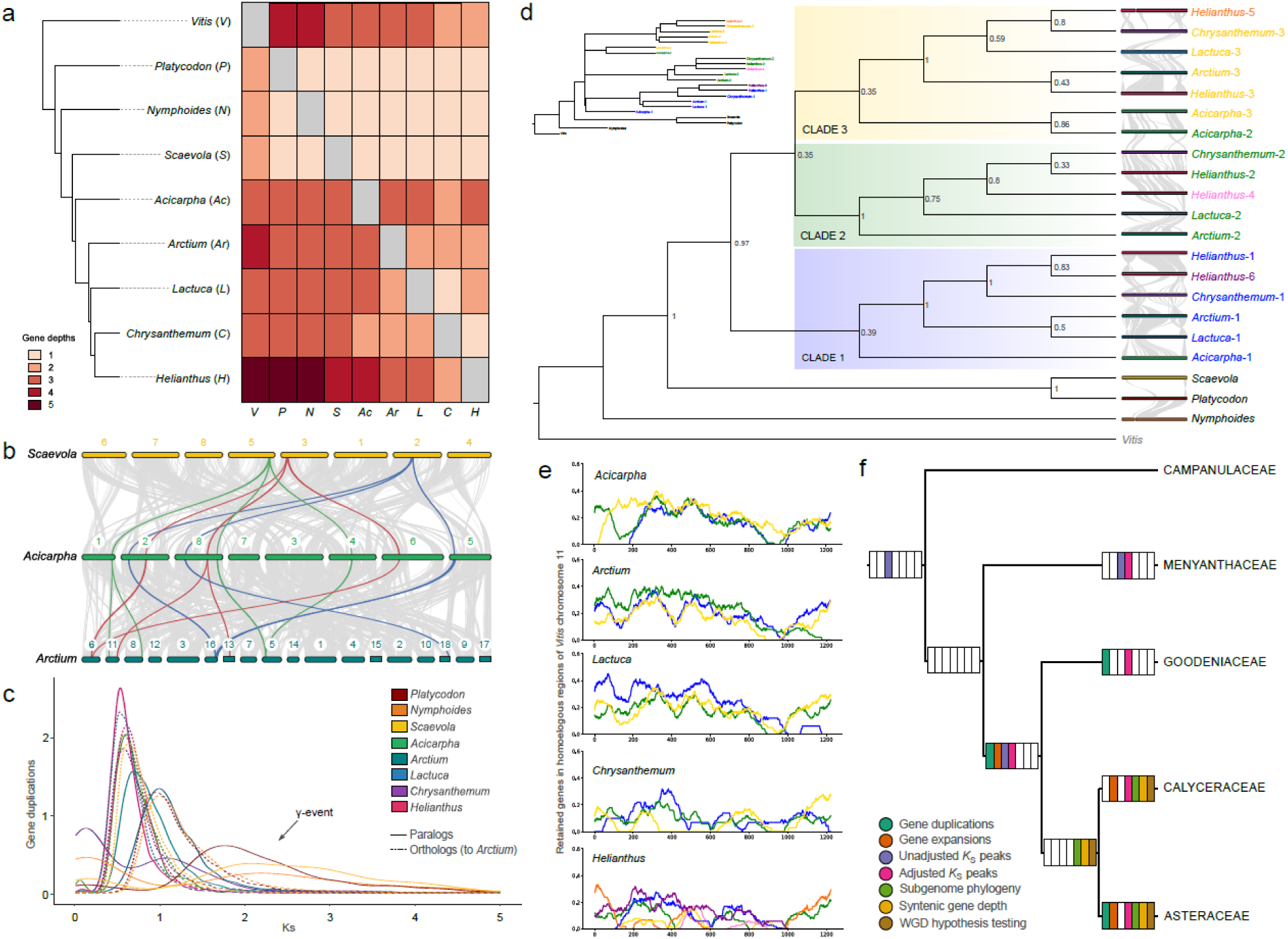
Evidence of WGD events across the eight tested Asterales taxa. **a)** Pairwise gene depth ratios from JCVI between the eight Asterales species plus *Vitis*. The top right portion of the matrix indicates the gene depth ratio of the y-axis, while the bottom left portion is for the x-axis (e.g., *Scaevola:Acicarpha* is 1:3). Colors follow the legend in the bottom left of the panel. **b)** Syntenic regions between *Scaevola* (Goodeniacae), *Acicarpha* (Calyceraceae), and *Arctium* (Asteraceae) and as indicated by JCVI. Colored synteny ribbons (green, red, and blue) are identified gene triplications between *Scaevola* to *Acicarpha* and follow those genes to *Arctium*. Bars represent chromosomes with circled numbers above the bars representing the chromosome number. **c)** *K*_S_ plot of paralogous gene pairs and orthologous gene pairs against *Arctium* for each species tested. Peaks closer to zero indicate a younger duplication event. Colors of lines correspond to the legend at the top right. The predicted gamma event is indicated with the arrow. **d)** *Vitis* chromosome 11 subgenome phylogeny as determined by WGDI. Potential duplication events are indicated by clade, from Clades 1-3. Species names at the tips are colored according to subgenome number: blue: 1, green: 2, yellow: 3, pink: 4, orange: 5, purple: 6. Synteny plots at each tip were generated with JCVI and colored according to the synteny plots of Figure 1d. Local posterior probability (LPP) support values are at each node. **e)** Genome fractionation results from the subgenome analysis of *Vitis* chromosome 11 from WGDI for *Acicarpha* and four Asteraceae subgenomes. Gene retention is on the y-axis and base pair number on the x-axis. Colors in the graph correspond to the subgenome numbers which are also represented as the tip label color in **d**. **f)** Summary of WGD results from across this study. Colored boxes indicate the findings, as indicated by the legend, that provide evidence for WGDs on that branch. White boxes indicate no evidence was found for that analysis. Campanulaceae has no boxes as WGDs were not directly tested for that family.

Comparing synonymous substitution rate (*K*_S_) distributions across Asterales, we inferred multiple duplications: an ancient WGD around *K*_S_ = 2 (likely the γ event); a WGD shared by Goodeniaceae, Calyceraecae, and Asteraceae; small-scale duplications in *Scaevola* and *Acicarpha*; and separate WGDs in *Chrysanthemum* and *Helianthus* (Fig. 2c, Supplementary Figs. 3 and 4). Alternatively, using *K*_S_ distributions adjusted to the rate of a focal taxon, we found: three small-scale duplications in *Nymphoides*; a WGD shared by Goodeniaceae, Calyceraceae, and Asteraceae; two WGDs in *Scaevola*; two WGDs in *Acicarpha*; one WGD at the crown of Asteraceae; one WGD in *Lactuca*; two in *Chrysanthemum*; and one in *Helianthus.* Prior to the divergence of Asteraceae, both analyses indicated a WGD shared by Goodeniaceae, Calyceraceae, and Asteraceae, but only the rate-adjusted method identified the Asteraceae-specific WGD (Supplementary Figs. 5 and 6).

To further investigate WGDs, we phased the genomes and examined relationships among duplicated subgenomes. Using phased sub-genomes that retained the most collinear genes homologous to *Vitis* (Supplementary Datasets 1 and 2; Supplementary Figs. 7-13), phylogenetic analyses revealed that *Acicarpha* and Asteraceae generally share at least one duplication event, while each lineage also experienced independent duplications (Fig. 2d). These findings were further supported by gene fractionation results, which indicated that *Acicarpha* sub-genomes differ in age from those of Asteraceae, and are generally more recent, although this trend was not always consistent (e.g., patterns of gene fractionation are distinct in *Lactuca* but overlap in *Chrysanthemum*; Fig. 2e).

Given the variation in WGD signals across analyses, we performed hypothesis testing to better place WGDs under various models. Under the critical model^35^, which allows duplication and loss rates to vary among branches, we found support for WGDs before and after the Calyceraceae-Asteraceae split and lineage-specific events within *Acicarpha*, *Chrysanthemum*, and *Helianthus*. Under the relaxed model^35^, which allows rate heterogeneity among and within branches, we found support for events in all tested lineages (Supplementary Fig. 14). These results propose multiple WGD scenarios (Figs. 2f and 3), underscoring the difficulty of placing Asteraceae WGDs. Some findings support earlier work that inferred two events: one shared between Calyceraceae and Asteraceae and a second, independent duplication in each lineage (Fig. 2f)^36^. However, other analyses support an older event predating the Goodeniaceae-Calyceraceae-Asteraceae split followed by lineage-specific duplications in Asteraceae and Calyceraceae (Fig. 2f), a pattern not previously reported.

**Fig. 3.**
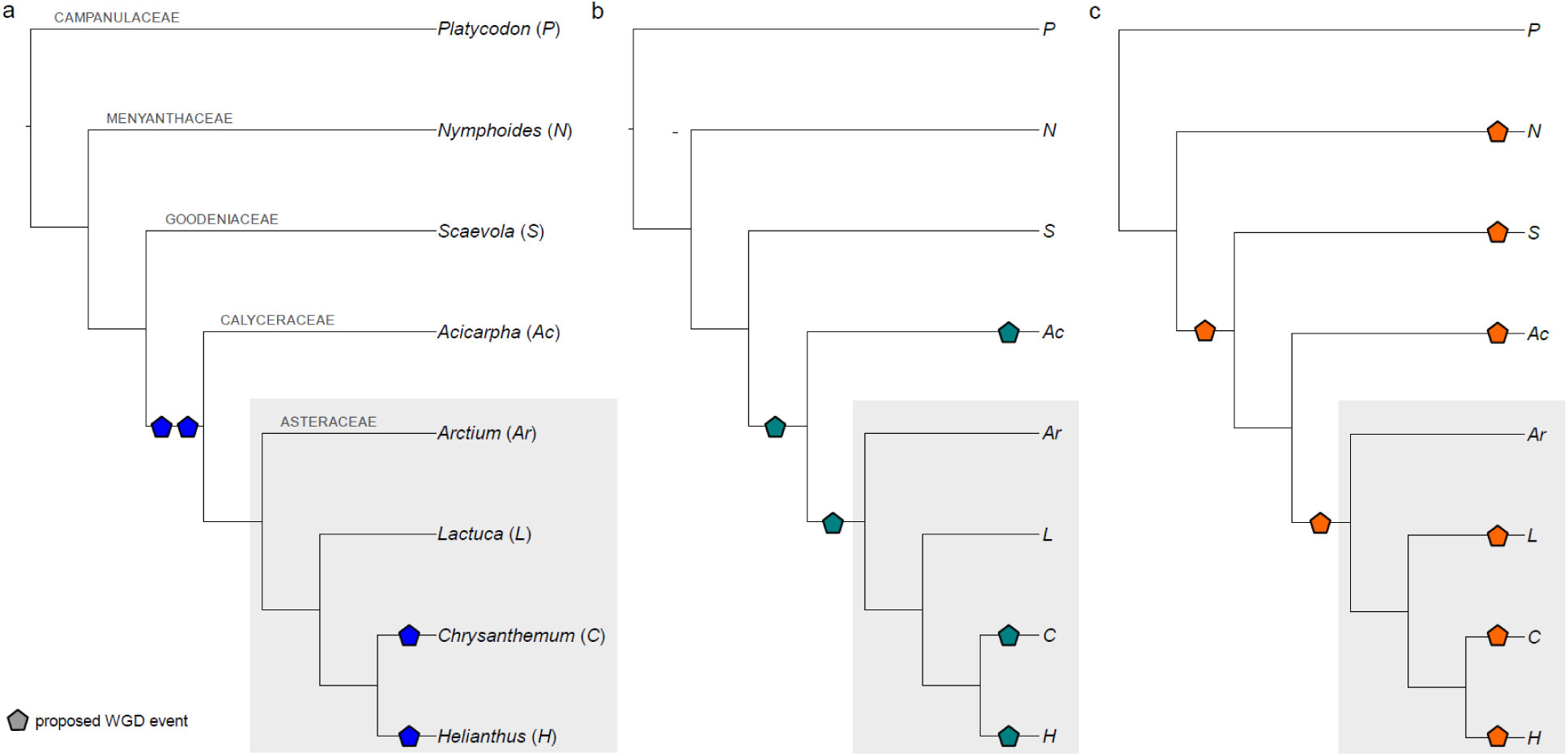
Possible WGD scenarios in Asterales. A colored pentagon indicates a predicted WGD placement. The scenarios are as follows: **a)** hypothesis for WGT at the crown of Asteraceae and Calyceraceae, followed by unique WGDs in *Chrysanthemum* and *Helianthus* lineages, as supported by previous literature, **b)** hypothesis for WGD at the crown of Asteraceae and Calyceraceae, followed by WGD at the base of Asteraceae, then unique WGD in *Acicarpha, Chrysanthemum,* and *Helianthus* lineages, supported by the critical model in WHALE, **c)** hypothesis for WGD at the crown of Asteraceae, Calyceraceae, and Goodeniaceae, as well as unique WGD in *Nymphoides, Scaevola, Acicarpha, Lactuca, Chrysanthemum,* and *Helianthus* lineages, supported by the relaxed model in WHALE.

### Asteraceae gene family expansions comprised of mostly tandem duplicates

To assess how duplications contributed to Asteraceae diversification, we examined gene orthogroup expansions across the family. Orthogroups that underwent substantial expansion at the base of Asteraceae (n = 115; Supplementary Fig. 15) were significantly enriched in processes related to stress response, signaling, and metabolic regulation, including cellular response to amino acid stimulus, sesquiterpene biosynthesis, and cell surface receptor signaling (Fig. 4a, Supplementary Fig. 16). Among the most expanded gene families were those related to stress response and signaling (e.g., *GLP9*^37^ and *WAKL2*^38^; Table 1), and to biosynthesis of primary and secondary metabolites (e.g., *FAD2*, involved in fatty acid synthesis^39^; and *CYP71B23*, involved in sesquiterpene synthesis and modification^40^; Table 1). These were primarily tandem duplicates (30%), followed by dispersed (28%), proximal (23%), and then segmental (18%) duplicates (Supplementary Fig. 17). Alternatively, *Acicarpha* expansions were not enriched in macromolecule biosynthesis, but in pathways responding to hormones like auxin and environmental stimuli such as wounding or organic substances (Fig. 4b). Of these, multiple orthogroups included genes related to *CRK8*, a cysteine-rich receptor-like kinase typically involved abiotic and biotic stress responses^41^ (Table 1). Over half (62%) of *Acicarpha*’s expanded genes were dispersed duplicates, while 17% were tandem duplicates, and only 4% were segmental duplicates (Supplementary Fig. 17).

**Fig. 4.**
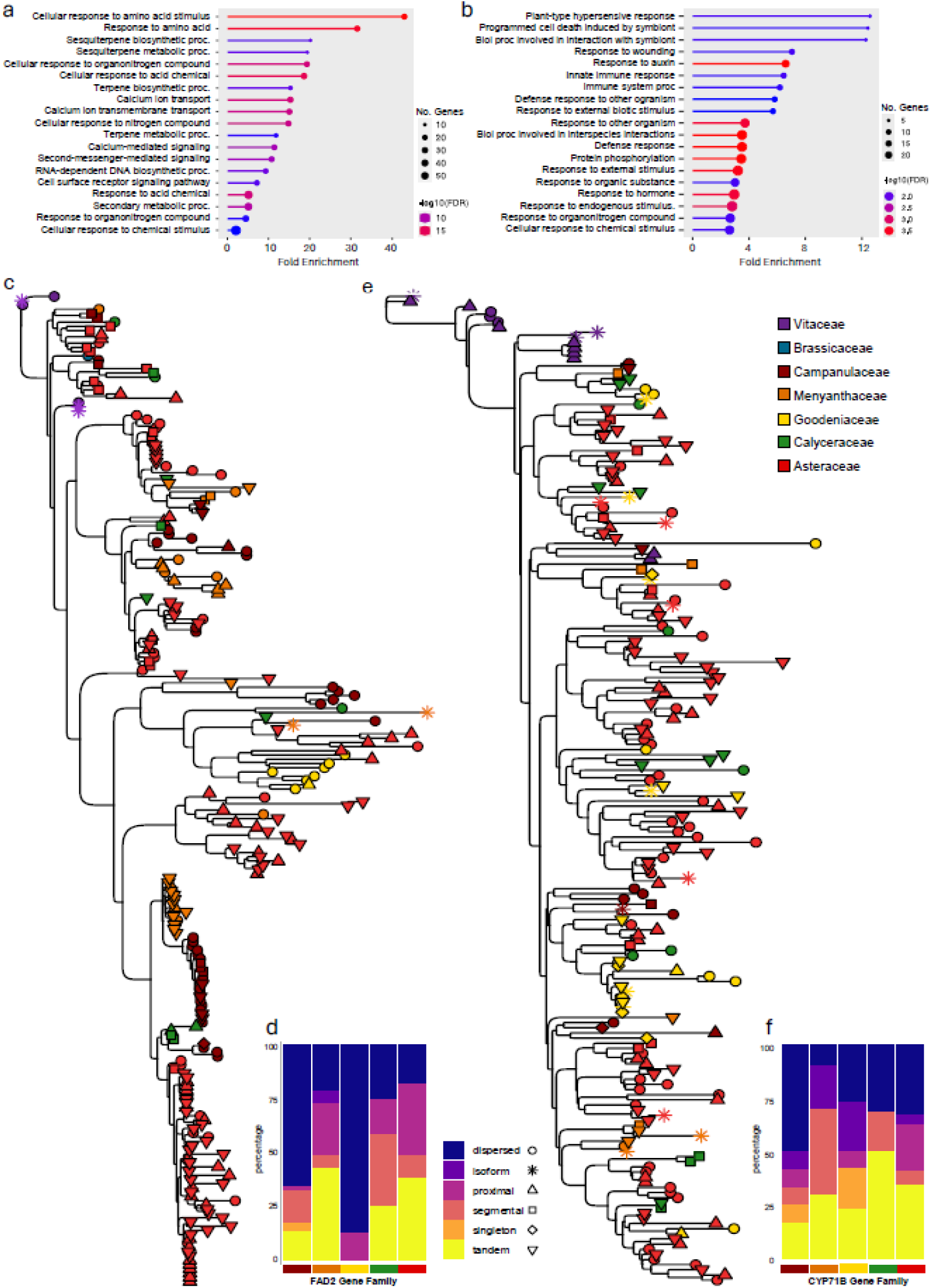
Genes underpinning Asteraceae diversification. GO enrichment of biological processes associated with expanded orthogroups in **a)** Asteraceae and **b)** *Acicarpha;* **c)** Expanded *FAD2* gene family tree where tip colors represent the taxonomic family of each gene as indicated by the legend at the right and tip shapes represent the duplication type of each gene as indicated by the legend in **d; d)** Composition of each duplication type by taxonomic family for the *FAD2* gene family; **e)** Expanded *CYP1B23* gene family tree with same tip shape and color designations as **c**. **f)** Composition of each duplication type by taxonomic family for the *CYP1B23* gene family.

**Table 1.** Representative genes from analyses aiming to understand the genomic mechanisms underpinning diversification in Asterales. Gene symbols were obtained from BLASTP search results to a TAIR10 protein database. Functions were summarized from representative publications. Asterisk indicates that this gene was the best match for six different orthogroups. Complete results are listed in Supplementary Dataset 3.

| Category | Gene | Description | Involved in |
| --- | --- | --- | --- |
| Top ten expanded orthogroups in Asteraceae | *GLP9* | germin-like protein 9 | disease resistance<sup>37</sup> |
|  | *WAKL2* | wall associated kinase-like 2 | cellular communication, disease resistance<sup>38</sup> |
|  | *FAD2* | fatty acid desaturase 2 | fatty acid synthesis<sup>39,42</sup> |
|  | *CYP71B23* | cytochrome P450, family 71, subfamily B, polypeptide 37 | sesquiterpene synthesis and modification<sup>40</sup> |
|  | *AT1G43760* | DNAse I-like superfamily protein | NA |
|  | *RPP13* | NB-ARC domain-containing disease resistance protein | disease resistance<sup>155</sup> |
|  | *WIP4* | Wound-responsive family protein | root development<sup>156</sup> |
|  | *HYR1* | UDP-Glycosyltransferase superfamily protein | growth regulation<sup>157</sup> |
|  | *CYP72A15* | cytochrome P450, family 72, subfamily A, polypeptide 15 | metabolic resistance<sup>158</sup> |
|  | *HTB9* | Histone superfamily protein | chromatin remodeling<sup>159</sup> |
| Top ten expanded orthogroups in *Acicarpa* | *NFU3* | NFU domain protein 3 | plant growth, development, and flowering<sup>160</sup> |
|  | *CRK8\** | cysteine-rich RECEPTOR-like kinase | abiotic and biotic stress response<sup>41</sup> |
|  | *LRR4* | NB-ARC domain-containing disease resistance protein | disease resistance<sup>161</sup> |
|  | *AT2G01050* | zinc ion binding; nucleic acid binding | NA |
|  | *SAUR2* | SAUR-like auxin-responsive protein family | stress response<sup>162</sup> |
| Lost in Goodeniaceae, Calyceraceae, and Asteraceae | *GLB1(PII)* | Nitrogen regulatory protein P-II homolog | nitrogen and metabolic regulation<sup>23</sup> |
| Positive Selection in Asteraceae | *CYP72A15* | cytochrome P450, family 72, subfamily A, polypeptide 15 | secondary metabolism<sup>163</sup> |
|  | *TMAC2* | AFP2 (ABI five-binding protein 2) family protein | stress response<sup>47</sup> |
|  | *TCP2,5,24* | TEOSINTE BRANCHED 1, cycloidea and PCF transcription factor 2,5,24) | plant growth and development<sup>50,51,55</sup> |
|  | *WRKY6,21,22,27,31,39,42,57,74,75* | WRKY DNA-binding proteins | stress response<sup>164</sup> |
|  | *GI* | gigantea protein | flowering time regulation<sup>56,57</sup> |
|  | *FVE* | Transducin family protein / WD-40 repeat family protein | regulation of flowering time and cold response<sup>165</sup> |
|  | *CGA1* | cytokinin-responsive gata factor 1 | regulation of chloroplast development, stress response, and flowering time<sup>60,166</sup> |
|  | *FD* | Basic-leucine zipper (bZIP) transcription factor family protein | regulation of floral transition, plant development, and environmental response<sup>167</sup> |
|  | *AGL24* | AGAMOUS-like 24 | regulation of flowering time<sup>62</sup> |

The Asteraceae-specific expansion of *FAD2* homologs is hypothesized to have catalyzed functional divergence^42,43^ and supported the evolution of a unique N-C balance system in conjunction with the loss of *PII,* a regulator of nitrogen-carbon assimilation^23^. The absence of *PII* in *Acicarpha* and *Scaevola* suggests that this loss predates the Goodeniaceae split, clarifying previous hypotheses about its timing (Fig. 4c, Supplementary Dataset 3). Duplication patterns consisting of a few segmental genes, with much larger proportions of proximal and tandem genes, mirrored previous research suggesting that this expansion in Asteraceae arose via tandem duplications following genome duplications^23^ (Fig. 4c,d, Supplementary Fig. 18), contrasting with Calyceraceae which has fewer tandem and proximal duplicates (Fig. 4d). Also involved in metabolism, the expansion of Asteraceae *CYP71B23* homologs involved few segmental genes, but with higher proportions of tandem, proximal, and dispersed genes (Fig. 4e,f, Supplementary Fig. 19). Overall, these patterns indicate that gene family expansions, particularly those arising through tandem duplications, have shaped metabolic and stress-response diversity in Asteraceae, linking genome duplication and remodeling to the uneven diversification of Asterales.

### Site-specific selection in Asteraceae acting on mostly segmental duplicates

Genes containing positively selected sites in all four Asteraceae taxa (n = 586 orthogroups) were significantly enriched for the regulation of biosynthetic and metabolic processes (Supplementary Fig. 20). Of these genes, segmental and dispersed duplicates comprised 45% and 37%, respectively (Supplementary Fig. 17). In addition to the expansions in the CYP superfamily—a large and functionally diverse gene family involved in secondary metabolite synthesis, hormone regulation, and stress responses^44,45^—we observed positively selected sites in all Asteraceae taxa for orthogroups related to *CYP96A1*, *CYP72A7*, *CYP706A4*, *CYP706A6*, *CYP72A13*, and *CYP72A15* (Supplementary Dataset 3). Previous studies also revealed strong lineage-specific diversification of CYP genes in Asteraceae, indicating rapid evolutionary shifts, likely reflecting a central role in metabolic innovation^46^. To test for lineage-specific positive selection, we used the resulting 586 orthogroups with sites under positive selection in Asteraceae species as input for branch model analyses. Of these, 47 showed evidence for episodic positive selection in an Asteraceae lineage (Supplementary Dataset 3). Genes involved in the predominant biological processes associated with this gene set (stress response and regulation of metabolic processes) include: *TMAC2*, a regulator of abscisic acid and salt stress^47^; *WRKY27,* a transcription factor (TF) involved in plant defense and floral development^48^; and *TCP5*, a TF that controls thermomorphogenesis^49^, regulates ethylene-mediated petal development and growth^50^, and impacts leaf margin patterning^51^ (Table 1). Other class II TCP TFs, *TCP2* and *TCP24*, also exhibiting episodic positive selection in Asteraceae, regulate key developmental transitions and have been linked to floral diversification in sunflower^52^.

### Lineage-specific selection in Asteraceae supports fine-tuning of flowering time

To focus on genes and pathways involved in floral innovation in Asteraceae, we performed branch model selection analyses on a curated set of 335 flowering-associated orthogroups (Supplementary Table 5). We identified 27 orthogroups with episodic positive selection specific to all Asteraceae taxa (Supplementary Dataset 3). Of these, several orthogroups contained genes involved in related pathways (photoperiod, autonomous, and gibberellin) that regulate flowering time (Fig. 5). These genes directly or indirectly suppress or activate *SOC1*, a floral integrator^53^, which activates floral meristem identity genes such as *LFY* (a floral meristem identity transcriptional regulator that promotes flowering transitions^54^) and *AP1* (a floral meristem identity MADS box gene^55^). Notable genes under selection in Asteraceae were: *GI*, which promotes flowering via *FT* through the photoperiod pathway^56,57^; *FVE*, which promotes flowering by suppressing *FLC* in the autonomous pathway^58,59^; *CGA1* and *GNC,* which delay flowering by suppressing *SOC1* in the gibberellin pathway^60,61^; *FD*, which interacts with *FT* to activate *SOC1*; and *AGL24,* which integrates with *SOC1* to link floral signals and promote flowering^62^ (Fig. 5, Table 1). Of these, episodic positive selection was only identified in *Acicarpha* homologs of *FD* and *CGA1*. Overall, selection on multiple genes across interconnected pathways likely fine-tuned flowering time regulation in Asteraceae.

**Fig. 5.**
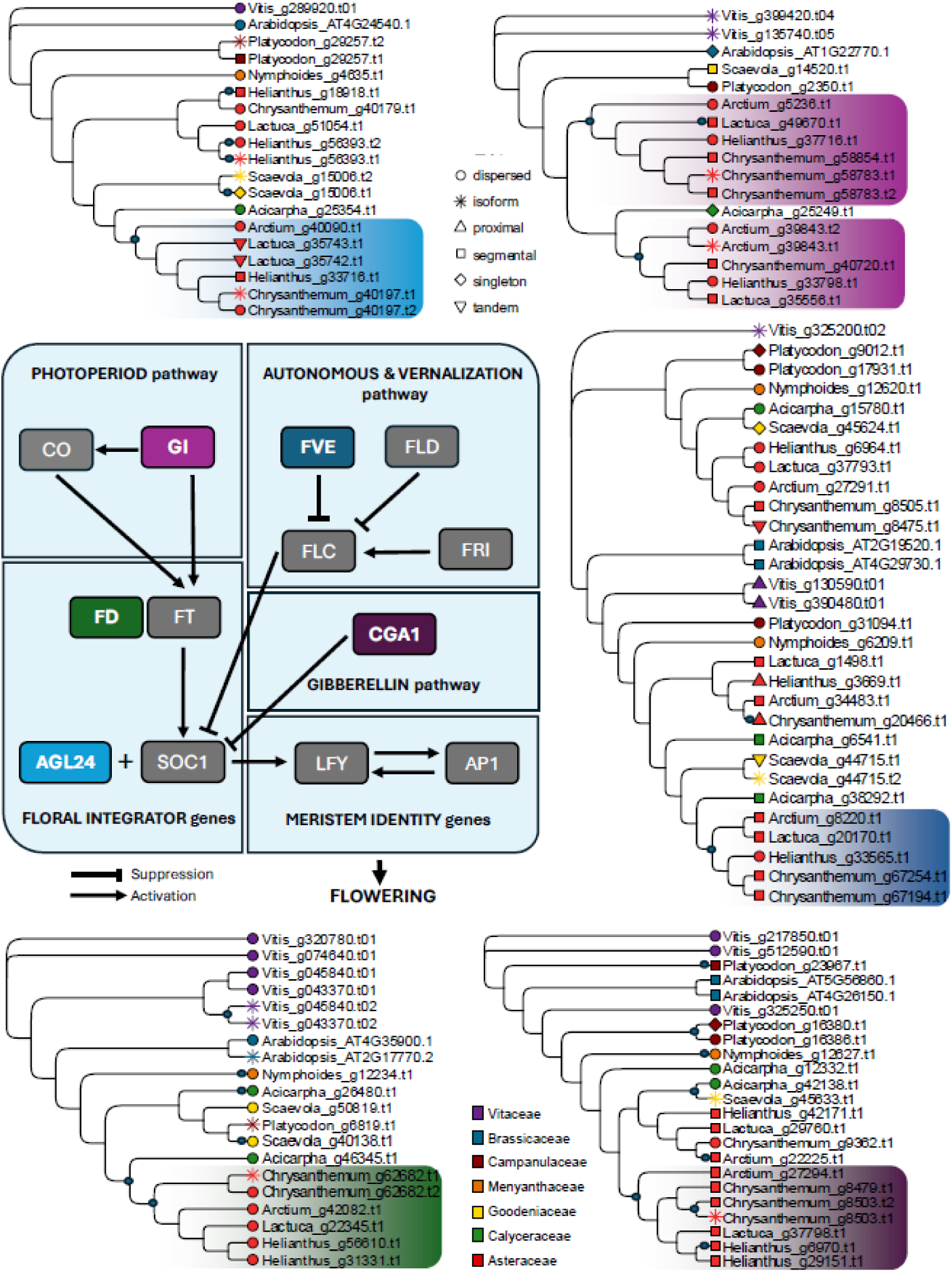
Positively selected flowering time regulatory genes. Flowering time gene network involving the photoperiod, autonomous, vernalization, and gibberellin pathways. Colored gene boxes indicate orthogroups which experienced episodic positive selection in a lineage of Asteraceae homologs. Episodes of positive selection for only Asteraceae lineages are visualized as shaded blocks in phylogenetic trees and correspond to their respective gene color. Within the phylogenetic gene trees, additional episodes of positive selection are indicated as small black circles along nodes and tips, duplication types of genes are indicated by tip shapes, and plant families are indicated by color.

## Discussion

While the role of WGDs in driving gene expansion, diversification, and selection is evident across our analyses, the number and timing of WGD events within Asterales remain difficult to resolve despite having high-quality genome assemblies. Contrasting results from different approaches (Figs. 2f, 3) highlight the inherent challenges in detecting ancient WGDs, particularly within large and diverse plant families. This complexity is compounded by processes such as biased gene retention and loss, genome rearrangements, chromosomal fission and fusion, incomplete lineage sorting, and hybridization, including allopolyploidy^63,64^. Synteny-based methods, while powerful, often yield incomplete reconstructions of ancient events, especially across distantly related taxa. Similarly, *K*_S_-based approaches face limitations: signals from ancient WGDs may be obscured by subsequent small-scale duplications and losses, attenuated with time, or confounded with speciation signals, especially in cases of allopolyploidy where increased variation in *K*_S_ values blurs inference^65^.

Our results support multiple hypotheses on WGD placement in Asterales. Triplicate syntenic gene depths echo previous studies indicating a shared paleohexaploidy event between Asteraceae and Calyceraceae^4–6^ (Fig. 3a). However, this inference was refined through sub-genome phasing and hypothesis testing to indicate that Asteraceae and Calyceraceae shared a WGD, then each experienced their own (Fig. 3b). Some *K_S_* distribution and gene family expansion results suggest that the core WGD is shared not only by Asteraceae and Calyceraceae, but also Goodeniaceae (Fig. 3c). However, this finding is unlikely given the comparative syntenic gene depths and is probably an artefact of unaccounted biological processes described above^65^.

Nonetheless, it is evident that both large-scale and small-scale duplications played a formative role in the divergence and diversification of Asterales. While WGDs account for most positively selected genes in Asteraceae, duplications originating from TE activity (i.e., tandem, proximal, and dispersed duplicates) explained most gene family expansions (Supplementary Fig. 17). A biased retention of segmental duplicates from polyploidy, particularly those supporting complex processes and protein interactions, has been observed in other systems, and the high proportion of segmental duplicates in Asteraceae and *Acicarpha* may reflect this^33,66,67^. Rather than being lost or pseudogenized, retained duplicate genes likely underwent subfunctionalization, neofunctionalization, or dosage amplification to increase adaptive fitness. These findings highlight the substantial roles tandem duplication and transposition, alongside WGD, have in generating genomic novelty^68^.

In Asteraceae, post-duplication outcomes may have enhanced ecologically beneficial processes such as secondary metabolite biosynthesis and cellular responses to environmental stimuli, which are enriched in expanded and positively selected gene families (Supplementary Fig. 20). Given Asteraceae’s rich array of secondary compounds used in defense, attraction, and environmental interaction^69^, the lineage-specific retention and expansion of genes involved in metabolite production and regulation likely reflect specialization in metabolite biosynthesis. Secondary chemistry contributes to key ecological interactions, including herbivore deterrence, pathogen resistance, and pollinator attraction, and its diversification may have enhanced environmental responsiveness and niche breadth^69^. Conversely, in *Acicarpha* the limited expansion of metabolite biosynthesis genes may have limited their geographic range through outcompetition. These patterns of gene family expansion, retention, and selection thus support the broader view that chemical diversification, driven by underlying genomic innovation, has been a major driver of Asteraceae’s evolutionary success^5^.

Advanced reproductive systems and strategies have also played a key role in Asteraceae’s evolutionary and ecological success. In a capitulum, flowering time and individual floret development is precise and highly coordinated^15,16^. Our findings suggest that selective pressures on duplicated genes and transposon activity may have fine-tuned flowering time gene regulation, conferring a key reproductive advantage in Asteraceae. Among positively selected flowering time genes, patterns of evolution varied. For example, *FD* homologs exhibited a two-step episode of positive selection: first on Asteraceae/*Acicarpha* dispersed duplicates, then again on Asteraceae and *Acicarpha* duplicates individually. The *FVE* homologs suggest that Asteraceae and *Acicarpha* segmental duplicates were derived from a shared WGD, followed by positive selection on only one Asteraceae lineage. These patterns reflect a shared history between Asteraceae and *Acicarpha*. Conversely, *GI* and *CGA1* patterns reflect an Asteraceae-specific WGD. *GI* homologs showed two episodes of diversifying selection in Asteraceae segmental duplicates, likely indicating subfunctionalization or dosage compensation (Fig. 5). Among *CGA1* homologs, however, episodic positive selection occurred only along one of the segmentally duplicated Asteraceae lineages. While likely originating from transposon activity rather than WGD, the dispersed duplicates of *Acicarpha CGA1* homologs exhibited a similar pattern of selection on one duplicated lineage. These findings underscore distinct mechanisms of functional divergence and the fate of duplicates between Asteraceae and *Acicarpha* that may have conferred unique adaptive advantages.

Furthermore, the absence of positive selection on core integrators (e.g., *SOC1*) and meristem identity genes (e.g., *LFY* and *AP1*) suggests that strong purifying selection preserves central flowering functions, while upstream regulators offer an evolutionary interface for adjusting developmental timing. This pattern suggests that adaptive evolution in Asteraceae has primarily targeted regulatory modifications within the flowering gene network, enabling flexibility without disrupting deeply conserved functions. These patterns imply lineage-specific fine-tuning of flowering time, likely in response to environmental heterogeneity. Such adjustments may have been especially advantageous during the late Paleocene-Eocene, when Asteraceae experienced climatic optimums, increasing aridity, and concurrent pollinator radiations^8,70,71^, favoring genes capable of modulating reproductive timing and capitulum organization. Because some genes in traditional plant model systems are lineage-specific, e.g., *FLC* in *Arabidopsis* (Supplementary Fig. 21), these species may not serve as broadly representative models for flowering time regulation. Instead, groups like Asteraceae and other diverse clades offer additional insights into the evolutionary dynamics of these pathways.

Genome duplication has influenced not only speciation, but also the phenotypic and ecological breadth of plants, shaping habitat use, life history strategies, competitive interactions, responses to herbivores and pathogens, and pollinator relationships^72–77^. In both Asteraceae and Calyceraceae, ancient large-and small-scale duplications left a strong genomic imprint, yet distinct evolutionary pressures acting on duplicated genes, particularly those involved in metabolite biosynthesis and flowering time regulation, drove the emergence of key ecological traits and ultimately enabled the extraordinary diversification of Asteraceae.

## Online Methods

### Taxon selection

For this study, we generated de novo chromosome-level assemblies of *Acicarpha procumbens* Less. (Calyceraceae), hereafter *Acicarpha*; *Scaevola aemula* R.Br. (Goodeniaceae), hereafter *Scaevola*; and *Platycodon grandiflorus* (Jacq.) A.DC. ‘Komachi’ (Campanulaceae), hereafter *Platycodon*, using the materials and methods described below. We included additional publicly available Asterales species for comparison: *Arctium lappa* L.^78^ (Asteraceae; NCBI: PRJNA764011), hereafter *Arctium*; *Lactuca sativa* L.^26^ (Asteraceae; NCBI: PRJNA173551), hereafter *Lactuca*; *Chrysanthemum lavandulifolium* (Fisch. ex Trautv.) Makino^79^, hereafter *Chrysanthemum* (Asteraceae; NCBI: PRJNA681093); *Helianthus annuus* L.^31,80,81^ (Asteraceae; HA412-HO v2: https://sunflowergenome.org/), hereafter *Helianthus*; and *Nymphoides indica* (L.) Kuntze^82^ (Menyanthaceae; CNGB: CNP0002767), hereafter *Nymphoides*. Additionally, for broader evolutionary and functional comparisons, we included the well-studied genomes: *Vitis vinifera* L. cv. Cabernet Sauvignon cl. 08 v 1.1^83^ (Vitaceae), hereafter *Vitis,* and *Arabidopsis thaliana* (L.) Heynh. (Brassicaceae; TAIR10; https://v2.arabidopsis.org/), hereafter *Arabidopsis* (Supplementary Table 6).

We collected seeds of *Acicarpha* in Uruguay, purchased seeds of *Platycodon* through Park Seed (https://parkseed.com/), and acquired cuttings of *Scaevola* through North Carolina Farms Inc. (https://www.ncfarmsinc.com/). We grew all plants in greenhouses at the University of Memphis (Memphis, Tennessee, USA) and Auburn University (Auburn, Alabama, USA). A single individual of *Platycodon* was taken through two successive rounds of inbreeding prior to use. *Acicarpha* and *Scaevola* individuals were not inbred before sequencing. Vouchers for each specimen of *Acicarpha*, *Scaevola*, and *Platycodon* are available at the University of Memphis herbarium (MEM) and Auburn University Herbarium (AUA) (see Supplementary Table 7 for details).

### Sequencing

For *Acicarpha*, we generated short- and long-read DNA, Omni-C, and RNA sequencing data. For short reads, genomic DNA was extracted from ∼25 mg of fresh leaf tissue from a single individual ground with a Bead Mill 24 Homogenizer (ThermoFisher Scientific, Atlanta, Georgia, USA) using the E.Z.N.A. SP Plant DNA Kit (Atlanta, Georgia, USA). High molecular weight (HMW) DNA for long-read sequencing was isolated from 2.5–3 g of flash-frozen young leaf tissue ground in liquid nitrogen with an autoclaved mortar and pestle using the TaKaRa NucleoBond HMW DNA kit (Düren, Germany). RNA was extracted from young leaf (YL), shoot apical meristem (SAM), and mature leaf (ML) tissues using the E.Z.N.A. Plant RNA kit (Omega Bio-tek, cat #R6827-01) and treated with RNase-free DNase (Omega Bio-tek, cat #E1091). Flash-frozen tissue was sent to the HudsonAlpha Institute for Biotechnology (Huntsville, Alabama, USA) for Omni-C library preparation using the Dovetail Omni-C kit (Dovetail Genomics, Inc.). The gDNA, Omni-C libraries, and the RNA were sequenced using an Illumina NovaSeq 6000 with PE150 chemistry, the HMW DNA was sequenced on a PacBio Sequel II at HudsonAlpha Institute for Biotechnology.

For *Scaevola*, short- and long-read DNA, Hi-C, and RNA data were generated using equivalent tissue amounts and extraction kits as for *Acicarpha*. Hi-C libraries were prepared with the Proximo Hi-C Kit-Plant (Phase Genomics, cat #KT3040) and sequenced at the University of North Carolina’s High Throughput Sequencing Facility (UNC HTSF) on a NovaSeq 6000 (PE-50 and PE-100). The gDNA and HMW DNA were sequenced on NovaSeq PE150 and PacBio Sequel II sequencers, respectively, at HudsonAlpha Institute for Biotechnology.

For *Platycodon*, we generated short- and long-read DNA, Hi-C, and RNA sequencing data. Genomic DNA was extracted from ∼100 mg of homogenized tissue using a modified CTAB protocol. Short-read libraries were prepared at UNC’s HTSF with the KAPA Hyper DNA kit and sequenced at BGI-Genomics on the DNBseq platform (PE100). Total RNA from YL and SAM tissues of five individuals was extracted using the E.Z.N.A. Plant RNA kit (Omega Bio-tek, #R6827-01), treated with RNase-free DNase (#E1091), and used for library preparation (KAPA stranded mRNA-seq kit, UNC HTSF). Hi-C libraries were prepared with the Proximo Hi-C Kit-Plant (Phase Genomics, #KT3040) following the v4.0 protocol (October 2020) and sequenced at UNC HTSF on a NovaSeq 6000 (PE100). For long reads, HMW DNA was extracted using the ONT “High molecular weight gDNA extraction from plant leaves” (CTAB–Genomic-tip) protocol, purified with Amicon Ultra filters (#UFC510008), and size-selected using the Circulomics Short Read Eliminator Kit (#SS-100-101-01, v2). ONT libraries were prepared with the Ligation Sequencing Kit (#SQK-LSK109) following previously described modifications¹²⁴ and sequenced on a MinION Mk1B (MIN-101B) using FLO-MIN106D flow cells (R9.4.1 pore) controlled by MinKNOW v3.6.5. Two flow cells were used; each washed three times with the Flow Cell Wash Kit (#EXP-WSH003) and reloaded with the same library.

We checked raw long-read and short-read sequence data quality using FastQC^84^ v. 0.11.9, a program designed to highlight potential issues with sequence data. Additionally, we checked raw long-read data quality using PAUVRE^85^ v. 0.2.3, which plots the length and quality distribution of raw Oxford Nanopore (ONT) or PacBio reads (Supplementary Fig. 19). We then trimmed raw reads to filter out inadequate reads and remove adapters using fastp^86^ v. 0.23.2.

### Genome assembly and scaffolding

We assembled long-read data for *Scaevola* and *Acicarpha* using HifiASM^87^ v. 0.19.5 and v. 0.18.5, respectively, with Hi-C data for *Scaevola* and Omni-C data for *Acicarpha* integrated into the assemblies, resulting in two haplotypes per species. We base-called the *Platycodon* MinION data using the python package guppy v. 3.1.6 (ONT) with model R9.4.1_450bps in high-accuracy mode. We then assembled the genome with a hybrid approach utilizing ONT long-read and DNBseq (BGI) short-read genomic DNA sequencing as inputs. For the hybrid assembly, we used MaSuRCA^88^ v. 3.4.2, and polished with short-read data using bowtie2^89^ v. 2.1.4 and pilon^90^ v. 1.22.

We aligned Omni-C and Hi-C reads to the assembly using BWA^91,92^ v. 0.7.17 and used Samtools^93^ v1.11 to convert the output sam file into a bam file, sorted the bam file, and created an index for the assembly fasta file to prepare for scaffolding. We conducted scaffolding, followed by manual curation of the *Scaevola*, *Acicarpha*, and *Platycodon* assemblies, using Yet another Hi-C Scaffolding tool (YaHS)^94^ v1.1 for *Acicarpha* and v. 1.2a.2 for *Platycodon* and *Scaevola*, with the Omni-C data for *Acicarpha*, Hi-C data for the *Scaevola*, and *Platycodon* used as input. We prepared the resulting assemblies for Juicebox^95^ (v. 2.17.00 for *Acicarpha* and v. 2.15.00 for *Scaevola* and *Platycodon*) visualization using the Juicer Pre tools v1.1 included in YaHS before inputting them into the Juicebox GUI for manual scaffolding and correction to produce pseudochromosomes. We assessed genome assembly quality and completeness using Benchmarking Universal Single-Copy Orthologs (BUSCO)^96^ v. 5.4.7 with embryophyta from OrthoDB (https://www.orthodb.org) as the reference lineage. We then generated assembly statistics of the finalized assemblies using the publicly available python script (https://github.com/lexnederbragt/sequencetools/blob/master/assemblathon_stats.py).

### Genome annotation

We screened the *Acicarpha*, *Scaevola*, and *Platycodon* genomes for bacterial contaminants using StandardDB from Kraken2^97^ v. 2.0.8 and removed all contigs with positive hits. Using the resulting files as input, we then used the Extensive de-novo TE Annotator (EDTA)^98^ pipeline v. 1.9.3 to run RepeatModeler2^99^ and identify remaining TEs utilizing the default parameters, except --overwrite, --sensitive, --anno, and --evaluate were set to ‘1’. Using the decontaminated genome file and the resulting TE library file from EDTA, we then ran RepeatMasker^100^ v. 4.1.5 to soft mask the genomes. We annotated the soft masked genomes with RNA sequences using Braker3^101^ v. 3.0.3 and labelled functional gene annotations using InterProScan^102^ v. 5.66-98.0. We then used OMArk v. 0.3.0 (https://omark.omabrowser.org/, accessed June 3, 2024) to assess the proteome quality of *Acicarpha*, *Scaevola*, and *Platycodon* with Asterids (NCBI ID: 71274) as the ancestral clade.

To ensure consistency between annotations and comparative genomic analyses, we re-annotated five publicly available genomes from *Nymphoides, Arctium, Chrysanthemum, Helianthus, Lactuca* following the steps detailed above. We assessed annotation quality across all eight taxa using the gff3_stats python script (https://github.com/HuffordLab/GenomeQC/blob/master/scripts/gff3_stats.py) from GenomeQC^103^.

### Plastid and mitochondrial genome assembly

We extracted chloroplast sequences from all trimmed paired end short read data using the GetOrganelle toolkit^104^ following the recommended parameters for embryophyte plant plastomes but increasing the number of rounds and the disentangle time limit (-R 75 -- disentangle-time-limit 7200). For long read data, PacBio and ONT minion, we used ptGAUL to extract chloroplast sequences^105^. We then annotated complete circular sequences using GeSeq^106^ and assessed collinearity between plastomes using the progressive alignment in Mauve^107^. In addition, we assembled mitochondrial genomes of *Platycodon, Scaevola,* and *Acicarpha* using OatK v. 1.0 (https://github.com/c-zhou/oatk).

### Genome structure

We characterized genomic sequences’ size, heterozygosity, and repetitiveness by k-mer frequency analyses (k = 21) using Jellyfish^108^ and GenomeScope^109^ 2.0. We assessed heterozygous k-mer pairs, obtained with KMC3^110^, using Smudgeplot^109^ to estimate ploidy levels and infer genomic complexity in terms of sub-genome composition. We identified lower (L) and upper (U) end cut-off values, below and above which all k-mers were discarded as errors, using k-mer coverage output from Genomescope 2.0, as recommended in the Smudgeplot documentation (https://github.com/KamilSJaron/smudgeplot), where L = (kcov/2)-5.

### Gene duplications

We investigated gene duplications of all eight target Asterales genomes, plus genomes from *Vitis* and *Arabidopsis*, by assessing homology and synteny to identify duplicated genes and genomic regions, then by analyzing *K*_S_ values to estimate the relative age of duplication events. To quantify and classify types of gene duplications within each genome, we used the *“*duplicate gene classifier” program in MCScanX^33^. Through this program, duplicated genes were classified as: segmental, proximal, tandem, or dispersed, as defined by ^33^.

We utilized OrthoFinder v. 2.5.5^111^ to identify conserved orthologs and gene duplications across the ten genomes, using the -M msa parameter to generate a multiple species alignment. We then identified the best evolutionary model and performed a maximum likelihood phylogenetic analysis on the alignment using IQTree^112^ v. 2.1.3 with 1,000 bootstraps to generate a species tree. We used the orthologous gene groups identified by OrthoFinder for downstream analyses to investigate evolutionary processes, however, we filtered the groups to exclude multiple isoforms of the same gene by keeping only the longest isoform.

### Syntenic gene depths

To investigate putative paleopolyploidy, we assessed synteny among the eight Asterales genomes. We used the JCVI^113^v. 1.3.8 python function “compara catalog ortholog” with the default c-score cutoff of 0.7 to identify syntenic regions, then generated genome-wide synteny plots of the full eight genomes following phylogenomic placement. We evaluated pairwise syntenic gene depth between all genomes using the “compara synteny depth” option. We then used these inferred values as input for subsequent analyses to phase subgenomes and further elucidate polyploid origins (detailed in “Phasing subgenomes” below).

### Phasing subgenomes

We implemented the two most prominent software tools to identify and separate subgenomes, SubPhaser^114^ and Whole-Genome Duplication Integrated (WGDI^115^). However, given that the tested taxa have not experienced recent polyploidization events and are currently recognized as diploids, SubPhaser could not be used with our data as it is unable to accurately phase genomes that experienced paleopolyploidy events. Instead, we used WGDI v. 0.6.5 to further investigate polyploidization events across Asterales using *Vitis* as a reference. Steps to investigate polyploidy followed the subgenome aware analyses available on GitHub (https://github.com/SunPengChuan/wgdi-example/blob/main/phase_subgenomes.md). Multiple chromosomes of *Vitis* were tested as a reference for phasing, though only the chromosome with the largest gene retention and collinearity, chromosome 11, was used for further analyses. Age estimates from *K*_S_ values (e.g., Supplementary Fig. 23) were used to help separate subgenomes if they were one the same chromosome or were not distinct. We determined phylogenomic relationships among the subgenomes using WGDI’s ‘-at’ parameter to generate gene trees with IQTree and then ASTRAL-III^116^ v. 5.7.8 to generate a final subgenome coalescent tree. We also assessed gene fractionation within each subgenome to determine the evolutionary age of each subgenome using the ‘r’ parameter of WGDI. We then reanalyzed the resulting subgenomes within Asteraceae, Calyceraceae, and Goodeniaceae in JCVI to investigate synteny across these phased regions using methods detailed above.

### Accelerated gene evolution in *Acicarpha* and Asteraceae

To identify major targets of accelerated evolution within conserved genomic regions, we used Cactus^117–119^ v2.5.1, to generate a reference-free alignment of the Asterales subgenomes that are syntenic to chromosome 11 of *Vitis*. Using the inferred phylogenomic relationship between subgenomes as the guide tree to inform hierarchical alignment, we performed three separate tests for accelerated evolution at different positions in the subgenome tree: across all Asteraceae and *Acicarpha* subgenomes (clade 1; Supplementary Fig. 11), in the *Acicarpha*-specific duplication event (clade 2; Supplementary Fig. 11), and in *Acicarpha* within the Subgenome 1 clade (clade 3; Supplementary Fig. 11).

Specifically, we created a neutral model of sequence evolution based on ancestral repetitive sequences^120^. Repetitive elements were identified within the most ancestral sequence in the subgenome alignment using RepeatModeler and RepeatMasker^100^. A random sample of 100,000 sites from these ancestral repeats was then selected to generate a multiple alignment file (MAF) of repetitive sites across all subgenomes. This MAF was used to construct the neutral model with PHAST phyloFit^121,122^, using default settings and the same phylogeny used to generate the subgenome alignment (Supplementary Fig. 11).

With this neutral model, we identified conserved regions within the Asterales subgenomes using PHAST phastCons^121,122^, with *Vitis* as the reference. This was performed separately for each of the three clade-specific analyses, with conserved regions detected in the outgroup subgenomes. For each analysis, we obtained a list of the most-conserved elements and converted their log-odds scores to length-normalized log-odds ratios. Elements with normalized log-odds ratios below the 30th percentile were excluded from further analyses^123^. This filtering step produced three distinct lists of conserved regions in outgroup subgenomes, allowing us to assess accelerated evolution within our clades of interest. We then tested for GC-biased gene conversion (gBGC) within our ingroups using PHAST phastBias^121^ and considered any region with a score >0.5 to be impacted by gBGC. We filtered our lists of conserved regions to remove any elements that were impacted by gBGC.

We used the likelihood ratio test of PHAST phyloP^121,124^ to test the conserved elements for accelerated evolution within our clades of interest. We corrected for multiple hypothesis testing using Storey’s correction method^125^ (FDR = 0.05). This produced a separate list of accelerated regions (ARs) for each of our clade-specific analyses. To assess the biological impacts of the accelerated evolution detected in each of our analyses, we identified the genes most likely to be impacted by each AR. We associated an AR with a gene if it overlapped an exon. Exon identities and locations were obtained from annotations of the *Vitis* genome^83^.

ARs can be distributed unevenly across the genome, often clustering into regions of high density, known as AR hotspots. To detect these hotspots, we employed the permutation method of Cahill et al.^126^. Specifically, we used a sliding window approach, counting ARs in 100 kb windows along *Vitis* chromosome 11, with windows offset by 10 kb steps. We conducted 100 permutations, each time randomly designating N conserved elements as accelerated, where N corresponds to the actual number of ARs on the chromosome. For each permutation, we repeated the sliding window approach to determine the maximum simulated AR density in a 100 kb window. Regions were designated as AR hotspots if their true AR density exceeded the 95th percentile of maximum AR densities observed across all permutations. This approach prevents regions with high conserved element density from being falsely identified as hotspots. We identified separate hotspots for each of our clade-specific analyses.

### *K*_S_ distributions

To investigate the influence of major genome duplication events and positive selection on Asterales genomes, we assessed the non-synonymous (*K*_a_) and synonymous (*K*_S_) substitution rates between gene pairs. Peaks in *K*_S_ value distributions, indicating the presence of duplicated genes, were used to infer histories of WGD. For homologous gene pairs, we estimated *K*_S_ values using an established pipeline^127^ in which orthologs were aligned using MUSCLE^128^, then converted into nucleotides using PAL2NAL^129^, then *K*_S_ values were calculated between gene pairs using the F3X4 model in codeml within PAML^130^. We plotted *K*_S_ values of paralogs and pairwise orthologous gene pairs in R^131,132^ with kernel density estimations using geom_density() from ggplot2^133^. Additionally, we used output from these analyses to identify signatures of positive selection (*K*_a_/*K*_S_ > 1), neutral selection (*K*_a_/*K*_S_ ≈ 1), and purifying selection (K_a_/*K*_S_ < 1) of paralogous gene pairs. To more accurately compare estimations of divergence derived from orthologous *K*_S_ distributions with estimations of WGD derived from paralogous *K*_S_ distributions, we then re-scaled ortholog distributions to the paralog scale of each focal species using *ksrates*^134^. For these analyses, *Platycodon* was used as the outgroup, therefore, WGD events were not assessed for this lineage.

### WGD hypothesis testing

Directly following previously published methods and scripts^135,136^ (https://gitlab.com/barker-lab/ajb_special_issue_2024.git), we tested predicted WGD events, hypothesized by previous literature and identified by *ksrates*, using WHALE^35^. WHALE is a tool designed to test WGD hypotheses by analyzing the relative support for different evolutionary scenarios. It uses gene tree topologies and considers the expected patterns of gene retention and loss following WGD events. By applying a likelihood-based framework, WHALE evaluates whether observed gene tree discordance is best explained by WGD or alternative models, making it a valuable approach for investigating genome evolution. Given results from *ksrates* and primary literature, we tested nine total WGD events split into three separate analyses: one based solely off findings from previous studies, the second analysis used only the results of *ksrates*, and the third analysis combined results of the first two analyses (Supplementary Fig. 15).

### Evolution of gene family size

To investigate the evolution of gene family sizes in Asterales, we reconstructed ancestral states of the orthologous groups identified by OrthoFinder by employing the fast algorithm^137^ through the ‘ace’ function in the *phytools*^138^ R package. From the reconstructed ancestral states, we applied targeted filters to classify orthogroups as: 1) specific to *Acicarpha*, 2) lost in *Acicarpha*, 3) specific to Asteraceae (evolved in the ancestral branch of Asteraceae and absent from other lineages), 4) lost in the last common ancestor of Asteraceae, 5) specific to both Asteraceae and *Acicarpha*, and 6) lost in the last common ancestor of both Asteraceae and *Acicarpha*. Filtering was based on a Bayes factor threshold of 10, indicating that the likelihood of an orthogroup being present or absent at a given node was ten times higher than the alternative scenario.

Next, we used CAFE^139^ v. 5.0 to further investigate the dynamics of orthogroup expansion and contraction. Since CAFE requires an ultrametric tree, we used TreePL^140,141^ to calibrate the phylogeny reconstructed by OrthoFinder. We selected four calibration points based on previous work^8^, constraining the minimum ages of ancestral nodes for the following groups: Asteraceae and Calyceraceae (*Tubulifloridites lillei* Type A^142^; 72.1 Mya), all Asteraceae included in the study (*Raiguenrayun cura*^143^; 47.5 Mya), *Helianthus* and *Lactuca* (*Cichorium intybus* type pollen^144^; 22 Mya), and *Helianthus* and *Chrysanthemum* (*Artemisia* type pollen^145^; 31 Mya). The root age was constrained between 121 and 126 Mya using a secondary calibration point for Pentapetalae^146^ (Supplementary Fig. 24). The optimal smoothing parameter was determined through cross-validation, testing 20 values across a range of magnitudes (from 1e-10 to 1e+09) to ensure the best fit.

The tested CAFE models varied in the number of gamma parameters (ranging from 1 to 3), reflecting assumed differences in expansion and contraction rates across orthogroups. Orthogroups identified by the best-fitting model as significantly expanded or contracted (Viterbi ≤ 0.05) in Asteraceae, Acicarpha, and both Asteraceae and Acicarpha were retained for downstream analyses, in addition to those identified as lost or specific to these groups. Genes from significantly expanded orthogroups were then blasted to the *Arabidopsis thaliana* database, TAIR10, using BLASTP^147^ to determine closest gene similarity and trees were then visualized in R.

### Positive selection

We further investigated positive selection across Asterales genomes, focusing on Asteraceae, using site-specific and branch-specific methods. For all taxa, we assessed orthology and subsequent positive selection of single-copy orthologous groups using site-specific methods within the AlexandrusPS pipeline^148^. For the orthogroups which were found to exhibit positive selection for genes pertaining to all four Asteraceae taxa and no others, we analyzed further by using branch-specific methods to determine if the found signatures of positive selection originated from taxon-specific or family-specific lineages. Additionally, we assessed a subset of orthogroups relating to floral development for lineage-specific positive selection using branch-specific methods. These floral-related orthogroups were chosen based on GO annotations for *Arabidopsis thaliana* genome (TAIR10: available at https://v2.arabidopsis.org/) to include 18 terms related to meristem maintenance and organization (GO:0010582, GO:0009933, GO:0009934, GO:0048507, GO:0010074, GO:0019827), flower development (GO:0009908, GO:0010076, GO:0009910, GO:009911, GO:0048438, GO:0010093, GO:2000032), vegetative shoot system/inflorescence development (GO:0048367, GO:0010077, GO:0010492, GO:0090506,), and vegetative-reproductive phase transition (GO:0048510). For both subsets of orthogroups, we generated codon alignments from coding sequences by adding gaps for frameshifts and replacing stop codons with gaps using MACSE^149^. Then we used the aBSREL (adaptive Branch-Site Random Effects Likelihood) function in HyPhy^150^ to test if positive selection has occurred on a proportion of branches. We inspected results using HyPhy Vision (http://vision.hyphy.org/aBSREL), then visualized resulting gene trees in R using Ape 5.0^151^ and ggtree^152^.

### GO enrichment

For inferences on the functional role of genes identified in these analyses, we used the basic local alignment search tool (BLAST^147^) to compare target proteins to a TAIR10 protein database and identify matches based on an e-value threshold of 1e-10. For groups of gene groups identified to be expanded or contracted or under positive selection, we performed gene ontology (GO) enrichment analyses to summarize the significant GO terms for biological processes^153^. We used the clusterProfiler^154^ package in R, with annotations based on BLASTP best-hit matches to the TAIR10 database. To account for multiple testing, p-values were adjusted using the Benjamini–Hochberg false discovery rate (FDR) correction. Enrichment was considered significant at an adjusted p-value (P-adjust) threshold of < 0.01.

## Acknowledgements

We thank Jane Grimwood (HudsonAlpha), Eric Lyons (University of California, Berkley) for assistance and consultation on CoGe, and Eric Spangler and Kristian Skjervold of the University of Memphis High-Performance Computing (HPC) cluster.

## Author contributions

ERMP, PE, JMB, DSJ, and JRM conceived the study. ERMP, PE, BRC, JB, MDP, and ZM performed data curation, formal analyses, and developed methodologies and software. ERMP, PE, JB, and JRM led the investigation, data validation, visualization, and drafting of the manuscript. ACW contributed to data visualization. MDP, SD, and ZM provided additional analytical and software support. AH, MB, ZN, JMB, DSJ, and JRM contributed to manuscript review and editing. MB and ZN provided resources, and ZN, JMB, DSJ, and JRM acquired funding and supervised the project. All authors read and approved of the final manuscript.

## Data availability

Raw data and genome assembly data for *Acicarpha*, *Scaevola*, and *Platycodon* are uploaded to NCBI BioProject PRJNA1253301. Accession numbers of these newly developed and publicly available data can be found in Supplementary Table 6. Hardmasked, annotated genome assemblies for *Acicarpha, Scaevola,* and *Platycodon,* as well as the re-annotated assemblies for *Helianthus, Chrysanthemum, Lactuca, Arctium,* and *Nymphoides* were uploaded and made publicly accessible on the Comparative Genomics Platform^152^ (id67898, id67904, id67903, id67900, id67899, id67901, id67867, id67902, respectively).

## References

1. Vamosi, J. C., Magallón, S., Mayrose, I., Otto, S. P. & Sauquet, H. Macroevolutionary Patterns of Flowering Plant Speciation and Extinction. Annu Rev Plant Biol 69, 685–706 (2018).

2. Benítez-Villaseñor, A. et al. The use of Anchored Hybrid Enrichment data to resolve higher-level phylogenetic relationships: A proof-of-concept applied to Asterales (Eudicotyledoneae; Angiosperms). Mol Phylogenet Evol 181, 107714 (2023).

3. Li, H.-T. et al. Origin of angiosperms and the puzzle of the Jurassic gap. Nat Plants 5, 461–470 (2019).

4. Barker, M. S. et al. Most Compositae (Asteraceae) are descendants of a paleohexaploid and all share a paleotetraploid ancestor with the Calyceraceae. Am J Bot 103, 1203– 1211 (2016).

5. Huang, C.-H. et al. Multiple Polyploidization Events across Asteraceae with Two Nested Events in the Early History Revealed by Nuclear Phylogenomics. Mol Biol Evol 33, 2820– 2835 (2016).

6. Zhang, C. et al. Phylotranscriptomic insights into Asteraceae diversity, polyploidy, and morphological innovation. J Integr Plant Biol 63, 1273–1293 (2021).

7. Palazzesi, L. et al. Asteraceae as a model system for evolutionary studies: from fossils to genomes. Bot. J. Linn. Soc. 200, 143–164 (2022).

8. Mandel, J. R. et al. A fully resolved backbone phylogeny reveals numerous dispersals and explosive diversifications throughout the history of Asteraceae. Proc Natl Acad Sci U S A 116, 14083–14088 (2019).

9. Panero, J. L. & Crozier, B. S. Macroevolutionary dynamics in the early diversification of Asteraceae. Mol Phylogenet Evol 99, 116–132 (2016).

10. Susanna, A. et al. The classification of the Compositae: A tribute to Vicki Ann Funk (1947–2019). Taxon 69, 807–814 (2020).

11. Diazgranados, M. et al. World Checklist of Useful Plant Species. KNB Data Repository https://knb.ecoinformatics.org/view/doi:10.5063/F1CV4G34 (2020).

12. Lundberg, J. G. Asteraceae and relationships within Asterales. in (2009).

13. Brignone, N. F., Mazet, N., Pozner, R. & Denham, S. S. Calyceraceae: Unexpected diversification pattern in the Southern Andes. Perspect Plant Ecol Evol Syst 60, 125744 (2023).

14. Denham, S. S., Zavala-Gallo, L., Johnson, L. A. & Pozner, R. E. Insights into the phylogeny and evolutionary history of Calyceraceae. Taxon 65, 1328–1344 (2016).

15. Zhang, T. & Elomaa, P. Development and evolution of the Asteraceae capitulum. New Phytol 242, 33–48 (2024).

16. Gurung, V., Muñoz-Gómez, S. & Jones, D. S. Putting heads together: Developmental genetics of the Asteraceae capitulum. Curr Opin Plant Biol 81, 102589 (2024).

17. Pozner, R., Zanotti, C. & Johnson, L. A. Evolutionary origin of the Asteraceae capitulum: Insights from Calyceraceae. Am J Bot 99, 1–13 (2012).

18. Teeri, T. H. et al. Reproductive meristem fates in *Gerbera*. J Exp Bot 57, 3445–3455 (2006).

19. Flagel, L. E. & Wendel, J. F. Gene duplication and evolutionary novelty in plants. New Phytol 183, 557–564 (2009).

20. Doyle, J. J. Polyploidy in Legumes. in Polyploidy and Genome Evolution 147–180 (Springer Berlin Heidelberg, Berlin, Heidelberg, 2012). doi:10.1007/978-3-642-31442-1_9.

21. Eric Schranz, M., Mohammadin, S. & Edger, P. P. Ancient whole genome duplications, novelty and diversification: the WGD Radiation Lag-Time Model. Curr Opin Plant Biol 15, 147–153 (2012).

22. Soltis, D. E. et al. Polyploidy and angiosperm diversification. Am J Bot 96, 336–348 (2009).

23. Shen, F. et al. Comparative genomics reveals a unique nitrogen-carbon balance system in Asteraceae. Nat Commun 14, (2023).

24. Kong, X. et al. Two-step model of paleohexaploidy, ancestral genome reshuffling and plasticity of heat shock response in Asteraceae. Hortic Res 10, (2023).

25. Barker, M. S. et al. Multiple Paleopolyploidizations during the Evolution of the Compositae Reveal Parallel Patterns of Duplicate Gene Retention after Millions of Years. Mol Biol Evol 25, 2445–2455 (2008).

26. Reyes-Chin-Wo, S. et al. Genome assembly with in vitro proximity ligation data and whole-genome triplication in lettuce. Nat Commun 8, 14953 (2017).

27. Oliver, K. R., McComb, J. A. & Greene, W. K. Transposable Elements: Powerful Contributors to Angiosperm Evolution and Diversity. Genome Biol Evol 5, 1886–1901 (2013).

28. Fedoroff, N. Transposons and genome evolution in plants. Proc Natl Acad Sci USA 97, 7002–7007 (2000).

29. Lippman, Z. et al. Role of transposable elements in heterochromatin and epigenetic control. Nature 430, 471–476 (2004).

30. Staton, S. E. & Burke, J. M. Evolutionary transitions in the Asteraceae coincide with marked shifts in transposable element abundance. BMC Genomics 16, 623 (2015).

31. Badouin, H. et al. The sunflower genome provides insights into oil metabolism, flowering and Asterid evolution. Nature 546, 148–152 (2017).

32. Schwacke, R., Bolger, M. E. & Usadel, B. PubPlant: a continuously updated online resource for sequenced and published plant genomes. Preprint at 10.1101/2025.03.12.642823 (2025).

33. Wang, Y. et al. MCScanX: a toolkit for detection and evolutionary analysis of gene synteny and collinearity. Nucleic Acids Res 40, e49–e49 (2012).

34. Jiao, Y. et al. A genome triplication associated with early diversification of the core eudicots. Genome Biol 13, R3 (2012).

35. Zwaenepoel, A. & Van de Peer, Y. Inference of Ancient Whole-Genome Duplications and the Evolution of Gene Duplication and Loss Rates. Mol Biol Evol 36, 1384–1404 (2019).

36. Zhang, C. et al. Asterid Phylogenomics/Phylotranscriptomics Uncover Morphological Evolutionary Histories and Support Phylogenetic Placement for Numerous Whole-Genome Duplications. Mol Biol Evol 37, 3188–3210 (2020).

37. Jin, Y. et al. Functional divergence of *GLP* genes between *G. barbadense* and *G. hirsutum* in response to *Verticillium dahliae* infection. Genomics 114, 110470 (2022).

38. Wang, Z. et al. Comparative genomics analysis of WAK/WAKL family in Rosaceae identify candidate WAKs involved in the resistance to *Botrytis cinerea*. BMC Genomics 24, 337 (2023).

39. Dar, A. A., Choudhury, A. R., Kancharla, P. K. & Arumugam, N. The *FAD2* Gene in Plants: Occurrence, Regulation, and Role. Front Plant Sci 8, (2017).

40. Li, Y. & Wei, K. Comparative functional genomics analysis of cytochrome P450 gene superfamily in wheat and maize. BMC Plant Biol 20, 93 (2020).

41. Quezada, E.-H. et al. Cysteine-Rich Receptor-Like Kinase Gene Family Identification in the Phaseolus Genome and Comparative Analysis of Their Expression Profiles Specific to Mycorrhizal and Rhizobial Symbiosis. Genes (Basel) 10, 59 (2019).

42. Feng, T., et al. *FAD2* Gene Radiation and Positive Selection Contributed to Polyacetylene Metabolism Evolution in Campanulids. Plant Physiol 181, 714–728 (2019).

43. Dong, Y. et al. The *Carthamus tinctorius* L. genome sequence provides insights into synthesis of unsaturated fatty acids. BMC Genomics 25, 510 (2024).

44. Pandian, B. A., Sathishraj, R., Djanaguiraman, M., Prasad, P. V. V. & Jugulam, M. Role of Cytochrome P450 Enzymes in Plant Stress Response. Antioxidants 9, 454 (2020).

45. Chakraborty, P., Biswas, A., Dey, S., Bhattacharjee, T. & Chakrabarty, S. Cytochrome P450 Gene Families: Role in Plant Secondary Metabolites Production and Plant Defense. J Xenobiot 13, 402–423 (2023).

46. Chen, Y., Klinkhamer, P. G. L., Memelink, J. & Vrieling, K. Diversity and evolution of cytochrome P450s of *Jacobaea vulgaris* and *Jacobaea aquatica*. BMC Plant Biol 20, 342 (2020).

47. Alshareef, N. O. et al. Root remodeling mechanisms and salt tolerance trade-offs: The roles of HKT1, TMAC2, and TIP2;2 in Arabidopsis. PLoS Genet 21, e1011713 (2025).

48. Mukhtar, M. S., Deslandes, L., Auriac, M., Marco, Y. & Somssich, I. E. The Arabidopsis transcription factor WRKY27 influences wilt disease symptom development caused by *Ralstonia solanacearum*. The Plant Journal 56, 935–947 (2008).

49. Han, X., et al. *Arabidopsis* Transcription Factor TCP5 Controls Plant Thermomorphogenesis by Positively Regulating PIF4 Activity. iScience 15, 611–622 (2019).

50. van Es, S. W. et al. Novel functions of the Arabidopsis transcription factor TCP5 in petal development and ethylene biosynthesis. The Plant Journal 94, 867–879 (2018).

51. Yu, H. et al. TCP5 controls leaf margin development by regulating KNOX and BEL-like transcription factors in Arabidopsis. J Exp Bot 72, 1809–1821 (2021).

52. Chapman, M. A., Leebens-Mack, J. H. & Burke, J. M. Positive Selection and Expression Divergence Following Gene Duplication in the Sunflower *CYCLOIDEA* Gene Family. Mol Biol Evol 25, 1260–1273 (2008).

53. Lee, J. & Lee, I. Regulation and function of SOC1, a flowering pathway integrator. J Exp Bot 61, 2247–2254 (2010).

54. Blackman, B. K. et al. Contributions of Flowering Time Genes to Sunflower Domestication and Improvement. Genetics 187, 271–287 (2011).

55. Li, D. et al. Arabidopsis Class II TCP Transcription Factors Integrate with the FT–FD Module to Control Flowering. Plant Physiol 181, 97–111 (2019).

56. Sawa, M. & Kay, S. A. GIGANTEA directly activates *Flowering Locus T* in *Arabidopsis thaliana*. Proc Natl Acad Sci USA 108, 11698–11703 (2011).

57. Sawa, M., Nusinow, D. A., Kay, S. A. & Imaizumi, T. FKF1 and GIGANTEA Complex Formation Is Required for Day-Length Measurement in *Arabidopsis*. Science (1979) 318, 261–265 (2007).

58. Xie, H. et al. DNA Methylation of the Autonomous Pathway Is Associated with Flowering Time Variations in *Arabidopsis thaliana*. Int J Mol Sci 25, 7478 (2024).

59. Simpson, G. G. The autonomous pathway: epigenetic and post-transcriptional gene regulation in the control of Arabidopsis flowering time. Curr Opin Plant Biol 7, 570–574 (2004).

60. Richter, R., Bastakis, E. & Schwechheimer, C. Cross-Repressive Interactions between SOC1 and the GATAs GNC and GNL/CGA1 in the Control of Greening, Cold Tolerance, and Flowering Time in Arabidopsis. Plant Physiol 162, 1992–2004 (2013).

61. Bellucci, E. et al. Selection and adaptive introgression guided the complex evolutionary history of the European common bean. Nat Commun 14, 1908 (2023).

62. Huang, N., Tien, H. & Yu, T. Arabidopsis leaf-expressed *AGAMOUS-LIKE 24* mRNA systemically specifies floral meristem differentiation. New Phytol 241, 504–515 (2024).

63. Van de Peer, Y., Mizrachi, E. & Marchal, K. The evolutionary significance of polyploidy. Nat Rev Genet 18, 411–424 (2017).

64. Soltis, P. S., Marchant, D. B., Van de Peer, Y. & Soltis, D. E. Polyploidy and genome evolution in plants. Curr Opin Genet Dev 35, 119–125 (2015).

65. Dunn, T. & Sethuraman, A. Accurate Inference of the Polyploid Continuum Using Forward-Time Simulations. Mol Biol Evol 41, (2024).

66. Freeling, M. Bias in Plant Gene Content Following Different Sorts of Duplication: Tandem, Whole-Genome, Segmental, or by Transposition. Annu Rev Plant Biol 60, 433–453 (2009).

67. Freeling, M. & Thomas, B. C. Gene-balanced duplications, like tetraploidy, provide predictable drive to increase morphological complexity. Genome Res 16, 805–814 (2006).

68. Qiao, X. et al. Different Modes of Gene Duplication Show Divergent Evolutionary Patterns and Contribute Differently to the Expansion of Gene Families Involved in Important Fruit Traits in Pear (Pyrus bretschneideri). Front Plant Sci 9, (2018).

69. Systematics, Evolution, and Biogeography of Compositae. (International Association for Plant Taxonomy, Vienna, 2009).

70. Zachos, J. C., Dickens, G. R. & Zeebe, R. E. An early Cenozoic perspective on greenhouse warming and carbon-cycle dynamics. Nature 451, 279–283 (2008).

71. Korasidis, V. A., Wing, S. L., Morse, P. E., Vitek, N. S. & Bloch, J. I. Evidence for increased animal pollination during the Paleocene–Eocene thermal maximum. Paleobiology 51, 574–587 (2025).

72. Ramsey, J. & Ramsey, T. S. Ecological studies of polyploidy in the 100 years following its discovery. Phil. Trans. R. Soc. B 369, 20130352 (2014).

73. Strong, D. R. & Ayres, D. R. Ecological and Evolutionary Misadventures of *Spartina*. Annu Rev Ecol Evol Syst 44, 389–410 (2013).

74. Martin, S. L. & Husband, B. C. Adaptation of diploid and tetraploid *Chamerion angustifolium* to elevation but not local environment. Evolution 67, 1780–1791 (2013).

75. Boalt, E., Arvanitis, L., Lehtilä, K. & Ehrlén, J. The association among herbivory tolerance, ploidy level, and herbivory pressure in *cardamine pratensis*. Evol Ecol 24, 1101–1113 (2010).

76. Thompson, J. N. & Merg, K. F. Evolution of polyploidy and the diversification of plant-pollinator interactions. Ecology 89, 2197–2206 (2008).

77. Oswald, B. P. & Nuismer, S. L. Neopolyploidy and pathogen resistance. Proc. R. Soc. B 274, 2393–2397 (2007).

78. Fan, W. et al. The genomes of chicory, endive, great burdock and yacon provide insights into Asteraceae palaeo-polyploidization history and plant inulin production. Mol Ecol Resour 22, 3124–3140 (2022).

79. Wen, X. et al. The *Chrysanthemum lavandulifolium* genome and the molecular mechanism underlying diverse capitulum types. Hortic Res 9, (2022).

80. Todesco, M. et al. Massive haplotypes underlie ecotypic differentiation in sunflowers. Nature 584, 602–607 (2020).

81. Huang, K. et al. The genomics of linkage drag in inbred lines of sunflower. Proc Natl Acad Sci USA 120, (2023).

82. Yang, J.-S., Qian, Z.-H., Shi, T., Li, Z.-Z. & Chen, J.-M. Chromosome-level genome assembly of the aquatic plant *Nymphoides indica* reveals transposable element bursts and NBS-LRR gene family expansion shedding light on its invasiveness. DNA Res. 29, (2022).

83. Minio, A., Cochetel, N., Vondras, A. M., Massonnet, M. & Cantu, D. Assembly of complete diploid-phased chromosomes from draft genome sequences. G3: Genes, Genomes, Genet. 12, (2022).

84. Andrews, S. FastQC: A Quality Control Tool for High Throughput Sequence Data. Available at https://www.bioinformatics.babraham.ac.uk/projects/fastqc/ (2010).

85. Schultz, D., Ebbert, M. & De Coster, W. PAUVRE. Available at https://github.com/conchoecia/pauvre/ (2019).

86. Chen, S., Zhou, Y., Chen, Y. & Gu, J. fastp: an ultra-fast all-in-one FASTQ preprocessor. Bioinformatics 34, i884–i890 (2018).

87. Cheng, H., Concepcion, G. T., Feng, X., Zhang, H. & Li, H. Haplotype-resolved de novo assembly using phased assembly graphs with hifiasm. Nat Methods 18, 170–175 (2021).

88. Zimin, A. V. et al. Hybrid assembly of the large and highly repetitive genome of *Aegilops tauschii*, a progenitor of bread wheat, with the MaSuRCA mega-reads algorithm. Genome Res 27, 787–792 (2017).

89. Langmead, B. & Salzberg, S. L. Fast gapped-read alignment with Bowtie 2. Nat Methods 9, 357–359 (2012).

90. Walker, B. J. et al. Pilon: An Integrated Tool for Comprehensive Microbial Variant Detection and Genome Assembly Improvement. PLoS One 9, e112963 (2014).

91. Li, H. Aligning sequence reads, clone sequences and assembly contigs with BWA-MEM. Preprint at 10.48550/arXiv.1303.3997 (2013).

92. Li, H. & Durbin, R. Fast and accurate short read alignment with Burrows–Wheeler transform. Bioinformatics 25, 1754–1760 (2009).

93. Danecek, P. et al. Twelve years of SAMtools and BCFtools. Gigascience 10, (2021).

94. Zhou, C., McCarthy, S. A. & Durbin, R. YaHS: yet another Hi-C scaffolding tool. Bioinformatics 39, (2023).

95. Durand, N. C. et al. Juicebox Provides a Visualization System for Hi-C Contact Maps with Unlimited Zoom. Cell Syst 3, 99–101 (2016).

96. Manni, M., Berkeley, M. R., Seppey, M., Simão, F. A. & Zdobnov, E. M. BUSCO Update: Novel and Streamlined Workflows along with Broader and Deeper Phylogenetic Coverage for Scoring of Eukaryotic, Prokaryotic, and Viral Genomes. Mol Biol Evol 38, 4647–4654 (2021).

97. Wood, D. E., Lu, J. & Langmead, B. Improved metagenomic analysis with Kraken 2. Genome Biol 20, 257 (2019).

98. Ou, S. et al. Benchmarking transposable element annotation methods for creation of a streamlined, comprehensive pipeline. Genome Biol 20, 275 (2019).

99. Flynn, J. M. et al. RepeatModeler2 for automated genomic discovery of transposable element families. Proc Natl Acad Sci USA 117, 9451–9457 (2020).

100. Smit, A. F. A., Hubley, R. & Green, P. RepeatMasker Open-4.0. Preprint at http://www.repeatmasker.org (2013).

101. Gabriel, L. et al. BRAKER3: Fully automated genome annotation using RNA-seq and protein evidence with GeneMark-ETP, AUGUSTUS and TSEBRA. Preprint at 10.1101/2023.06.10.544449 (2023).

102. Jones, P. et al. InterProScan 5: genome-scale protein function classification. Bioinformatics 30, 1236–1240 (2014).

103. Manchanda, N. et al. GenomeQC: a quality assessment tool for genome assemblies and gene structure annotations. BMC Genomics 21, 193 (2020).

104. Jin, J. J. et al. GetOrganelle: A fast and versatile toolkit for accurate de novo assembly of organelle genomes. Genome Biol 21, 1–31 (2020).

105. Zhou, W. et al. Plastid Genome Assembly Using Long-read data. Mol Ecol Resour 23, 1442–1457 (2023).

106. Tillich, M. et al. GeSeq-versatile and accurate annotation of organelle genomes. Nucleic Acids Res 45, (2017).

107. Darling, A. C. E., Mau, B., Blattner, F. R. & Perna, N. T. Mauve: Multiple Alignment of Conserved Genomic Sequence With Rearrangements. Genome Res 14, 1394–1403 (2004).

108. Marçais, G. & Kingsford, C. A fast, lock-free approach for efficient parallel counting of occurrences of k-mers. Bioinformatics 27, 764–770 (2011).

109. Ranallo-Benavidez, T. R., Jaron, K. S. & Schatz, M. C. GenomeScope 2.0 and Smudgeplot for reference-free profiling of polyploid genomes. Nat Commun 11, (2020).

110. Kokot, M., Dlugosz, M. & Deorowicz, S. KMC 3: counting and manipulating k-mer statistics. Bioinformatics 33, 2759–2761 (2017).

111. Emms, D. M. & Kelly, S. OrthoFinder: Phylogenetic orthology inference for comparative genomics. Genome Biol 20, (2019).

112. Minh, B. Q. et al. IQ-TREE 2: New Models and Efficient Methods for Phylogenetic Inference in the Genomic Era. Mol Biol Evol 37, 1530–1534 (2020).

113. Tang, H., et al. JCVI: A versatile toolkit for comparative genomics analysis. iMeta 3, (2024).

114. Jia, K. et al. Sub Phaser: a robust allopolyploid subgenome phasing method based on subgenome-specific k-mers. New Phytol 235, 801–809 (2022).

115. Sun, P. et al. WGDI: A user-friendly toolkit for evolutionary analyses of whole-genome duplications and ancestral karyotypes. Mol Plant 15, 1841–1851 (2022).

116. Zhang, C., Rabiee, M., Sayyari, E. & Mirarab, S. ASTRAL-III: Polynomial time species tree reconstruction from partially resolved gene trees. BMC Bioinf 19, 15–30 (2018).

117. Zhang, R.-G., Shang, H.-Y., Jia, K.-H. & Ma, Y.-P. Subgenome phasing for complex allopolyploidy: case-based benchmarking and recommendations. Briefings Bioinf 25, (2023).

118. Paten, B. et al. Cactus: Algorithms for genome multiple sequence alignment. Genome Res 21, 1512–1528 (2011).

119. Armstrong, J. et al. Progressive Cactus is a multiple-genome aligner for the thousand-genome era. Nature 587, 246–251 (2020).

120. Sullivan, P. F. et al. Leveraging base-pair mammalian constraint to understand genetic variation and human disease. Science 380, (2023).

121. Hubisz, M. J., Pollard, K. S. & Siepel, A. PHAST and RPHAST: phylogenetic analysis with space/time models. Briefings Bioinf 12, 41–51 (2011).

122. Siepel, A. et al. Evolutionarily conserved elements in vertebrate, insect, worm, and yeast genomes. Genome Res 15, 1034–1050 (2005).

123. Keough, K. C. et al. Three-dimensional genome rewiring in loci with human accelerated regions. Science 380, (2023).

124. Pollard, K. S., Hubisz, M. J., Rosenbloom, K. R. & Siepel, A. Detection of nonneutral substitution rates on mammalian phylogenies. Genome Res 20, 110–121 (2010).

125. Storey, J. D. & Tibshirani, R. Statistical significance for genomewide studies. Proc Natl Acad Sci USA 100, 9440–9445 (2003).

126. Cahill, J. A. et al. Positive selection in noncoding genomic regions of vocal learning birds is associated with genes implicated in vocal learning and speech functions in humans. Genome Res 31, 2035–2049 (2021).

127. Yocca, A. et al. A chromosome-scale assembly for ‘d’Anjou’ pear. G3: Genes, Genomes, Genet. 14, (2024).

128. Edgar, R. C. MUSCLE: Multiple sequence alignment with high accuracy and high throughput. Nucleic Acids Res 32, 1792–1797 (2004).

129. Suyama, M., Torrents, D. & Bork, P. PAL2NAL: Robust conversion of protein sequence alignments into the corresponding codon alignments. Nucleic Acids Res 34, (2006).

130. Yang, Z. PAML: A Program Package for Phylogenetic Analysis by Maximum Likelihood. CABIOS APPLICATIONS NOTE vol. 13 http://mw511.biol.berkeley.edu/ziheng/paml.html. (1997).

131. R Development Core Team (2008). R: A language and environment for statistical computing. R Foundation for Statistical Computing, Vienna, Austria. ISBN 3-900051-07-0, URL http://www.R-project.org. Available at https://stat.ethz.ch/pipermail/r-help/2008-May/161481.html.

132. R Core Team. R: A language and environment for statistical computing. Preprint at https://www.r-project.org/ (2017).

133. Wickham, H. *Ggplot2: Elegant Graphics for Data Analysis*. (Springer-Verlag, New York, 2016).

134. Sensalari, C., Maere, S. & Lohaus, R. ksrates: positioning whole-genome duplications relative to speciation events in Ks distributions. Bioinformatics 38, 530–532 (2022).

135. McKibben, M. T. W., Finch, G. & Barker, M. S. Species-tree topology impacts the inference of ancient whole-genome duplications across the angiosperm phylogeny. Am J Bot 111, (2024).

136. Chen, H. et al. Revisiting ancient polyploidy in leptosporangiate ferns. New Phytol 237, 1405–1417 (2023).

137. Pupko, T., Pe, I., Shamir, R. & Graur, D. A Fast Algorithm for Joint Reconstruction of Ancestral Amino Acid Sequences. Mol Biol Evol 17, 890–896 (2000).

138. Revell, L. J. phytools: An R package for phylogenetic comparative biology (and other things). Methods Ecol Evol 3, 217–223 (2012).

139. De Bie, T., Cristianini, N., Demuth, J. P. & Hahn, M. W. CAFE: a computational tool for the study of gene family evolution. Bioinformatics 22, 1269–1271 (2006).

140. Maurin, K. J. L. An empirical guide for producing a dated phylogeny with treePL in a maximum likelihood framework. (2020).

141. Smith, S. A. & O’Meara, B. C. treePL: divergence time estimation using penalized likelihood for large phylogenies. Bioinformatics 28, 2689–2690 (2012).

142. Barreda, V. D. et al. Early evolution of the angiosperm clade Asteraceae in the Cretaceous of Antarctica. Proc Natl Acad Sci USA 112, 10989–10994 (2015).

143. Barreda, V. D. et al. An extinct Eocene taxon of the daisy family (Asteraceae): evolutionary, ecological and biogeographical implications. Ann Bot 109, 127–134 (2012).

144. Hochuli, P. A. Palynologische Untersuchungen Im Oligozän Und Untermiozän Der Zentralen Und Westlichen Paratethys. (Geologischen Institut der Eidg. Technischen Hochschule und der Universtät Zürich, Zurich, 1978).

145. Beentje, H. J. & Caligari, P. D. S. Compositae: Proceedings of the International Compositae Conference, Kew, 1994. (Royal Botanic Gardens, 1996).

146. Magallón, S., Gómez-Acevedo, S., Sánchez-Reyes, L. L. & Hernández-Hernández, T. A metacalibrated time-tree documents the early rise of flowering plant phylogenetic diversity. New Phytol 207, 437–453 (2015).

147. Altschul, S. F., Gish, W., Miller, W., Myers, E. W. & Lipman, D. J. Basic local alignment search tool. J Mol Biol 215, 403–410 (1990).

148. Ceron-Noriega, A., Schoonenberg, V. A. C., Butter, F. & Levin, M. AlexandrusPS: A User-Friendly Pipeline for the Automated Detection of Orthologous Gene Clusters and Subsequent Positive Selection Analysis. Genome Biol Evol 15, (2023).

149. Ranwez, V., Harispe, S., Delsuc, F. & Douzery, E. J. P. MACSE: Multiple Alignment of Coding SEquences Accounting for Frameshifts and Stop Codons. PLoS One 6, e22594 (2011).

150. Kosakovsky Pond, S. L., et al. HyPhy 2.5—A Customizable Platform for Evolutionary Hypothesis Testing Using Phylogenies. Mol Biol Evol 37, 295–299 (2020).

151. Paradis, E. & Schliep, K. Ape 5.0: An environment for modern phylogenetics and evolutionary analyses in R. Bioinformatics 35, 526–528 (2019).

152. Yu, G., Smith, D. K., Zhu, H., Guan, Y. & Lam, T. T. ggtree: an r package for visualization and annotation of phylogenetic trees with their covariates and other associated data. Methods Ecol Evol 8, 28–36 (2017).

153. Ashburner, M. et al. Gene Ontology: tool for the unification of biology. Nat Genet 25, 25– 29 (2000).

154. Yu, G., Wang, L.-G., Han, Y. & He, Q.-Y. clusterProfiler: an R Package for Comparing Biological Themes Among Gene Clusters. OMICS 16, 284–287 (2012).

155. Yuan, B. et al. Identification and functional characterization of the *RPP13* gene family in potato (*Solanum tuberosum* L.) for disease resistance. Front Plant Sci 15, (2025).

156. Diaz-Ramirez, D. et al. Expression and Functional Analyses of the *WIP* Gene Family in Arabidopsis. Plants 11, 2010 (2022).

157. Tan, Y. et al. Functional Characterization of UDP-Glycosyltransferases Involved in Anti-viral Lignan Glycosides Biosynthesis in *Isatis indigotica*. Front Plant Sci 13, (2022).

158. Zhou, F.-Y. et al. Cytochrome *CYP72A15* may play a role in metabolic resistance to mesotrione in wild radish. Pestic Biochem Physiol 210, 106380 (2025).

159. Liu, Y., Koornneef, M. & Soppe, W. J. J. The Absence of Histone H2B Monoubiquitination in the Arabidopsis hub1 (rdo4) Mutant Reveals a Role for Chromatin Remodeling in Seed Dormancy. Plant Cell 19, 433–444 (2007).

160. Nath, K., O’Donnell, J. P. & Lu, Y. Chloroplastic iron-sulfur scaffold protein NFU3 is essential to overall plant fitness. Plant Signal Behav 12, e1282023 (2017).

161. Cao, Y. et al. Functional characterization of *NBS-LRR* genes reveals an *NBS-LRR* gene that mediates resistance against *Fusarium* wilt. BMC Biol 22, 45 (2024).

162. Li, C., et al. *OsSAUR2*, a small auxin-up RNA gene, is crucial for arsenic tolerance and accumulation in rice. Environ Exp Bot 226, 105894 (2024).

163. Prall, W., Hendy, O. & Thornton, L. E. Utility of a Phylogenetic Perspective in Structural Analysis of CYP72A Enzymes from Flowering Plants. PLoS One 11, e0163024 (2016).

164. Bakshi, M. & Oelmüller, R. WRKY transcription factors. Plant Signal Behav 9, e27700 (2014).

165. Baek, I.-S., Park, H.-Y., You, M. K., Lee, J. H. & Kim, J.-K. Functional Conservation and Divergence of FVE Genes that Control Flowering Time and Cold Response in Rice and Arabidopsis. Mol Cells 26, 368–372 (2008).

166. Chiang, Y.-H. et al. Functional Characterization of the GATA Transcription Factors GNC and CGA1 Reveals Their Key Role in Chloroplast Development, Growth, and Division in Arabidopsis. Plant Physiol 160, 332–348 (2012).

167. Yang, H. et al. Multifaceted roles of *FD* gene family in flowering, plant architecture, and adaptation. Front Plant Sci 16, (2025).

168. https://GBIF.org User. Occurrence Download. 10.15468/DL.RJ2PNP (2025).

169. https://GBIF.org User. Occurrence Download. 10.15468/DL.BRQ5TF (2025).

170. https://GBIF.org User. Occurrence Download. 10.15468/DL.66G87F (2025).

