## Supplementary Information for "Gene duplication dynamics and regulatory evolution shape the diversification of Asteraceae"

Supplementary Results and Discussion, along with Supplementary Table, Dataset, and Figure legends

### Table of Contents


Supplementary Results 4

Supplementary Discussion 10

Supplementary References 13

Supplementary Table, Dataset, and Figure legends 19

- Supplementary Table 1 | Newly assembled genome statistics. 19
- Supplementary Table 2 | Assembly statistics for the eight Asterales genomes. 19
- Supplementary Table 3 | Syntenic gene depth ratios of the eight Asterales taxa plus *Vitis*. 19
- Supplementary Table 4 | Triplications and duplications between Goodeniaceae, Calyceraceae, and Asteraceae. 19
- Supplementary Table 5 | Curated list of floral GO terms. 19
- Supplementary Table 6 | Detailed accession information. 19
- Supplementary Table 7 | Voucher information for the three genomes generated in this study. 19
- Supplementary Table 8 | Raw data statistics of the three genomes generated in this study. 20
- Supplementary Table 9 | Annotation quality results. 20
- Supplementary Table 10 | Proteome quality results. 20
- Supplementary Table 11 | InterProScan GO terms. 20
- Supplementary Table 12 | Gene depth ratios of generated subgenomes. 20
- Supplementary Table 13 | Re-annotated publicly available genome statistics. 20
- Supplementary Table 14 | Pairwise syntenic gene counts. 20
- Supplementary Dataset 1 | Subgenome syntenic gene information. 20
- Supplementary Dataset 2 | Subgenome locations in each Asterales genome tested. 20
- Supplementary Dataset 3 | Gene results from all analyses in this study. 20

Supplementary Dataset 4 | List of single-copy orthogroups. 21
Supplementary Dataset 5 | Accelerated subgenome genes. 21
Supplementary Dataset 6 | Subgenome accelerated regions. 21
Supplementary Fig. 1 | Juicebox contact maps. 21
Supplementary Fig. 2 | BUSCO genome assembly results. 21
Supplementary Fig. 3 | $K_S$ plots of paralogous and orthologous gene pairs. 21
Supplementary Fig. 4 | Orthologous and paralogous $K_S$ plot inferences. 21
Supplementary Fig. 5 | Rate-adjusted $K_S$ plots of paralogous and orthologous gene pairs. 22
Supplementary Fig. 6 | Estimated WGD placements from *ksrates*. 22
Supplementary Fig. 7 | Gene blocks retained across all chromosomes from WGDI. 22
Supplementary Fig. 8 | Subgenome collinearity, phylogeny, and gene fractionation results from chromosome 2 of *Vitis*. 22
Supplementary Fig. 9 | Subgenome collinearity, phylogeny, and gene fractionation results from chromosome 7 of *Vitis*. 22
Supplementary Fig. 10 | Subgenome collinearity, phylogeny, and gene fractionation results from chromosome 11 of *Vitis*. 23
Supplementary Fig. 11 | Subgenome collinearity, phylogeny, and gene fractionation results from chromosome 14 of *Vitis*. 23
Supplementary Fig. 12 | Subgenome collinearity, phylogeny, and gene fractionation results from chromosome 15 of *Vitis*. 23
Supplementary Fig. 13 | Subgenome gene composition of each Asterales species within each tested chromosome (chromosomes 2, 7, 11, 14, and 15). 24
Supplementary Fig. 14 | Results of the WHALE analysis. 24
Supplementary Fig. 15 | Gene expansions and contractions across Asterales. 24
Supplementary Fig. 16 | Plot of GO terms indicating most expanded, contracted, lost, and lineage-specific genes. 24
Supplementary Fig. 17 | Composition of each duplication type. 24
Supplementary Fig. 18 | *FAD2* gene family expansion in Asteraceae. 24
Supplementary Fig. 19 | *CYP71B23* gene family expansion in Asteraceae. 25
Supplementary Fig. 20 | GO enrichment plot of most positively selected genes in Asteraceae. 25
Supplementary Fig. 21 | *FLC*-like gene tree. 25
Supplementary Fig. 22 | Calibration points for WHALE analysis. 25

Supplementary Fig. 23 | Long-read sequence data quality scores. 25

Supplementary Fig. 24 | $K_S$ dotplots of the eight Asterales taxa against chromosome 11 of *Vitis*. 25

Supplementary Fig. 25 | Time-calibrated phylogeny of the eight Asterales taxa plus *Arabidopsis* and *Vitis*. 25

Supplementary Fig. 26 | Proteome quality assessment of *Acicarpha*, *Scaevola*, and *Platycodon*. 25

Supplementary Fig. 27 | Plastome of *Platycodon grandiflorus*. 25

Supplementary Fig. 28 | Plastome of *Nymphoides indica*. 26

Supplementary Fig. 29 | Plastome of *Scaevola aemula*. 26

Supplementary Fig. 30 | Plastome of *Acicarpha procumbens*. 26

Supplementary Fig. 31 | Plastome of *Arctium lappa*. 26

Supplementary Fig. 32 | Plastome of *Lactuca sativa*. 26

Supplementary Fig. 33 | Plastome of *Chrysanthemum lavandulifolium*. 26

Supplementary Fig. 34 | Plastome of *Helianthus annuus*. 26

Supplementary Fig. 35 | Plastome alignments indicating rearrangements. 26

Supplementary Fig. 36 | Genome composition plots of each Asterales taxon. 26

Supplementary Fig. 37 | Species tree of eight target taxa within Asterales plus *Arabidopsis* and *Vitis*. 26

Supplementary Fig. 38 | Accelerated subgenome evolution in Asterales relative to chromosome 11 of *Vitis*. 26

### Supplementary Results

#### Genome Sequencing

For *Acicarpha*, two Pacbio HiFi flow cells yielded 40 billion base pairs (bp) total of long read data, 187.86 billion bp of paired end data from Omni-C sequencing, and 53 billion bp of paired-end Illumina short-reads, resulting in 16x, 71.16x, and 20x coverage, respectively (Supplementary Table 8). For *Scaevola*, one Pacbio HiFi flow cell yielded ~23.4 billion bp of long read data, ~170.9 billion bp of paired end data from Hi-C sequencing, and 140 billion bp of paired-end Illumina short-read sequences, resulting in 19.5x, 213.6x, and 30x coverage, respectively (Supplementary Table 8). For *Platycodon*, one flow cell of Oxford Nanopore MinION sequencing resulted in ~14.46 billion bp of long read data, resulting in 24.6x coverage given a genome size estimate of 587 Mbp[1]. A sequenced Hi-C library yielded ~60.46 billion bp of data, resulting in 103x coverage of the genome, while Illumina sequencing yielded ~94 billion bp and 160x coverage of paired-end short read data (Supplementary Table 8).

#### Genome annotation and quality

*Platycodon*, *Scaevola*, and *Acicarpha* had fairly complete genome assemblies with BUSCO scores of 98%, 96.6%, and 98.2%, respectively (Supplementary Table 2). Out of those three taxa, *Scaevola* contained the most missing data (2.5%; Supplementary Fig. 2). Across the eight Asterales taxa, *Arctium* was the most complete genome with a BUSCO score of 99.1% (Supplementary Table 2), though *Lactuca* had the most complete and single-copy genes (Supplementary Fig. 2). *Chrysanthemum* had the lowest BUSCO score (92%), the most duplicated genes (27.1%), and was considered the most fragmented (1.5%) compared to the remaining genomes (Supplementary Table 2).

*Nymphoides* had the lowest number of annotated genes (29,527 genes), while *Chrysanthemum* had the most (84,764 genes). Average exon lengths ranged from 181 bp in *Chrysanthemum* to 261 bp in *Helianthus* (Supplementary Table 9). Of the newly generated genomes, *Scaevola* had the highest number of annotated genes compared to *Platycodon* and *Acicarpha*, though *Scaevola* also had the second lowest minimum gene length (~92 bp; Supplementary Table 9) and the highest proportion of unknown gene homologs. Otherwise, the genome annotation qualities were comparable among the three species (Supplementary Fig. 26, Supplementary Table 10). Investigating functional annotations with InterProScan identified 30 total GOTerms across all eight taxa. Of those, seven GO Terms were found among all eight taxa and only one was found exclusively in Asteraceae (membrane; GO:0016020) (Supplementary Table 10).

**Plastid and mitochondrial genome assemblies**

Asteraceae chloroplasts exhibited high degrees of similarity in terms of size (from 149 Kbp to 153 Kbp; Supplementary Figs. 27-34) and collinearity (Supplementary Fig. 35). *Acicarpha* and *Nymphoides* also exhibited plastome sizes within this range, and also exhibited high collinearity with each other, although neither exhibited the large double inversion in the large single copy (LSC) region of the plastome, a notable feature of Asteraceae plastomes[2–5]. Both *Scaevola* and *Platycodon* exhibited high degrees of rearrangements compared to the other taxa. As has been shown before, *Scaevola* exhibited a much larger plastome (184 Kbp), mainly through expansions of the LSC and inverted repeats, and displayed extreme structural reconfigurations, highly differentiating it from its closely related lineages[2]. Mitochondrial genomes varied in length from 333 Kbp for *Acicarpha,* 335 Kbp for *Scaevola,* and 1.24 Mbp for *Platycodon.*

**Genome structure**

K-mer frequency analyses for all genomes (except *Chrysanthemum,* due to lack of short-read sequencing data) resulted in the following estimated genome sizes: 567.2 Mbp for *Platycodon*, 474.7 Mbp for *Nymphoides*, 409.6 Mbp for *Scaevola*, 152.8 Mbp for *Acicarpha,* 641.5 Mbp for *Arctium,* 1.9 Gbp for *Lactuca,* and 1.1 Gbp for *Helianthus.* Low sequencing coverage and/or high error rates for some genomic data, along with high levels of repetitive sequences, may have contributed to the lower genome sizes estimated for *Scaevola, Acicarpha, Arctium, Lactuca,* and *Helianthus* as compared to their reported genome assembly sizes (Fig. 1; Supplementary Table 8). Smudgeplot analyses confirmed diploid levels for all taxa and revealed signatures of multiple sub-genomes for all taxa, except *Scaevola,* although this finding may be an artifact of its low sequencing coverage (Supplementary Fig. 36).

**Gene and genome duplications**

To investigate the number of single-copy versus duplicated genes across Asteraceae and in *Acicarpha*, we ran OrthoFinder[6] on the eight Asterales taxa plus *Vitis* and *Arabidopsis*. OrthoFinder identified 105,766 orthologous sequences across *Vitis*, *Arabidopsis*, and the eight Asterales taxa. Of those, gene trees were constructed from 29,145 orthogroups with only 45 orthogroups considered single-copy (Supplementary Dataset 4). OrthoFinder indicated that crown Asteraceae shared 3,269 gene duplication events across analyzed Asteraceae taxa, compared to the node splitting Asteraceae from *Acicarpha* which shares only 744 (Supplementary Fig. 37), consistent with one to multiple duplication events within Asteraceae's history. *Helianthus* and *Chrysanthemum* have 31,648 and 47,083 unique gene duplications, respectively, which is almost double the amount of gene duplications of *Arctium* (18,330), *Lactuca* (18,092), and *Acicarpha* (20,804) (Supplementary Fig. 37), indicating unique duplication events within their individual histories.

**Phasing subgenomes**

Given the putative WGD and gene duplication events (Fig. 2f), we further investigated polyploidy by phasing the genomes into subgenomes to better understand family-wide polyploidy events. Our eight taxa are currently recognized as diploids, so we could not use SubPhaser[7] as it is unable to accurately phase genomes that experienced older, paleopolyploidy events[8]. Using Whole-Genome Duplication Integrated analysis (WGDI)[9], we found that some chromosomes assembled into subgenomes more easily than others. For example, chromosome 11 had distinct subgenomes that retained similar gene composition across multiple chromosomes and spanned the entire chromosome, compared to subgenome 15 which was much more fragmented (Supplementary Figs. 8-12). Additionally, the subgenomes of *Acicarpha* and *Arctium* were consistently easier to reconstruct compared to *Chrysanthemum*, *Helianthus*, and *Lactuca* which proved more difficult (Supplementary Fig. 8-12). *Helianthus* was the most challenging to phase as the subgenomes were typically spread over multiple chromosomes and sometimes only three out of the six putative subgenomes could be phased (Supplementary Table 12, Supplementary Figs. 8-12). The *Helianthus* subgenomes presented lower gene retention, potentially as a result of recent duplications and accompanying rearrangements[10,11] (Supplementary Fig. 13). *Platycodon*, *Nymphoides*, and *Scaevola* did not generate any subgenomes, which is expected given the consistent 1:1 gene depth ratio (Supplementary Table 12).

**Accelerated gene evolution in *Acicarpha* and Asteraceae**

We attempted to create a reference-free whole genome alignment for all species using Progressive Cactus[12,13]; however, this attempt was largely unsuccessful, likely due to the lack of synteny among species as a result of WGD/WGT events and subsequent gene/sequence loss. Further, Progressive Cactus is not optimized to align highly divergent and repeat-rich plant genomes[14]. Therefore, we produced an alignment of the subgenomes corresponding to chromosome 11 of *Vitis*, given its high gene retention (Supplementary Fig. 13) and collinearity (Supplementary Fig. 10). Using this alignment, we identified regions of accelerated molecular evolution (accelerated regions; ARs) across three positions within the subgenome phylogeny: across all Asteraceae and *Acicarpha* subgenomes (Position 1; Supplementary Fig. 10), in the *Acicarpha*-specific duplication event (Position 2; Supplementary Fig. 10), and in *Acicarpha* within subgenome clade 1 (Position 3; Supplementary Fig. 10). This was carried out for each clade-specific analysis by first identifying subgenomic regions that were conserved in the outgroups using PHAST phastCons[15,16] and then by using PHAST phyloP[16,17] to identify which conserved regions showed evidence of rapid evolution in the clades of interest. We identified 106 ARs across all Asteraceae and *Acicarpha* subgenomes, 227 ARs associated with the *Acicarpha*-specific duplication, and 19 ARs associated with *Acicarpha* in Subgenome 1 (Supplementary Datasets 5 and 6). This corresponded to 59, 107, and 14 AR-associated genes, respectively (Supplementary Dataset 5).

We then identified major targets of accelerated evolution by identifying regions of high AR density (AR hotspots[18]). We found two AR hotspots in the analysis of all Asteraceae and *Acicarpha* subgenomes (Position 1; Supplementary Fig. 10), and two in the analysis of the *Acicarpha*-specific duplication event (Position 2; Supplementary Figs. 10, 38). No hotspots were found in the analysis of *Acicarpha* within the Subgenome 1 clade (Position 3; Supplementary Figs. 10, 38). Across Asteraceae and *Acicarpha*, genes that overlap the highest peaks within hotspots are known to function in auxin-regulation (*ARF7*[19,20]), floral organ identity (*HEN2*[21,22]), and root hair growth (*VLN4*[23,24]). In the *Acicarpha*-specific duplication, the hotspot genes associated with peaks are *CBP60E*, a tetratricopeptide repeat (TPR)-like superfamily protein, and *ARR10* (Supplementary Dataset 3, Supplementary Fig. 38).

**$K_S$ distributions**

To investigate the influence of major genome duplication events and positive selection on Asterales genomes, we assessed the non-synonymous ($K_a$) and synonymous ($K_S$) substitution rates between paralogous and orthologous gene pairs by identifying peaks in $K_S$ value distributions to infer histories of WGD. Through the comparison of paralogous gene pairs of each species, we found evidence for the most recent duplication events in *Chrysanthemum, Nymphoides, Scaevola,* and *Acicarpha* (Fig. 2c). Of these, the height of these $K_S$ value peaks, compared to other peaks in their respective $K_S$ distributions, suggests recent small-scale ditsributions. The high density of gene duplications and the low $K_S$ value of the most prominent peak of *Helianthus*, indicates a recent WGD within the lineage. Peaks identified along higher $K_S$-values, e.g., those of *Platycodon, Nymphoides*, and *Scaevola,* are likely signatures of the gamma event (Fig. 2c), the ancestral whole-genome triplication of the core eudicots[25–27], which has been consistently shown with other studies in Asterales[28,29]. We found high densities of gene duplications for *Acicarpha* and all Asteraceae, between $K_S$ values of 0.4 and 1.0 (Fig. 2c).

Due to the similarities in $K_S$ peak values, these results suggest a shared WGD among these taxa, or the occurrence of separate WGD around the same time.

### WGD hypothesis testing

Using the results of *ksrates* and prior knowledge from the literature of WGD events, we performed hypothesis-based testing using WHALE[30] with two models (critical and relaxed) to determine the placement of nine total WGD events on our phylogeny, split into three analyses (see main text for details). Prior to running WHALE, we filtered out the extremely large or small OrthoFinder gene families as suggested by McKibben et al.[31], which decreased the number of orthogroups from 49,719 to 15,758. This number was further reduced to 14,480 gene trees after removing gene families with uncertain gene-tree topologies.

### Evolution of gene family size

We identified significant expansion of 39 gene families at the ancestral branch of Asteraceae+*Acicarpha*, with only a single contraction and no evidence of gene losses or lineage-specific orthogroups (Supplementary Fig. 15, Supplementary Dataset 3). The expanded genes were enriched in functions related to nitrogen compound response, metal ion transport, and defense against fungal and bacterial pathogens. These orthogroups encompass diverse functional categories, including receptor-like kinases (e.g., *CRK8*[32], *CRK41*[33], *ARK3*[34]), CYP enzymes (e.g., *CYP71B37*[35], *CYP83B1*[36], *CYP82G1*[37]), and signaling components (e.g., *ATHK1*[38], *ATGLR2.3*[39]). Several genes are linked to stress responses, such as peroxidases, matrix metalloproteinases (MMPs), and nitrate transporters (*ATNRT2.4*[40]). Additionally, expansions include genes involved in photosynthesis (*PSAA* and *PSAB*[41]), secondary metabolism (FAD-binding Berberine family proteins[42]), and cell wall modification (e.g., *VLN2*[43] and *CER2*[44]).

In the case of *Acicarpha*, we detected 61 significantly expanded orthogroups enriched, however, those were not enriched in any specific biological process and pathway. Interestingly, 2,054 gene families were specific to *Acicarpha* (see Supplementary Dataset 3). The gene families contracted at the *Acicarpha* branch included terpene synthases and various reductases involved in response to oxidative stress. *Acicarpha* also lost several gene families related to pollen-pistil interaction and recognition of pollen and general cell recognition (mostly *WAKL* genes[45]), as well as various root growth factors.

The CAFE results also detected 151 orthogroups specific to Asteraceae (Supplementary Fig. 16, Supplementary Dataset 3). These orthogroups were enriched in GO terms related to polysaccharide metabolism, as well as pollen recognition, and included orthogroups with best

BLAST hits to several receptor-kinases (e.g., *ARK1* and *ARK2*, involved in root hair growth and root morphology, respectively[46]), pectin lyases, and CTP synthases. In contrast to expanded gene families, there were few (10) that underwent significant contractions, and we did not detect any complete losses (Supplementary Dataset 3). The significantly contracted Asteraceae orthogroups are enriched in biological processes related to pollen tube growth development, mainly through association with *PMEI2*, a protein that regulates pectin methylesterase in the cell walls of pollen tubes[47].

Notably, all Asteraceae, Calyceraceae, and Goodeniaceae taxa lack the *PII* gene, a universal regulator of nitrogen-carbon assimilation[48]. This pattern supports a previous report of *PII* loss in Asteraceae and Goodeniaceae[28] and further extends that finding by demonstrating that Calyceraceae also lacks this gene. The absence of *PII* across these families likely reflects a single ancestral loss predating their divergence, followed by lineage-specific duplications and selective shifts that shaped their distinct evolutionary trajectories.

**Positive selection in Asteraceae**

Other notable genes under selection in Asteraceae were: *ATCHR12, CHR11, CHR23, SYD* (SWI2/SNF2-like proteins involved in chromatin remodeling[49–52]), *PDF1.4* (a plant defensin gene[53]), *SWEET9* (a sucrose transporter and nectar secretion gene[54]), and *WUS*, a central regulator of meristem maintenance[55] (Supplementary Dataset 3). Given the role of *WUS* in promoting stem cell identity and maintaining floral meristem activity[56,57], this finding suggests evolutionary change in meristem determinacy and floral organ patterning. This may be particularly relevant in Asteraceae, where the capitulum inflorescence represents a key innovation, integrating multiple floral meristems into a highly organized and determinate structure[58,59].

***FLC*-like Asterales clades**

Selection on *FVE*, which activate and repress *FLC*, respectively[60–62], suggests the control of FLC expression may be a target of adaptive evolution. *FLC* is a floral repressor[63–65], and its homologs have been found almost exclusively in Brassicaceae, with the exception of *Beta vulgaris* (Amaranthaceae)[65–69]. Previous work in *Cichorium* and *Chrysanthemum* (Asteraceae) identified *FLC*-like genes that retain features functionally conserved with the *FLC* homolog found in *B. vulgaris* and Brassicaceae[65,68,70]. Across our genomes here, and by extending the analyses to a larger number of angiosperm genomes, we identified highly duplicated genes in the *FLC*-like clade that share a unique duplication pattern in Asterales with two major Asterales *FLC*-like clades recovered (Supplementary Fig. 21; see Jones et al.[71] for methods).

### Supplementary Discussion

#### Evidence for WGD

Synteny patterns did not follow a strong phylogenetic signal (Fig. 1d), which is unsurprising given that the analyzed genomes span five families across Asterales and differ in quality and resolution (Supplementary Tables 1, 2, 9, and 13). In Asteraceae, synteny may be particularly low due to the high repeat content[72] (Fig. 1c) and extensive history of duplications within the family[73]. Additionally, the *Chrysanthemum* genome assembly was more fragmented and contained more missing data compared to the other genomes used in this study (Supplementary Tables 2 and 13, Supplementary Fig. 2), potentially explaining its low synteny[74]. It is possible that JCVI did not identify all synteny blocks as MCScanX, the tool used in JCVI for synteny detection, has been shown to recover fewer syntenic regions than some other softwares[74].

The $K_S$ plots show a large *Acicarpha* $K_S$ peak within the larger peak ranges of the Asteraceae taxa, with *Helianthus* and *Chrysanthemum* having peaks closer to zero, and therefore indicating more recent duplication events than *Acicarpha*, and *Arctium* and *Lactuca* having slightly older. *Helianthus* and *Chrysanthemum* are already suspected of having more recent WGD/T events since the origination of Asteraceae[10,11], so a more recent peak for *Acicarpha* than the remaining Asteraceae taxa may indicate that *Acicarpha* has a more recent and unique duplication event than its split from Goodeniaceae, which has been hypothesized previously[75] (Fig. 2f). This hypothesis is also supported with our synteny analysis as we see gene duplications from *Acicarpha* to *Arctium*, and vice versa, with genes that were triplicated since the split from Goodeniaceae (Fig. 2b, Supplementary Table 4).

To more accurately place WGD events and to directly compare peaks, we ran *ksrates* which investigates non-synonymous substitution rates relative to the taxa used in the study. In doing so, we identified eight tentative events, all of which do not fully align with previous studies. Most notably, *ksrates* did not identify the shared WGD/T event that has previously been proposed between Calyceraceae and Asteraceae[10,11,29]. Instead, the event was shared with Goodeniaceae, Calyceraceae, and Asteraceae.

Given the different WGD placements from multiple analyses in this study, we tested WGD event hypotheses using the results from *ksrates*, primary literature, and a combination of both (Supplementary Fig. 22). In doing so, we found model disagreement on the placements of

WGD events, where the critical model identified WGD events at only five branches, while the relaxed model identified WGD events at all tested branches (Supplementary Fig. 14). These findings are consistent with other studies that report that the critical model tends to underestimate the number of WGD events compared to the relaxed model, while the relaxed model tends to hypothesize a species-specific WGD bias[30,31].

**Phasing subgenomes**

This is the first study to generate a *de novo* alignment across highly diverged plant lineages. Previous studies have performed *de novo* alignments across the same plant species, varieties, or cultivars[76–78]; however, no other study to our knowledge has attempted *de novo* alignments across multiple distantly related plant species given challenges associated with chromosome rearrangements and fractionation, high sequence diversity, widespread structural variation, and high repeat content[15,79]. Our study overcame these issues by phasing the genomes into subgenomes and then generating a *de novo* alignment with the subgenomes, minimizing the issues detailed above, though it is important to note that the *de novo* alignments were still not complete and present some limitations. For example, we were only able to phase a small portion of the genomes into subgenomes (e.g., chromosome 11 of *Vitis*), limiting potential AR hotspots within our target clades. Even so, we addressed these challenges to the extent possible, given the complexities of plant genomes outlined above. We view this analysis as a proof of concept demonstrating the potential of more complete alignments. Future work would greatly benefit from the development of plant-specific *de novo* alignment tools designed to accommodate the substantial structural and sequence diversity characteristic of plant genomes.

**Key traits underpinning diversification in *Acicarpha***

AR analyses point to the relevance of stress and developmental regulation. For instance, *ARR10*, an AR hotspot gene in the *Acicarpha*-specific analysis, is a well-known regulator of drought response[80], and plays a critical role in meristem activity. Specifically, *ARR10* directly activates *WUS* expression by binding to its promoter[81–84], while simultaneously repressing *YUC* genes that encode key enzymes in auxin biosynthesis[85]. These dual functions suggest that *ARR10* fine-tunes the balance between hormone signaling and developmental fate, consistent with the broader patterns of auxin- and ABA-related gene evolution in *Acicarpha* described above.

Additional evidence for the importance of hormonal signaling in *Acicarpha* comes from the *Acicarpha*-specific gene family expansion of *ARF5* (also known as *MP*) and acceleration of *ARF7* (Supplementary Dataset 3). *ARF* genes play pivotal roles in root development, with distinct members contributing to specific root types. For example, *ARF5* is a key transcription factor in the auxin pathway that regulates vascular patterning, embryogenesis, and lateral root development, further supporting a shift in developmental regulation through auxin-responsive networks. Its interaction with auxin-responsive genes also complements the selection patterns seen with *IBR5* and *NPH3*, underscoring the importance of fine-tuning auxin signaling in *Acicarpha*.

The prominence of *CRK8* gene family members in the *Acicarpha*-specific and Asteraceae-*Acicarpha* CAFE analysis highlights the lineage's adaptation to oxidative and abiotic stress. *CRK8*, part of a receptor-like kinase family, has been implicated in oxidative stress tolerance and the regulation of reactive oxygen species, which play dual roles in signaling and cellular damage during environmental stress[86]. The expansion of *CRK8* suggests that *Acicarpha* may have evolved enhanced mechanisms for stress sensing and response, potentially contributing to its survival in fluctuating or challenging environments.

### Supplementary References


1. Zhang, L., Cao, B. & Bai, C. New reports of nuclear DNA content for 66 traditional Chinese medicinal plant taxa in China. *Caryologia* **66**, 375–383 (2013).
2. Pascual-Díaz, J. P., Garcia, S. & Vitales, D. Plastome Diversity and Phylogenomic Relationships in Asteraceae. *Plants* **10**, 2699 (2021).
3. Kim, K.-J., Choi, K.-S. & Jansen, R. K. Two Chloroplast DNA Inversions Originated Simultaneously During the Early Evolution of the Sunflower Family (Asteraceae). *Mol Biol Evol* **22**, 1783–1792 (2005).
4. Loeuille, B. *et al.* Extremely low nucleotide diversity among thirty-six new chloroplast genome sequences from *Aldama* (Heliantheae, Asteraceae) and comparative chloroplast genomics analyses with closely related genera. *PeerJ* **9**, e10886 (2021).
5. Timme, R. E., Kuehl, J. V., Boore, J. L. & Jansen, R. K. A comparative analysis of the *Lactuca* and *Helianthus* (Asteraceae) plastid genomes: identification of divergent regions and categorization of shared repeats. *Am J Bot* **94**, 302–312 (2007).
6. Emms, D. M. & Kelly, S. OrthoFinder: Phylogenetic orthology inference for comparative genomics. *Genome Biol* **20**, (2019).
7. Jia, K. *et al.* SubPhaser: a robust allopolyploid subgenome phasing method based on subgenome-specific k-mers. *New Phytol* **235**, 801–809 (2022).
8. Zhang, R.-G., Shang, H.-Y., Jia, K.-H. & Ma, Y.-P. Subgenome phasing for complex allopolyploidy: case-based benchmarking and recommendations. *Briefings Bioinf* **25**, (2023).
9. Sun, P. *et al.* WGDI: A user-friendly toolkit for evolutionary analyses of whole-genome duplications and ancestral karyotypes. *Mol Plant* **15**, 1841–1851 (2022).
10. Zhang, C. *et al.* Phylotranscriptomic insights into Asteraceae diversity, polyploidy, and morphological innovation. *J Integr Plant Biol* **63**, 1273–1293 (2021).
11. Huang, C.-H. *et al.* Multiple Polyploidization Events across Asteraceae with Two Nested Events in the Early History Revealed by Nuclear Phylogenomics. *Mol Biol Evol* **33**, 2820–2835 (2016).
12. Armstrong, J. *et al.* Progressive Cactus is a multiple-genome aligner for the thousand-genome era. *Nature* **587**, 246–251 (2020).
13. Paten, B. *et al.* Cactus: Algorithms for genome multiple sequence alignment. *Genome Res* **21**, 1512–1528 (2011).
14. Zhou, H., Su, X. & Song, B. ACMGA: a reference-free multiple-genome alignment pipeline for plant species. *BMC Genomics* **25**, 515 (2024).
15. Siepel, A. *et al.* Evolutionarily conserved elements in vertebrate, insect, worm, and yeast genomes. *Genome Res* **15**, 1034–1050 (2005).
16. Hubisz, M. J., Pollard, K. S. & Siepel, A. PHAST and RPHAST: phylogenetic analysis with space/time models. *Briefings Bioinf* **12**, 41–51 (2011).
17. Pollard, K. S., Hubisz, M. J., Rosenbloom, K. R. & Siepel, A. Detection of nonneutral substitution rates on mammalian phylogenies. *Genome Res* **20**, 110–121 (2010).
18. Cahill, J. A. *et al.* Positive selection in noncoding genomic regions of vocal learning birds is associated with genes implicated in vocal learning and speech functions in humans. *Genome Res* **31**, 2035–2049 (2021).

19. Okushima, Y. *et al.* Functional Genomic Analysis of the *AUXIN RESPONSE FACTOR* Gene Family Members in Arabidopsis thaliana: Unique and Overlapping Functions of *ARF7* and *ARF19*. *Plant Cell* **17**, 444–463 (2005).
20. Okushima, Y., Fukaki, H., Onoda, M., Theologis, A. & Tasaka, M. ARF7 and ARF19 Regulate Lateral Root Formation via Direct Activation of *LBD/ASL* Genes in *Arabidopsis*. *Plant Cell* **19**, 118–130 (2007).
21. Chen, X., Liu, J., Cheng, Y. & Jia, D. HEN1 functions pleiotropically in Arabidopsis development and acts in C function in the flower. *Development* **129**, 1085–1094 (2002).
22. Western, T. L., Cheng, Y., Liu, J. & Chen, X. HUA ENHANCER2, a putative DExH-box RNA helicase, maintains homeotic B and C gene expression in *Arabidopsis*. *Development* **129**, 1569–1581 (2002).
23. Zhang, Y. *et al.* Arabidopsis VILLIN4 is involved in root hair growth through regulating actin organization in a Ca2+-dependent manner. *New Phytol* **190**, 667–682 (2011).
24. Li, M. *et al.* Dynamics of Actin Filaments Play an Important Role in Root Hair Growth under Low Potassium Stress in *Arabidopsis thaliana*. *Int J Mol Sci* **25**, 8950 (2024).
25. Jiao, Y. *et al.* Ancestral polyploidy in seed plants and angiosperms. *Nature* **473**, 97–100 (2011).
26. Alix, K., Gérard, P. R., Schwarzacher, T. & Heslop-Harrison, J. S. (Pat). Polyploidy and interspecific hybridization: partners for adaptation, speciation and evolution in plants. *Ann Bot* **120**, 183–194 (2017).
27. Bowers, J. E., Chapman, B. A., Rong, J. & Paterson, A. H. Unravelling angiosperm genome evolution by phylogenetic analysis of chromosomal duplication events. *Nature* **422**, 433–438 (2003).
28. Shen, F. *et al.* Comparative genomics reveals a unique nitrogen-carbon balance system in Asteraceae. *Nat Commun* **14**, (2023).
29. Barker, M. S. *et al.* Most Compositae (Asteraceae) are descendants of a paleohexaploid and all share a paleotetraploid ancestor with the Calyceraceae. *Am J Bot* **103**, 1203–1211 (2016).
30. Zwaenepoel, A. & Van de Peer, Y. Inference of Ancient Whole-Genome Duplications and the Evolution of Gene Duplication and Loss Rates. *Mol Biol Evol* **36**, 1384–1404 (2019).
31. McKibben, M. T. W., Finch, G. & Barker, M. S. Species-tree topology impacts the inference of ancient whole-genome duplications across the angiosperm phylogeny. *Am J Bot* **111**, (2024).
32. Idänheimo, N. *et al.* The *Arabidopsis thaliana* cysteine-rich receptor-like kinases CRK6 and CRK7 protect against apoplastic oxidative stress. *Biochem. Biophys Res Commun* **445**, 457–462 (2014).
33. Li, X., Zhao, J., Sun, Y. & Li, Y. *Arabidopsis thaliana* CRK41 negatively regulates salt tolerance via H2O2 and ABA cross-linked networks. *Environ Exp Bot* **179**,104210 (2020).
34. Sakai, T. *et al.* Armadillo repeat-containing kinesins and a NIMA-related kinase are required for epidermal-cell morphogenesis in *Arabidopsis*. *Plant J* **53**, 157–171 (2008).
35. Wei, Y. *et al.* The susceptibility factor *Resistance To Phytophthora parasitica 1* negatively regulates Arabidopsis immunity by interacting with the cytochrome P450 protein CYP71B3. *Plant Physiol* **198**, (2025).

36. Naur, P. *et al.* CYP83A1 and CYP83B1, Two Nonredundant Cytochrome P450 Enzymes Metabolizing Oximes in the Biosynthesis of Glucosinolates in Arabidopsis. *Plant Physiol* **133**, 63–72 (2003).
37. Lee, S. *et al.* Herbivore-induced and floral homoterpene volatiles are biosynthesized by a single P450 enzyme (CYP82G1) in *Arabidopsis*. *Proc Natl Acad Sci USA* **107**, 21205–21210 (2010).
38. Wohlbach, D. J., Quirino, B. F. & Sussman, M. R. Analysis of the Arabidopsis histidine kinase ATHK1 reveals a connection between vegetative osmotic stress sensing and seed maturation. *Plant Cell* **20**, 1101–1117 (2008).
39. Weiland, M., Mancuso, S. & Baluska, F. Signaling via glutamate and GLRs in Arabidopsis thaliana. *Funct Plant Biol* **43**, 1–25 (2015).
40. Zhang, H. *et al.* Beyond nitrate transport: AtNRT2.4 responds to local and systemic nitrogen signaling in *Arabidopsis*. *BMC Plant Biol* **25**, 655 (2025).
41. Liu, J. *et al.* PSBP-DOMAIN PROTEIN1, a Nuclear-Encoded Thylakoid Lumenal Protein, Is Essential for Photosystem I Assembly in *Arabidopsis*. *Plant Cell* **24**, 4992–5006 (2013).
42. Tjallinks, G., Mattevi, A. & Fraaije, M. W. Biosynthetic Strategies of Berberine Bridge Enzyme-like Flavoprotein Oxidases toward Structural Diversification in Natural Product Biosynthesis. *Biochemistry* **63**, 2089–2110 (2024).
43. van der Honing, H. S., Kieft, H., Emons, A. M. C. & Ketelaar, T. Arabidopsis VILLIN2 and VILLIN3 Are Required for the Generation of Thick Actin Filament Bundles and for Directional Organ Growth. *Plant Physiol* **158**, 1426–1438 (2012).
44. Negruk, V., Yang, P., Subramanian, M., McNevin, J. P. & Lemieux, B. Molecular cloning and characterization of the CER2 gene of Arabidopsis thaliana. *Plant J* **9**, 137–145 (1996).
45. Wang, Z. *et al.* Comparative genomics analysis of WAK/WAKL family in Rosaceae identify candidate WAKs involved in the resistance to *Botrytis cinerea*. *BMC Genomics* **24**, 337 (2023).
46. Yoo, C.-M. & Blancaflor, E. B. Overlapping and divergent signaling pathways for ARK1 and AGD1 in the control of root hair polarity in *Arabidopsis thaliana*. *Front Plant Sci* **4**, (2013).
47. Giovane, A. *et al.* Pectin methylesterase inhibitor. *Biochim Biophys Acta* **1696**, 245–252 (2004).
48. Hsieh, M.-H., Lam, H.-M., van de Loo, F. J. & Coruzzi, G. A PII-like protein in *Arabidopsis*: Putative role in nitrogen sensing. *Proc Natl Acad Sci USA* **95**, 13965–13970 (1998).
49. Shu, J. *et al.* Genome-wide occupancy of Arabidopsis SWI/SNF chromatin remodeler SPLAYED provides insights into its interplay with its close homolog BRAHMA and Polycomb proteins. *Plant J* **106**, 200–213 (2021).
50. Sang, Y. *et al.* Mutations in two non-canonical Arabidopsis SWI2/SNF2 chromatin remodeling ATPases cause embryogenesis and stem cell maintenance defects. *Plant J* **72**, 1000–1014 (2012).
51. Folta, A. *et al.* Over-expression of Arabidopsis AtCHR23 chromatin remodeling ATPase results in increased variability of growth and gene expression. *BMC Plant Biol* **14**, 76 (2014).

52. Huanca-Mamani, W., Garcia-Aguilar, M., León-Martínez, G., Grossniklaus, U. & Vielle-Calzada, J.-P. CHR11, a chromatin-remodeling factor essential for nuclear proliferation during female gametogenesis in *Arabidopsis thaliana*. *Proc Natl Acad Sci USA* **102**, 17231–17236 (2005).
53. Liu, Y. *et al.* Genome-scale identification of *plant defensin* (*PDF*) family genes and molecular characterization of their responses to diverse nutrient stresses in allotetraploid rapeseed. *PeerJ* **9**, e12007 (2021).
54. Wang, J., Xue, X., Zeng, H., Li, J. & Chen, L. Sucrose rather than GA transported by AtSWEET13 and AtSWEET14 supports pollen fitness at late anther development stages. *New Phytol* **236**, 525–537 (2022).
55. Jha, P., Ochatt, S. J. & Kumar, V. WUSCHEL: a master regulator in plant growth signaling. *Plant Cell Rep* **39**, 431–444 (2020).
56. Schoof, H. *et al.* The stem cell population of Arabidopsis shoot meristems is maintained by a regulatory loop between the CLAVATA and WUSCHEL genes. *Cell* **100**, 635–644 (2000).
57. Lenhard, M., Bohnert, A., Jürgens, G. & Laux, T. Termination of Stem Cell Maintenance in Arabidopsis Floral Meristems by Interactions between *WUSCHEL* and *AGAMOUS*. *Cell* **105**, 805–814 (2001).
58. *Systematics, Evolution, and Biogeography of Compositae*. (International Association for Plant Taxonomy, Vienna, 2009).
59. Zhang, T. & Elomaa, P. Development and evolution of the Asteraceae capitulum. *New Phytol* **242**, 33–48 (2024).
60. Lee, J. & Amasino, R. M. Two FLX family members are non-redundantly required to establish the vernalization requirement in Arabidopsis. *Nat Commun* **4**, 2186 (2013).
61. Geraldo, N., Bäurle, I., Kidou, S., Hu, X. & Dean, C. FRIGIDA Delays Flowering in Arabidopsis via a Cotranscriptional Mechanism Involving Direct Interaction with the Nuclear Cap-Binding Complex. *Plant Physiol* **150**, 1611–1618 (2009).
62. Jeon, J. & Kim, J. FVE, an Arabidopsis Homologue of the Retinoblastoma-Associated Protein That Regulates Flowering Time and Cold Response, Binds to Chromatin as a Large Multiprotein Complex. *Mol Cells* **32**, 227–234 (2011).
63. Deng, W. *et al.* FLOWERING LOCUS C (FLC) regulates development pathways throughout the life cycle of Arabidopsis. *Proc Natl Acad Sci USA* **108**, 6680–6685 (2011).
64. Sheldon, C. C., Rouse, D. T., Finnegan, E. J., Peacock, W. J. & Dennis, E. S. The molecular basis of vernalization: The central role of FLOWERING LOCUS C (FLC). *Proc Natl Acad Sci USA* **97**, 3753–3758 (2000).
65. Périlleux, C. *et al.* A root chicory MADS box sequence and the Arabidopsis flowering repressor FLC share common features that suggest conserved function in vernalization and de-vernalization responses. *Plant J* **75**, 390–402 (2013).
66. Blackman, B. K. *et al.* Contributions of Flowering Time Genes to Sunflower Domestication and Improvement. *Genetics* **187**, 271–287 (2011).
67. Becker, A. The major clades of MADS-box genes and their role in the development and evolution of flowering plants. *Mol Phylogenet Evol* **29**, 464–489 (2003).

68. Reeves, P. A. *et al.* Evolutionary Conservation of the FLOWERING LOCUS C-Mediated Vernalization Response: Evidence From the Sugar Beet (*Beta vulgaris*). *Genetics* **176**, 295–307 (2007).
69. Hileman, L. C. *et al.* Molecular and Phylogenetic Analyses of the MADS-Box Gene Family in Tomato. *Mol Biol Evol* **23**, 2245–2258 (2006).
70. Hu, Q. *et al.* Late flowering in chrysanthemum induced by low ambient temperature is mediated by FLOWERING LOCUS C-like. *J Exp Bot* **76**, 2192–2206 (2025).
71. Jones, D. S. *et al.* Floral innovation through modifications in stem cell peptide signaling. Preprint at https://doi.org/10.1101/2025.06.27.661788 (2025).
72. Staton, S. E. & Burke, J. M. Evolutionary transitions in the Asteraceae coincide with marked shifts in transposable element abundance. *BMC Genomics* **16**, 623 (2015).
73. Palazzesi, L. *et al.* Asteraceae as a model system for evolutionary studies: from fossils to genomes. *Bot J Linn Soc* **200**, 143–164 (2022).
74. Liu, D., Hunt, M. & Tsai, I. J. Inferring synteny between genome assemblies: a systematic evaluation. *BMC Bioinformatics* **19**, 26 (2018).
75. Zhang, C. *et al.* Asterid Phylogenomics/Phylotranscriptomics Uncover Morphological Evolutionary Histories and Support Phylogenetic Placement for Numerous Whole-Genome Duplications. *Mol Biol Evol* **37**, 3188–3210 (2020).
76. Hufford, M. B. *et al.* De novo assembly, annotation, and comparative analysis of 26 diverse maize genomes. *Science* **373**, 655–662 (2021).
77. Du, H. *et al.* Sequencing and de novo assembly of a near complete indica rice genome. *Nat Commun* **8**, 15324 (2017).
78. Jiao, W.-B. & Schneeberger, K. Chromosome-level assemblies of multiple *Arabidopsis* genomes reveal hotspots of rearrangements with altered evolutionary dynamics. *Nat Commun* **11**, 989 (2020).
79. Song, B., Buckler, E. S. & Stitzer, M. C. New whole-genome alignment tools are needed for tapping into plant diversity. *Trends Plant Sci* **29**, 355–369 (2024).
80. Nguyen, K. H. *et al.* Arabidopsis type B cytokinin response regulators ARR1, ARR10, and ARR12 negatively regulate plant responses to drought. *Proc Natl Acad Sci USA* **113**, 3090–3095 (2016).
81. Zhu, M. *et al.* The type-B response regulators ARR10, ARR12, and ARR18 specify the central cell in Arabidopsis. *Plant Cell* **34**, 4714–4737 (2022).
82. Liu, Z. *et al.* The Type-B Cytokinin Response Regulator ARR1 Inhibits Shoot Regeneration in an ARR12-Dependent Manner in Arabidopsis. *Plant Cell* **32**, 2271–2291 (2020).
83. Xie, M. *et al.* Author Correction: A B-ARR-mediated cytokinin transcriptional network directs hormone cross-regulation and shoot development. *Nat Commun* **14**, 5988 (2023).
84. Zubo, Y. O. *et al.* Cytokinin induces genome-wide binding of the type-B response regulator ARR10 to regulate growth and development in *Arabidopsis*. *Proc Natl Acad Sci USA* **114**, (2017).
85. Meng, W. J. *et al.* Type-B ARABIDOPSIS RESPONSE REGULATORs Specify the Shoot Stem Cell Niche by Dual Regulation of WUSCHEL. *Plant Cell* **29**, 1357–1372 (2017).

86. Wrzaczek, M. *et al.* Transcriptional regulation of the CRK/DUF26 group of Receptor-like protein kinases by ozone and plant hormones in Arabidopsis. *BMC Plant Biol* **10**, 95 (2010).

### Supplementary Table, Dataset, and Figure legends

**Supplementary Table 1 | Newly assembled genome statistics.** Genome statistics generated from a custom python script (https://github.com/lexnederbragt/sequencetools/blob/master/assemblathon_stats.pl) for the three genomes generated in this study: *Acicarpha procumbens*, *Scaevola aemula*, and *Platycodon grandiflorus*.

**Supplementary Table 2 | Assembly statistics for the eight Asterales genomes.** Haploid chromosome numbers and genome sizes were obtained from the original publication of that assembly. BUSCO scores were calculated in this study: C = complete; S = single-copy; D = duplicated; F = fragmented; M = missing; n = number of genes from embryophyta_10. The number of total base pairs (bp), the number of masked bp, and percent (%) masked are for the entire assembly. Columns J-O are DNA and repeat content values as indicated by EDTA and visualized in Fig. 1e. Asterisk (*) indicates that the genomes were generated in this study.

**Supplementary Table 3 | Syntenic gene depth ratios of the eight Asterales taxa plus *Vitis*.** Gene depth ratios generated with JCVI. The left numerical value is for the species on the y-axis, while the right numerical value is for the species on the top, x-axis (e.g., the gene depth ratio of *Vitis*:*Platycodon* is 5:3 for the c-score of 0.5). The top, middle, and bottom matrix used c-score value cutoffs of 0.5, 0.7, and 0.99, respectively.

**Supplementary Table 4 | Triplications and duplications between Goodeniaceae, Calyceraceae, and Asteraceae**. Number of triplications (3x) or duplications (2x) between Goodeniaceae (*Scaevola*), Calyceraceae (*Acicarpha*), and Asteraceae (*Arctium*). Numbers in parentheses correspond to the percentage of those duplicated or triplicated genes in the entire genome. Numbers originated from JCVI.

**Supplementary Table 5 | Curated list of floral GO terms.** Biological Process GO terms associated with reproductive development. Column F, 'Orthogroup', corresponds to our OrthoFinder run containing the eight Asterales taxa plus *Arabidopsis* and *Vitis*.

**Supplementary Table 6 | Detailed accession information.** Accession information for all raw and assembled data used in this study.

**Supplementary Table 7 | Voucher information for the three genomes generated in this study.** This includes *Platycodon grandiflorus*, *Scaevola aemula*, and *Acicarpha procumbens*.

**Supplementary Table 8 | Raw data statistics of the three genomes generated in this study.** 'bp' = base pairs. Numbers in parentheses refer to the sequence coverage depth given the estimated genome sizes.

**Supplementary Table 9 | Annotation quality results.** GenomeQC results for all eight genome annotations used in this study. Asterisk (*) next to species name indicates that the genomes were generated in this study.

**Supplementary Table 10 | Proteome quality results.** OMArk results used to assess the proteome quality of *Acicarpha*, *Scaevola*, and *Platycodon*.

**Supplementary Table 11 | InterProScan GO terms.** List of 30 GO terms annotated by InterProScan for each species. NA indicates that the GO term was not found in that species.

**Supplementary Table 12 | Gene depth ratios of generated subgenomes.** Gene depth ratios of the eight Asterales genomes against chromosomes 2, 7, 11, 14, and 15 of *Vitis*. Bolded ratios indicate differing ratios between the different subgenomes. Gene depths were determined via dotplots generated from WGDI.

**Supplementary Table 13 | Re-annotated publicly available genome statistics.** Genome statistics generated from a custom python script (https://github.com/lexnederbragt/sequencetools/blob/master/assemblathon_stats.pl) for the five publicly available genomes used in this study: *Arctium lappa*, *Chrysanthemum lavandulifolium*, *Helianthus annuus*, *Lactuca sativa*, and *Nymphoides indica*.

**Supplementary Table 14 | Pairwise syntenic gene counts.** Matrix showing the total number of syntenic gene regions found between each species used in this study with JCVI. Bolded numbers indicate the pairwise synteny that is represented in Fig. 1. Cells are colored by lowest number of syntenic regions (green) to highest (yellow).

**Supplementary Dataset 1 | Subgenome syntenic gene information.** Syntenic gene matches of the eight Asterales taxa and their subgenomes to the various chromosomes of *Vitis*. An empty cell indicates a lack of a gene match for the *Vitis* gene.

**Supplementary Dataset 2 | Subgenome locations in each Asterales genome tested.** Color of subgenome number matches the colors as indicated in Supplementary Fig. 4.

**Supplementary Dataset 3 | Gene results from all analyses in this study.** Category corresponds to the subanalysis and Method to the software/tool used. TAIR gene, symbol, description, and length came from BLASTP. Duplication type came from McScanX. NA indicates an empty cell with no information.

**Supplementary Dataset 4 | List of single-copy orthogroups.** List of 45 single-copy gene families or orthogroups identified by OrthoFinder. TAIR, Symbol, and Description came from BLASTP.

**Supplementary Dataset 5 | Accelerated subgenome genes.** Genes associated with accelerated subgenomic regions in the various subgenome analyses (see Supplementary Fig. 38). A gene was associated with an accelerated region (AR) if it landed within an exon. ARs may cluster into high-density regions known as hotspots, and AR-associated genes within such regions are indicated in the table.

**Supplementary Dataset 6 | Subgenome accelerated regions.** Accelerated regions (ARs) in the various subgenome analyses (see Supplementary Fig. 38). Subgenomes in Figure 2d-e represent genomic sequences that align to chromosome 11 of *Vitis*. Multiple hypothesis testing corrections were performed by generating Q-values using Storey's correction method.

**Supplementary Fig. 1 | Juicebox contact maps**. Plots of **a)** *Scaevola* haplotype 1, **b)** *Scaevola* haplotype 2, **c)** *Acicarpha* haplotype 1, **d)** *Acicarpha* haplotype 2, and **e)** *Platycodon*. Red horizontal line indicates continuous sequences. Blue half squares designate chromosome boundaries.

**Supplementary Fig. 2 | BUSCO genome assembly results.** Visualized using the generate_plot.py script in BUSCO. Text in bars correspond to the full BUSCO scores of each species, with proportion of BUSCOs on the x-axis and species on the y-axis.

**Supplementary Fig. 3 | $K_S$ plots of paralogous and orthologous gene pairs.** For each focal taxon, $K_S$ plots of all paralogous gene pairs (solid lines) and orthologous gene pairs (dotted line). Colors of lines correspond to species as indicated by key.

**Supplementary Fig. 4 | Orthologous and paralogous $K_S$ plot inferences.** Inferences made from $K_S$ ortholog/paralog plots (Supplementary Fig. 3) where WGD were considered shared if the paralogous peak from the focal taxon had a higher $K_S$ than the orthologous peaks. WGD were considered unique/more recent if the paralogous peak from the focal taxon had a lower $K_S$ than the orthologous peaks. Duplications were considered to be small-scale, if $K_S$ peaks exhibited substantially lower gene duplicate densities.

**Supplementary Fig. 5 | Rate-adjusted $K_S$ plots of paralogous and orthologous gene pairs.** For each focal taxon, a mixed plot of paralogs and orthologs, adjusted for substitution rates, produced by *ksrates*. The gray histogram represents the distribution of anchor gene pairs with the likely WGD peaks identified through lognormal mixture model clustering, shown in blue, red, and green. The vertical dashed lines marked 'a', 'b', and 'c' correspond to the peak positions of these components and are used as estimates for WGD timing. Vertical long-dashed lines mark adjusted peak estimates of ortholog distributions between the focal taxon and all other taxa assessed. These are interpreted as speciation events and are color-coded and numbered to match the speciation events shown in the legend, with colored boxes around each line indicating one standard deviation above and below the mean estimate. Horizontal arrows at the bottom illustrate how substitution rate corrections by *ksrates* shift the inferred timing of these speciation events. Additionally, the phylogram produced by *ksrates* for each focal taxon is provided. Branch lengths reflect distances calculated from ortholog $K_S$ distributions.

**Supplementary Fig. 6 | Estimated WGD placements from *ksrates*.** WGD inferences based on identified WGD peaks and speciation events from *ksrates* (Supplementary Fig. 5).

**Supplementary Fig. 7 | Gene blocks retained across all chromosomes from WGDI**. Teal boxes to the right indicate the *Vitis* (vvi161s) chromosomes used to generate subgenomes. Boxes around gene blocks within the plot showcase the gene blocks used specifically for that chromosome's subgenome. Colors follow the subgenome numbers specified in Supplementary Fig. 10: Subgenome 1: blue, 2: green, 3: yellow, 4: pink, 5: orange, 6: purple.

**Supplementary Fig. 8 | Subgenome collinearity, phylogeny, and gene fractionation results from chromosome 2 of *Vitis***. Generated using WGDI. **a)** The top portion corresponds to the subgenome dotplots of each species against *Vitis*. Each dot in the dotplot corresponds to a syntenic gene included in the subgenome and is colored according to the color legend of Fig 2e. Specific subgenome orders are noted with circled numbers above the dots in the plot. **b)** Subgenome phylogenetic relationships are indicated with the phylogeny on the bottom left. Values at each node show local posterior probability (LPP) values. **c)** Gene fractionation results of each subgenome, following the color schemes of **a**. The x-axis corresponds to the basepair number, while the y-axis corresponds to the proportion of retained genes in homoeologous regions of *Vitis* chromosome 2.

**Supplementary Fig. 9 | Subgenome collinearity, phylogeny, and gene fractionation results from chromosome 7 of *Vitis***. Generated using WGDI. **a)** The top portion corresponds to the subgenome dotplots of each species against *Vitis*. Each dot in the dotplot corresponds to a syntenic gene included in the subgenome and is colored according to the color legend of Fig 2e. Specific subgenome orders are noted with circled numbers above the dots in the plot. **b)** Subgenome phylogenetic relationships are indicated with the phylogeny on the bottom left. Values at each node show local posterior probability (LPP) values. **c)** Gene fractionation results of each subgenome, following the color schemes of **a**. The x-axis corresponds to the basepair number, while the y-axis corresponds to the proportion of retained genes in homoeologous regions of *Vitis* chromosome 7.

**Supplementary Fig. 10 | Subgenome collinearity, phylogeny, and gene fractionation results from chromosome 11 of *Vitis***. Generated using WGDI. **a)** The top portion corresponds to the subgenome dotplots of each species against *Vitis*. Each dot in the dotplot corresponds to a syntenic gene included in the subgenome and is colored according to the color legend of Fig 2e. Specific subgenome orders are noted with circled numbers above the dots in the plot. **b)** Subgenome phylogenetic relationships are indicated with the phylogeny on the bottom left. Values at each node show local posterior probability (LPP) values. **c)** Gene fractionation results of each subgenome, following the color schemes of **a**. The x-axis corresponds to the basepair number, while the y-axis corresponds to the proportion of retained genes in homoeologous regions of *Vitis* chromosome 11. Circled and blue numbers on phylogeny of chromosome 11 correspond to the branches where accelerated molecular evolution (AR, see Methods) were estimated.

**Supplementary Fig. 11 | Subgenome collinearity, phylogeny, and gene fractionation results from chromosome 14 of *Vitis***. Generated using WGDI. **a)** The top portion corresponds to the subgenome dotplots of each species against *Vitis*. Each dot in the dotplot corresponds to a syntenic gene included in the subgenome and is colored according to the color legend of Fig 2e. Specific subgenome orders are noted with circled numbers above the dots in the plot. **b)** Subgenome phylogenetic relationships are indicated with the phylogeny on the bottom left. Values at each node show local posterior probability (LPP) values. **c)** Gene fractionation results of each subgenome, following the color schemes of **a**. The x-axis corresponds to the basepair number, while the y-axis corresponds to the proportion of retained genes in homoeologous regions of *Vitis* chromosome 14.

**Supplementary Fig. 12 | Subgenome collinearity, phylogeny, and gene fractionation results from chromosome 15 of *Vitis***. Generated using WGDI. **a)** The top portion corresponds to the subgenome dotplots of each species against *Vitis*. Each dot in the dotplot corresponds to a syntenic gene included in the subgenome and is colored according to the color legend of Fig

2e. Specific subgenome orders are noted with circled numbers above the dots in the plot. **b)** Subgenome phylogenetic relationships are indicated with the phylogeny on the bottom left. Values at each node show local posterior probability (LPP) values. **c)** Gene fractionation results of each subgenome, following the color schemes of **a**. The x-axis corresponds to the basepair number, while the y-axis corresponds to the proportion of retained genes in homoeologous regions of *Vitis* chromosome 15.

**Supplementary Fig. 13 | Subgenome gene composition of each Asterales species within each tested chromosome (chromosomes 2, 7, 11, 14, and 15).** Numbers in stacked bars represent the percentage of genes retained for that designated subgenome. Actual numbers are found in the table below the plots.

**Supplementary Fig. 14 | Results of the WHALE analysis.** Utilizing the critical (left) and relaxed (right) models. The tested WGD event locations are represented by diamonds, with each color representing an analysis: blue is based solely off primary literature (analysis 5 from Supplementary Fig. 22), orange is from *ksrates* (analysis 8 from Supplementary Fig. 22), and green includes all tested locations (analysis 9 from Supplementary Fig. 22). The diamond fill determines whether the model was significant with the critical (dark gray) or relaxed (light gray) model. White fill indicates that the result was not significant.

**Supplementary Fig. 15 | Gene expansions and contractions across Asterales.** CAFE5 tree showing total gene expansions and contractions (top, black font) and significant gene expansions and contractions (bottom, red font) at nodes and tips. Node-associated numbers are shown in black circles.

**Supplementary Fig. 16 | Plot of GO terms indicating most expanded, contracted, lost, and lineage-specific genes.** Biological process GO terms enriched among contracted, expanded, lost, and lineage-specific gene families in *Acicarpha*, Asteraceae, and their shared clade (*Acicarpha* + Asteraceae), clustered based on semantic similarity. Word clouds summarize each GO cluster using keywords, with font size proportional to the enrichment significance.

**Supplementary Fig. 17 | Composition of each duplication type.** As indicated by McScanX, by taxonomic grouping and analysis.

**Supplementary Fig. 18 | *FAD2* gene family expansion in Asteraceae.** Tips are colored by family, where Vitaceae are purple, Brassicaceae are blue, Campanulaceae are dark red, Menyanthaceae are orange, Goodeniaceae are yellow, Calyceraceae are green, and Asteraceae are red. Genes are classified by their duplication type as indicated in the legend.

**Supplementary Fig. 19 | *CYP71B23* gene family expansion in Asteraceae.** Tips are colored by family, where Vitaceae are purple, Brassicaceae are blue, Campanulaceae are dark red, Menyanthaceae are orange, Goodeniaceae are yellow, Calyceraceae are green, and Asteraceae are red. Genes are classified by their duplication type as indicated in the legend.

**Supplementary Fig. 20 | GO enrichment plot of most positively selected genes in Asteraceae.** GO enrichment of biological processes associated with genes experiencing site-specific positive selection in Asteraceae.

**Supplementary Fig. 21 | *FLC*-like gene tree.** Shaded regions show Asterales-specific lineages. Tips are colored by family as indicated in the legend.

**Supplementary Fig. 22 | Calibration points for WHALE analysis.** Locations of input calibrations for WHALE on a phylogeny of the eight Asterales taxa. Colored boxes to the left indicate whether that analysis (top) included the calibration (right). Analysis 5 is based on primary literature, 8 is from *ksrates*, and 9 is a combination of both. White boxes indicate that the calibration was not included.

**Supplementary Fig. 23 | Long-read sequence data quality scores.** Pauvre plots showing quality phred reports of raw, long-read sequence data of **a)** *Acicarpha*, **b)** *Scaevola*, and **c)** *Platycodon*. Individual plots represent a single flow cell.

**Supplementary Fig. 24 | $K_S$ dotplots of the eight Asterales taxa against chromosome 11 of *Vitis*.** Dotplots of each Asterales genome (x-axis) against chromosome 11 subgenomes of *Vitis* (or Grape, y-axis) generated with WGDI. Dots are colored by their individual $K_S$ value, as indicated by the scale to the right of each block.

**Supplementary Fig. 25 | Time-calibrated phylogeny of the eight Asterales taxa plus *Arabidopsis* and *Vitis*.** Generated with TreePL. Values at nodes are in millions of years. Pentagons at some nodes indicate locations of fossil calibrations used to estimate geologic age.

**Supplementary Fig. 26 | Proteome quality assessment of *Acicarpha*, *Scaevola*, and *Platycodon*.** Generated with OMArk. Bottom portion is showing the consistency percentage of the proteome, while the top portion is indicating the percentage of missing, duplicated, and single-copy conserved genes. Numerical representation of this figure can be found in Supplementary Table 10.

**Supplementary Fig. 27 | Plastome of *Platycodon grandiflorus*.** Bandage assembly graph and visualization of annotated plastome of *Platycodon grandiflorus*.

**Supplementary Fig. 28 | Plastome of *Nymphoides indica*.** Bandage assembly graph and visualization of annotated plastome of *Nymphoides indica*.

**Supplementary Fig. 29 | Plastome of *Scaevola aemula*.** Bandage assembly graph and visualization of annotated plastome of *Scaevola aemula*.

**Supplementary Fig. 30 | Plastome of *Acicarpha procumbens*.** Bandage assembly graph and visualization of annotated plastome of *Acicarpha procumbens*.

**Supplementary Fig. 31 | Plastome of *Arctium lappa*.** Bandage assembly graph and visualization of annotated plastome of *Arctium lappa*.

**Supplementary Fig. 32 | Plastome of *Lactuca sativa*.** Bandage assembly graph and visualization of annotated plastome of *Lactuca sativa*.

**Supplementary Fig. 33 | Plastome of *Chrysanthemum lavandulifolium*.** Bandage assembly graph and visualization of annotated plastome of *Chrysanthemum lavandulifolium*.

**Supplementary Fig. 34 | Plastome of *Helianthus annuus*.** Bandage assembly graph and visualization of annotated plastome of *Helianthus annuus*.

**Supplementary Fig. 35 | Plastome alignments indicating rearrangements.** Collinearity of Asterales plastomes using Mauve progressive alignment.

**Supplementary Fig. 36 | Genome composition plots of each Asterales taxon.** K-mer distribution and fitted models produced from GenomeScope 2.0. Genome composition inferred from Smudgeplot, where each haplotype structure is visualized by a smudge and the heat of each smudge indicates the relative frequency of each haplotype structure in the genome.

**Supplementary Fig. 37 | Species tree of eight target taxa within Asterales plus *Arabidopsis* and *Vitis*.** Generated with OrthoFinder. Numbers at nodes indicate branch support (top, posterior probability) and the number of well-supported gene duplication events shared among common ancestors (bottom). Numbers at the tip labels correspond to unique gene duplications for that taxon.

**Supplementary Fig. 38 | Accelerated subgenome evolution in Asterales relative to chromosome 11 of *Vitis***. Distribution of accelerated regions (ARs) specific to **a)** Asteraceae + *Acicarpha* subgenomes and **b)** the *Acicarpha*-specific duplication event. Regions of high AR density (AR hotspots) are numbered. **c)** and **d)** Within each hotspot, the distribution of ARs are clustered around specific genes (green, orange), which represent candidate targets for the accelerated evolution.
