## Supplementary Figures 1-38 for "Gene duplication dynamics and regulatory evolution shape the diversification of Asteraceae"

SupplementaryFig1.pdf

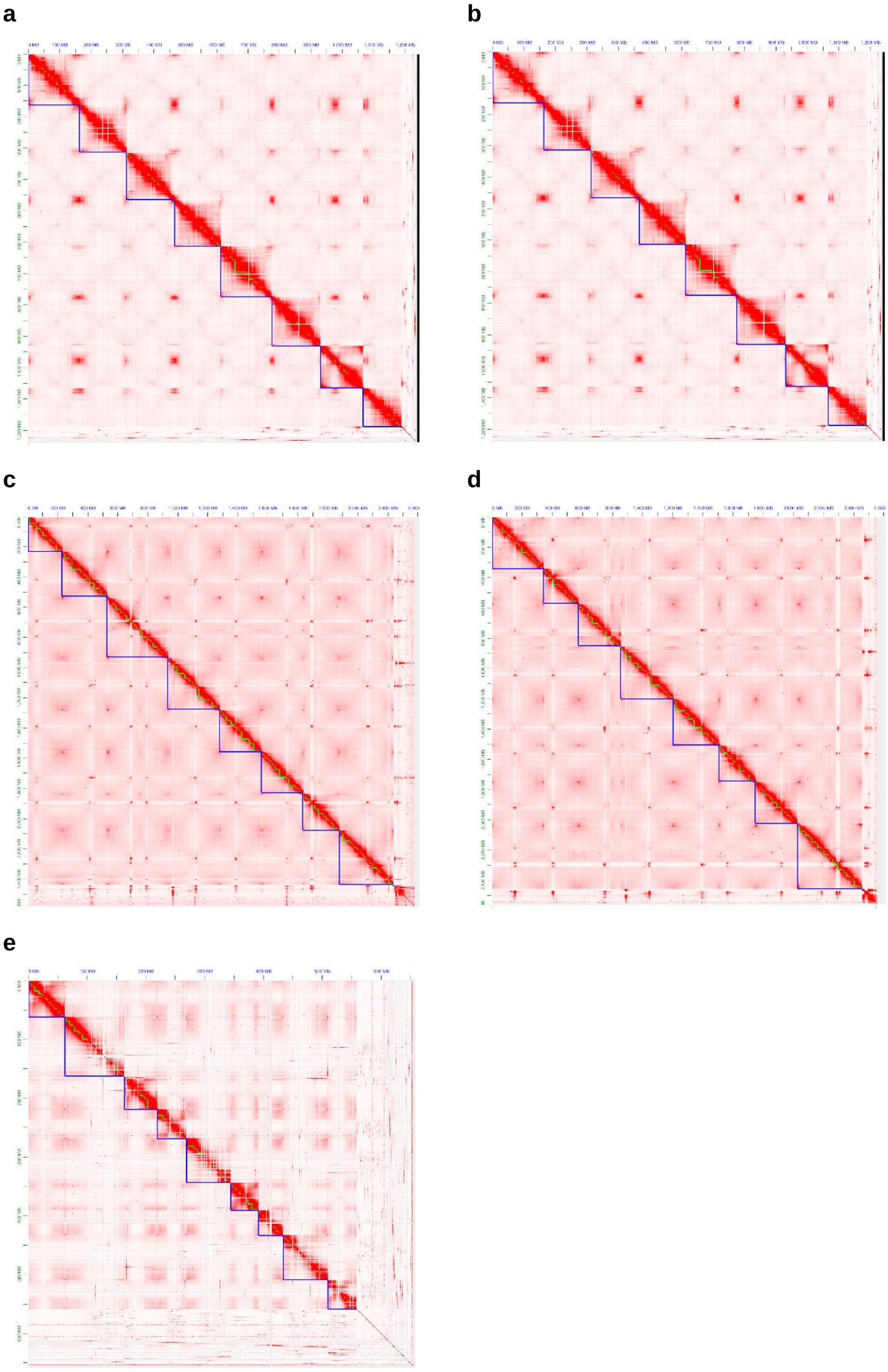

SupplementaryFig2.pdf

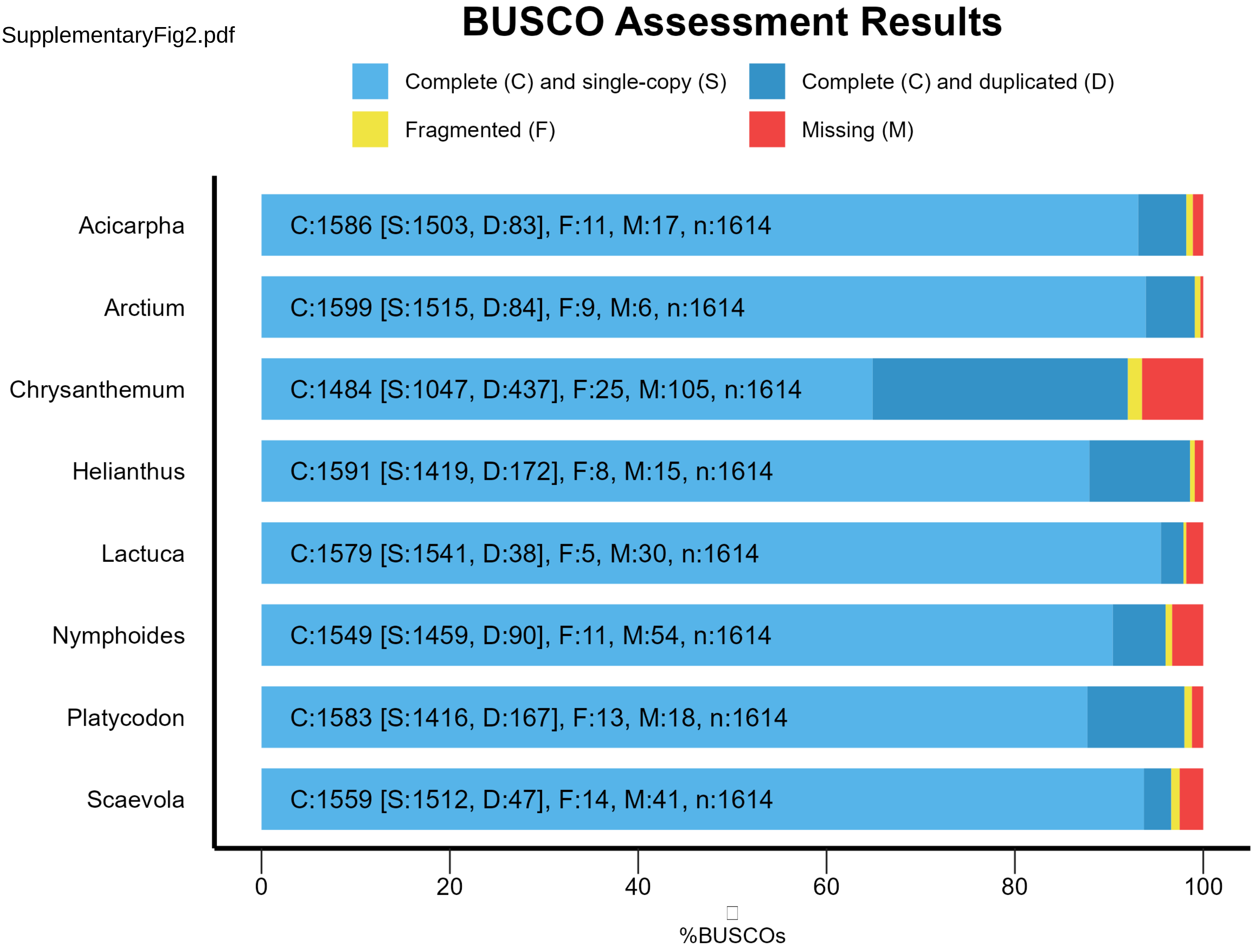

SupplementaryFig3.pdf

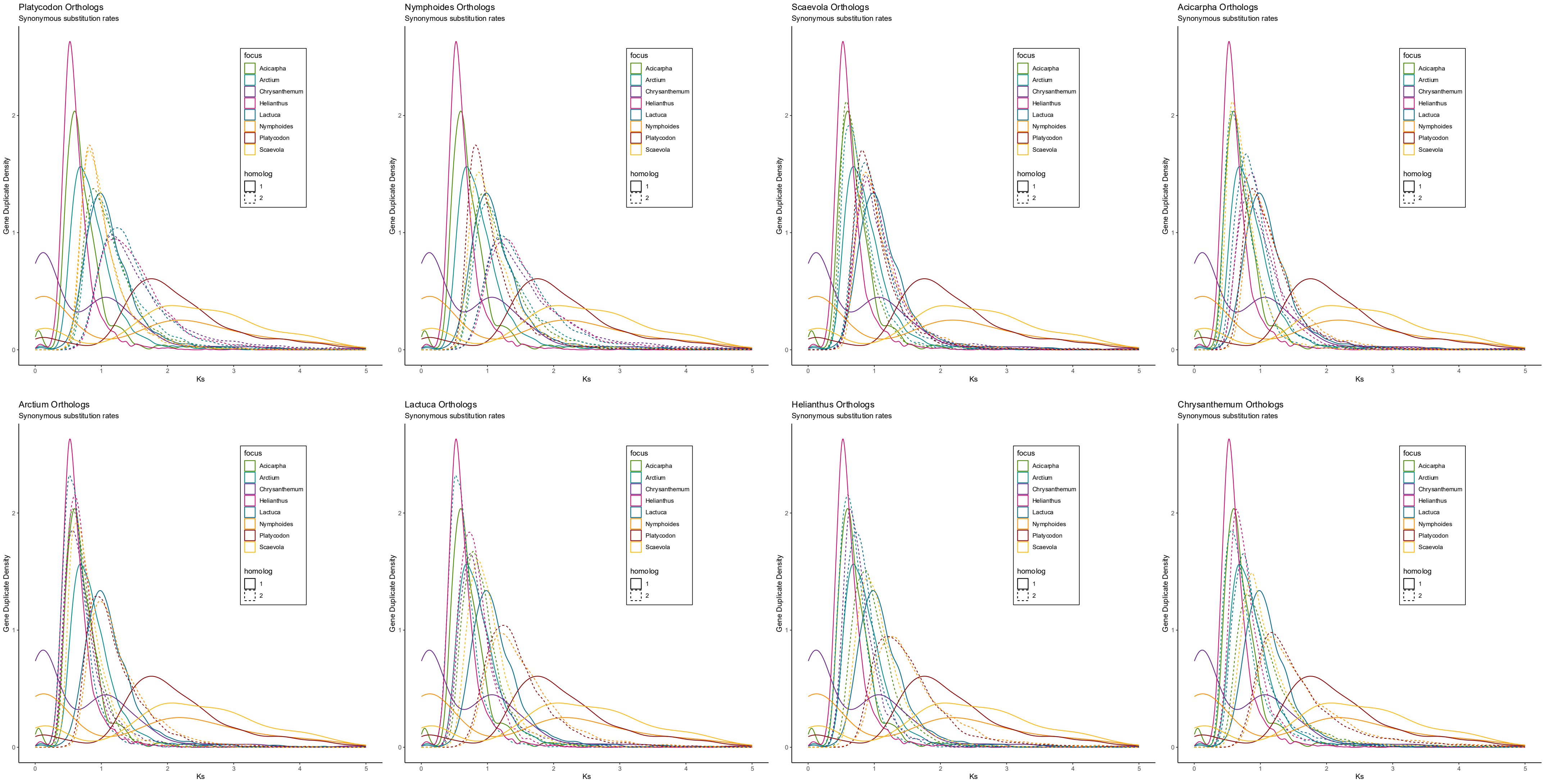

SupplementaryFig4.pdf

### WGD and small-scale duplication inferences from Ks distributions

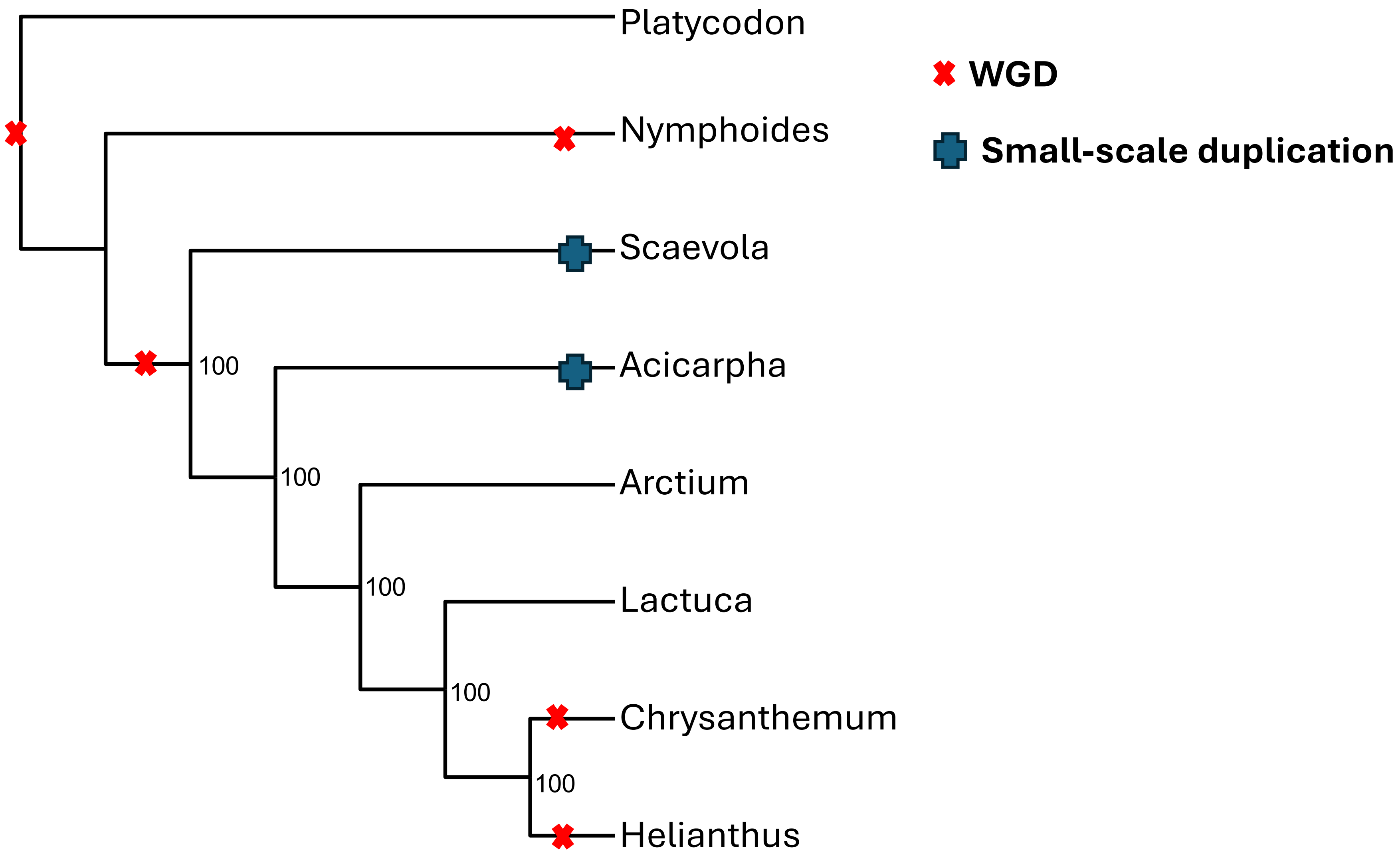

SupplementaryFig5.pdf

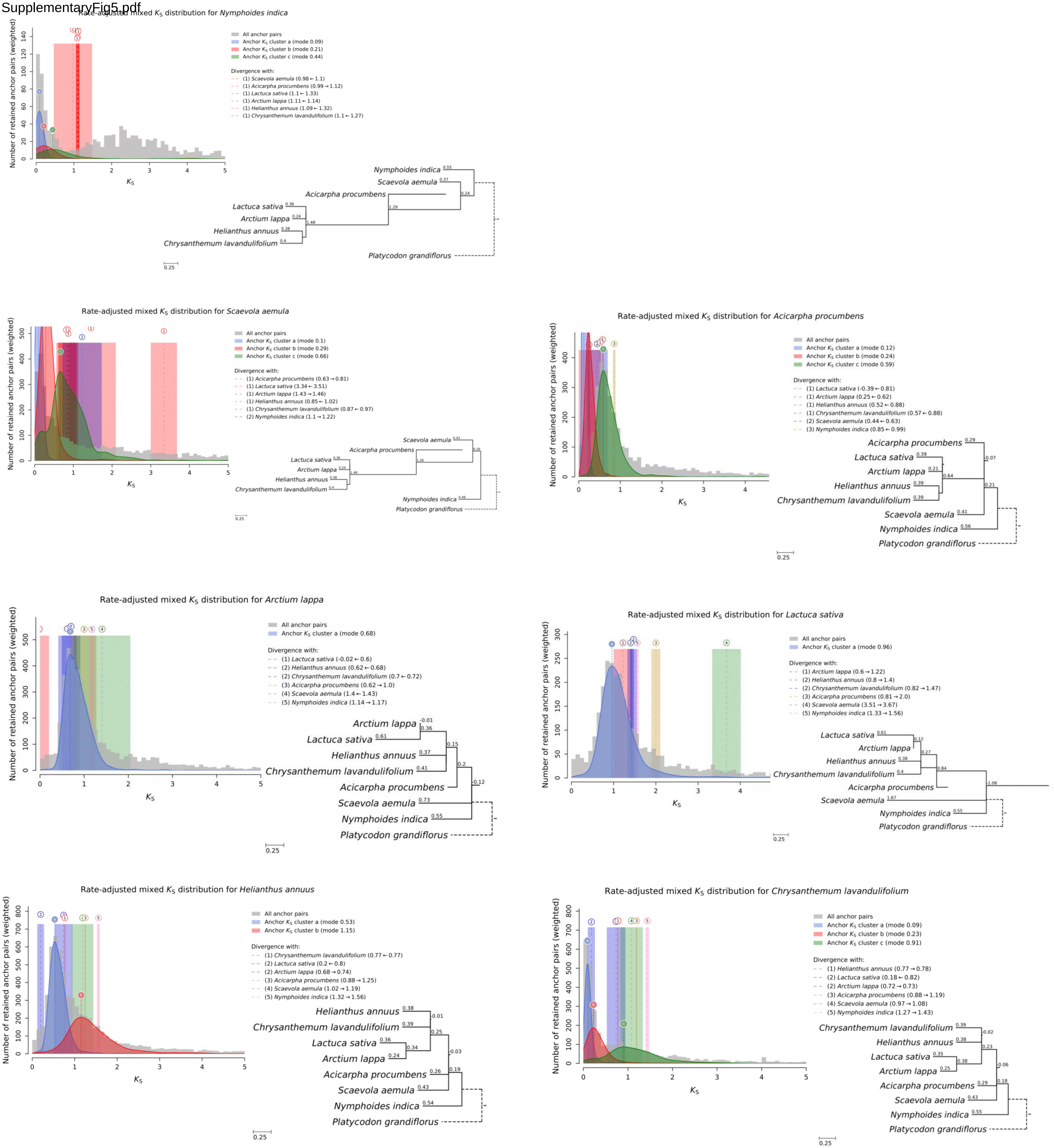

SupplementaryFig6.pdf

#### WGD inferences from *ksrates* results

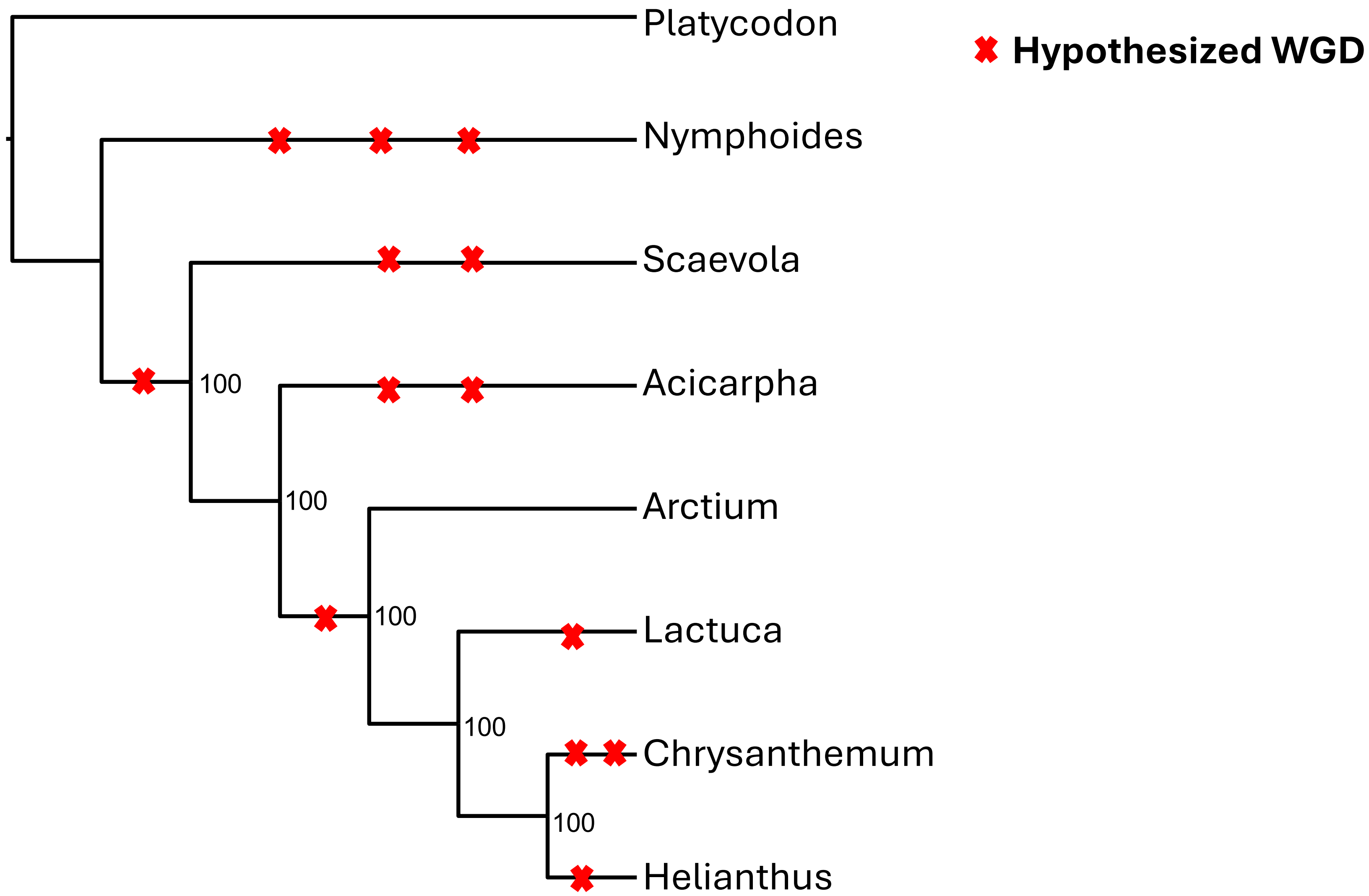

SupplementaryFig7.pdf

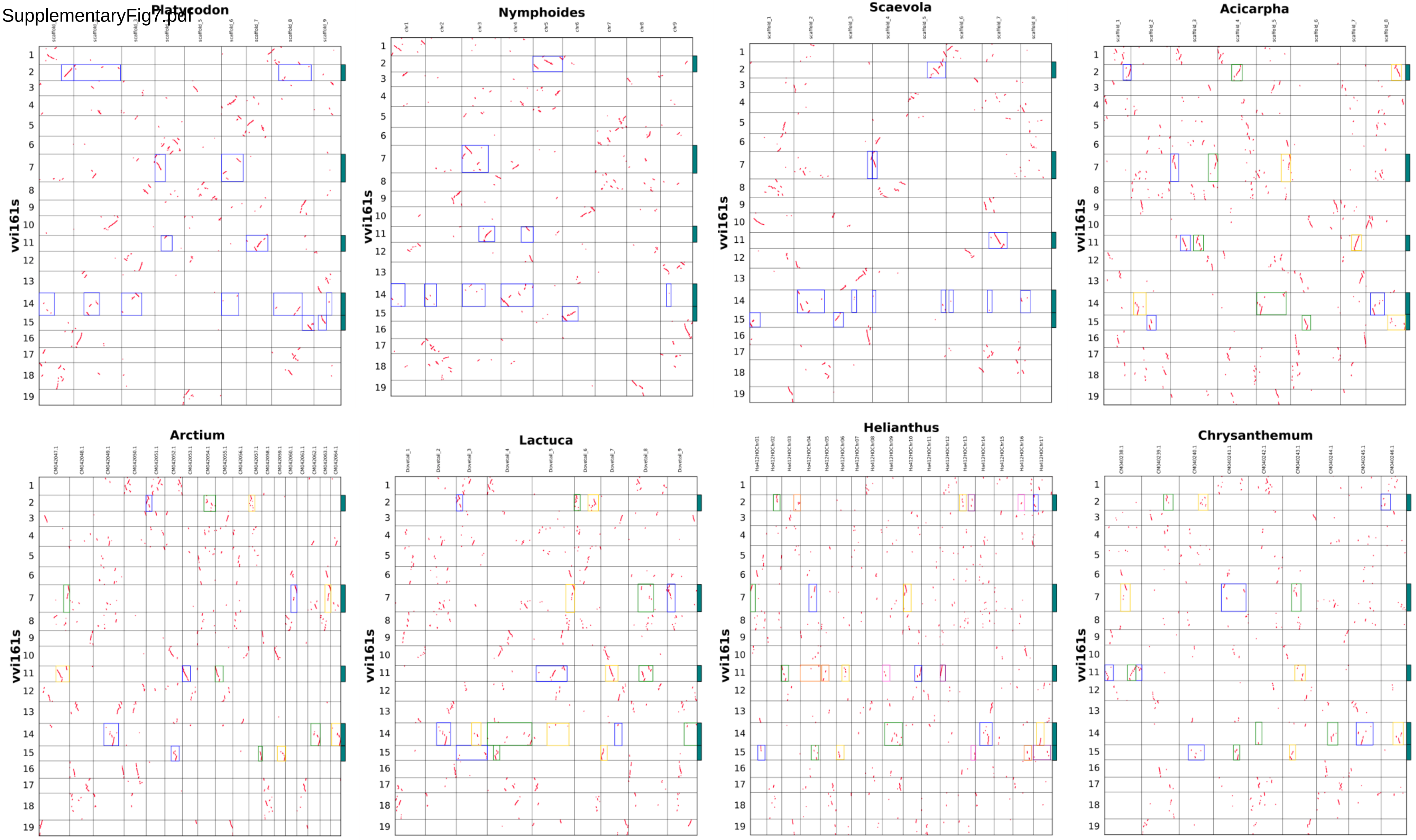

SupplementaryFig8.pdf

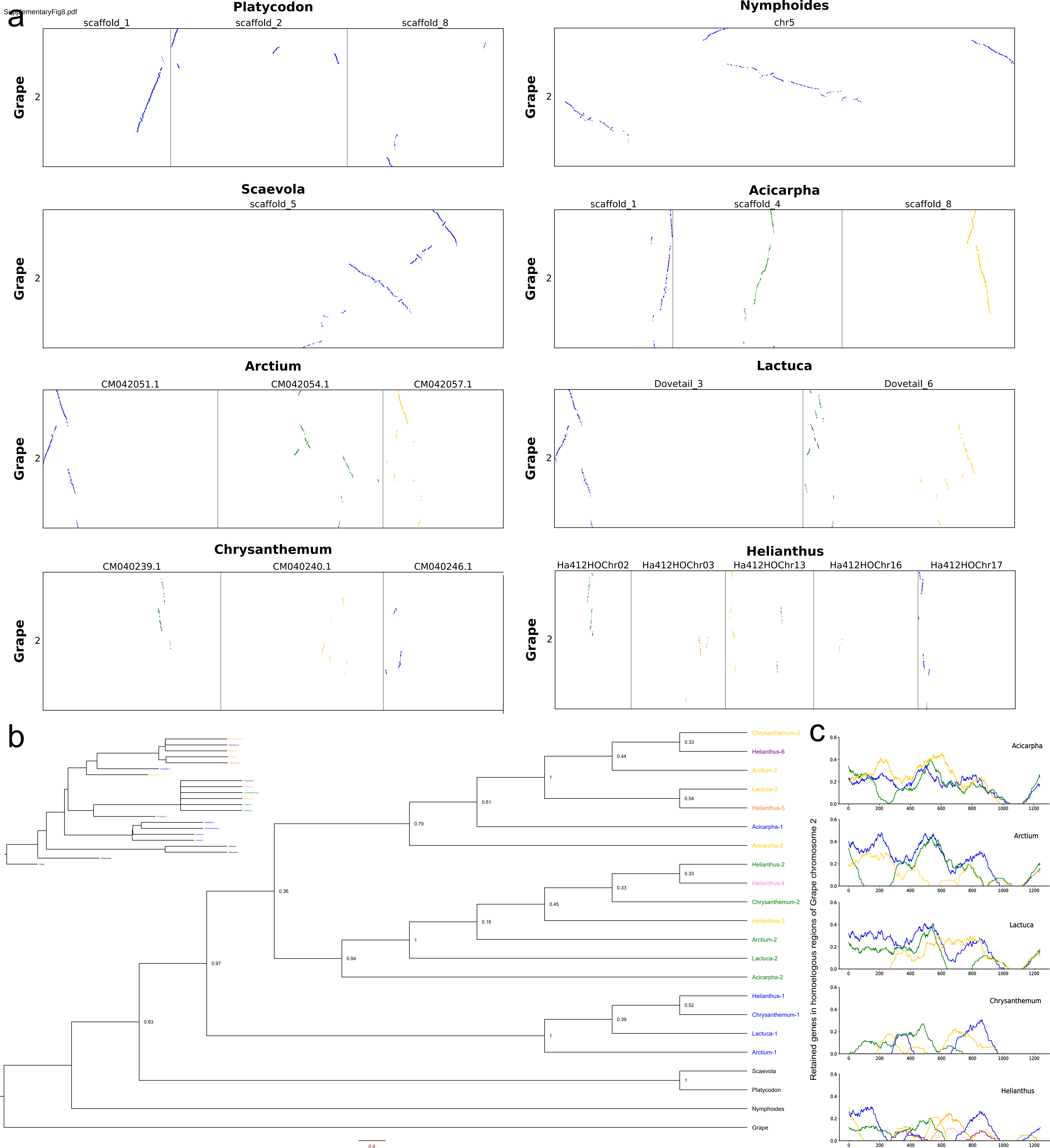

SupplementaryFig9.pdf

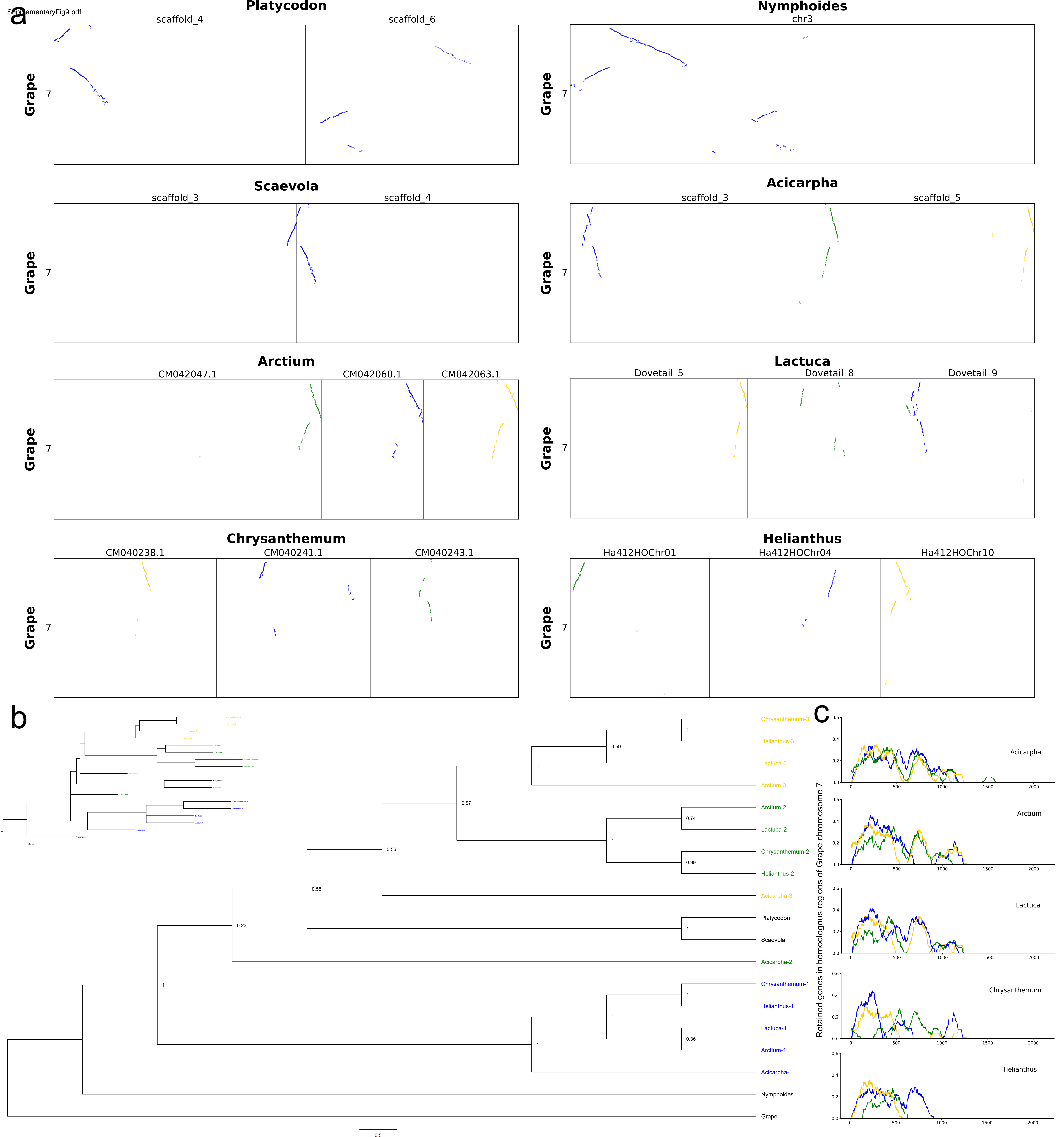

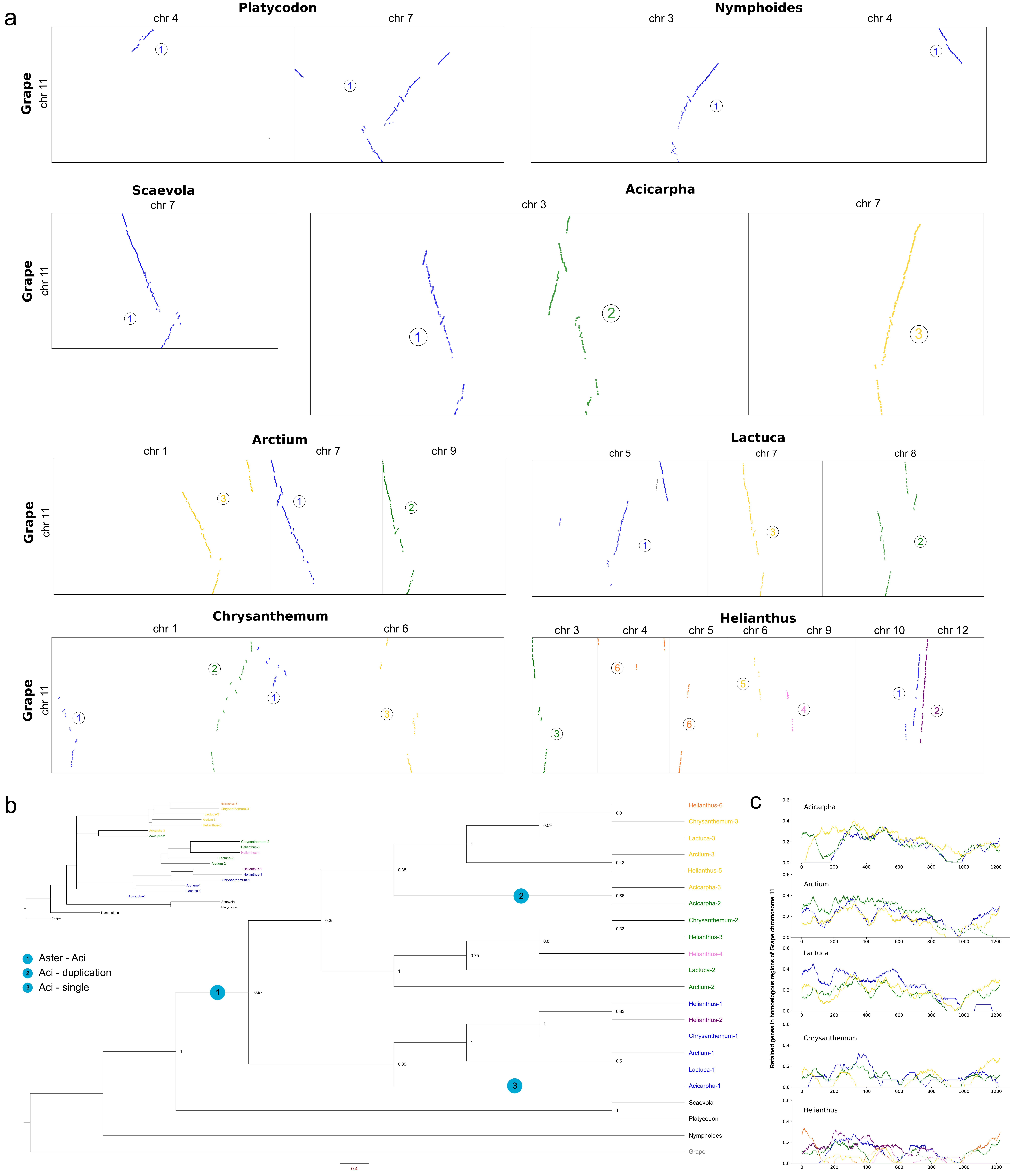

a
Platycodon
chr 4
chr 7
Nymphoides
chr 3
chr 4
Grape
chr 11
Scaevola
chr 7
Acicarpha
chr 3
chr 7
Arctium
chr 1
chr 7
chr 9
Lactuca
chr 5
chr 7
chr 8
Chrysanthemum
chr 1
chr 6
Helianthus
chr 3
chr 4
chr 5
chr 6
chr 9
chr 10
chr 12
b
1 Aster - Aci
2 Aci - duplication
3 Aci - single
Helianthus-6
Chrysanthemum-3
Lactuca-3
Arctium-3
Helianthus-5
Acicarpha-3
Acicarpha-2
Chrysanthemum-2
Helianthus-3
Helianthus-4
Lactuca-2
Arctium-2
Helianthus-1
Helianthus-2
Chrysanthemum-1
Arctium-1
Lactuca-1
Acicarpha-1
Scaevola
Platycodon
Nymphoides
Grape
0.8
0.59
1
0.43
0.35
0.86
0.35
0.33
0.8
0.75
1
0.97
0.83
1
1
0.5
0.39
1
1
0.4
c
Acicarpha
Arctium
Lactuca
Chrysanthemum
Helianthus
Retained genes in homoeologous regions of Grape chromosome 11
0.0
0.2
0.4
0.6
0
200
400
600
800
1000
1200

SupplementaryFig11.pdf

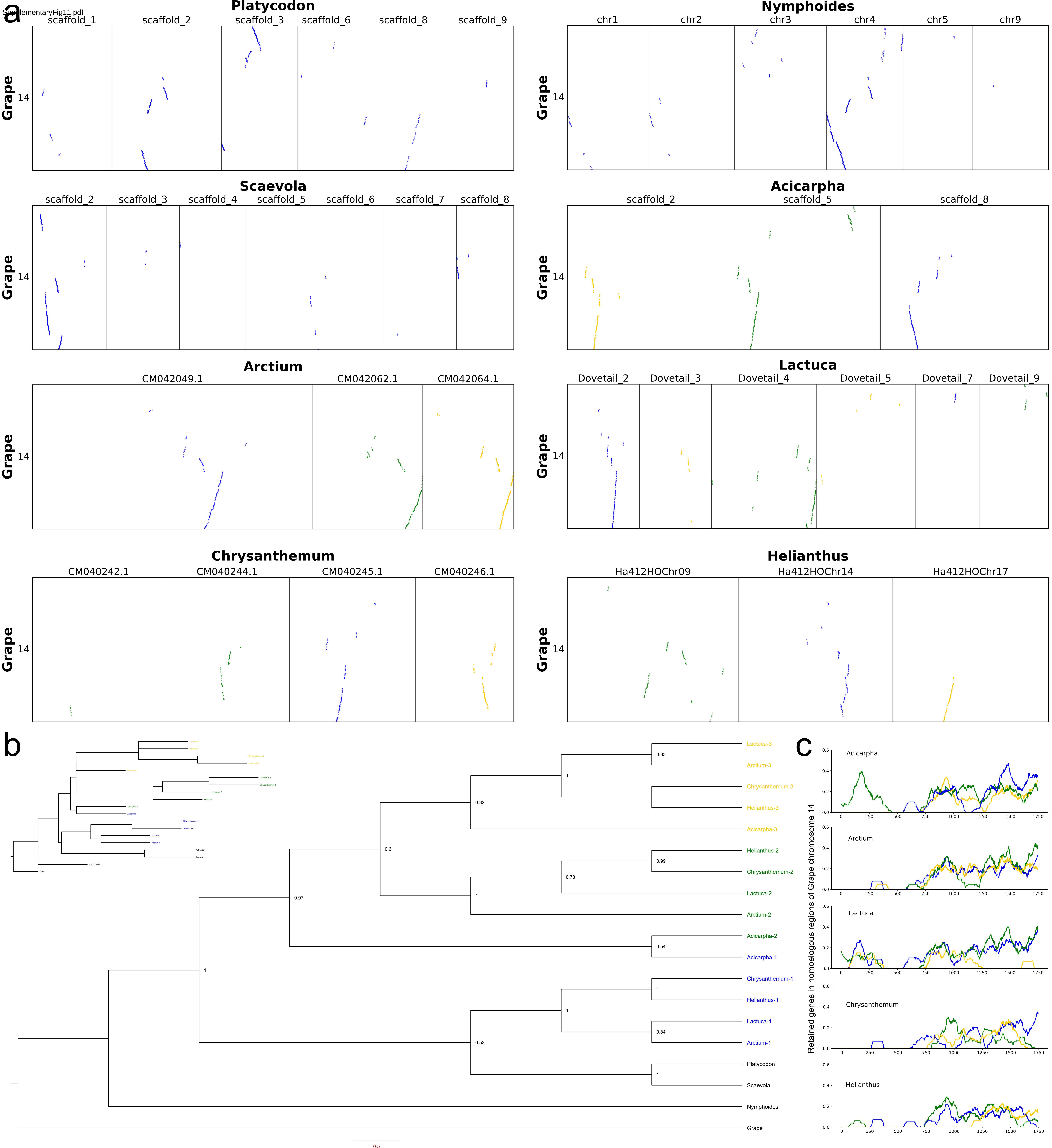

SupplementaryFig12.pdf

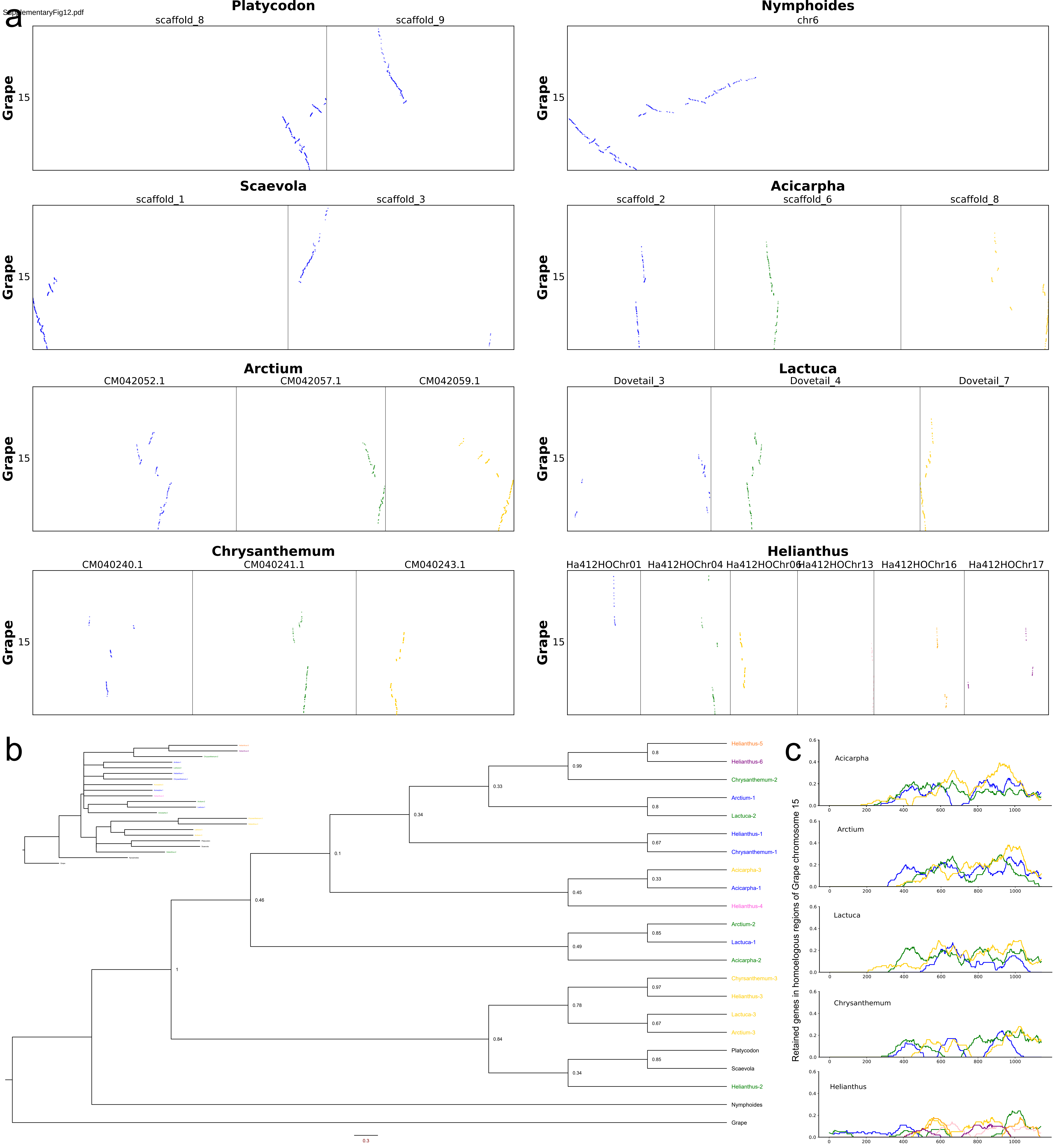

SupplementaryFig13.pdf

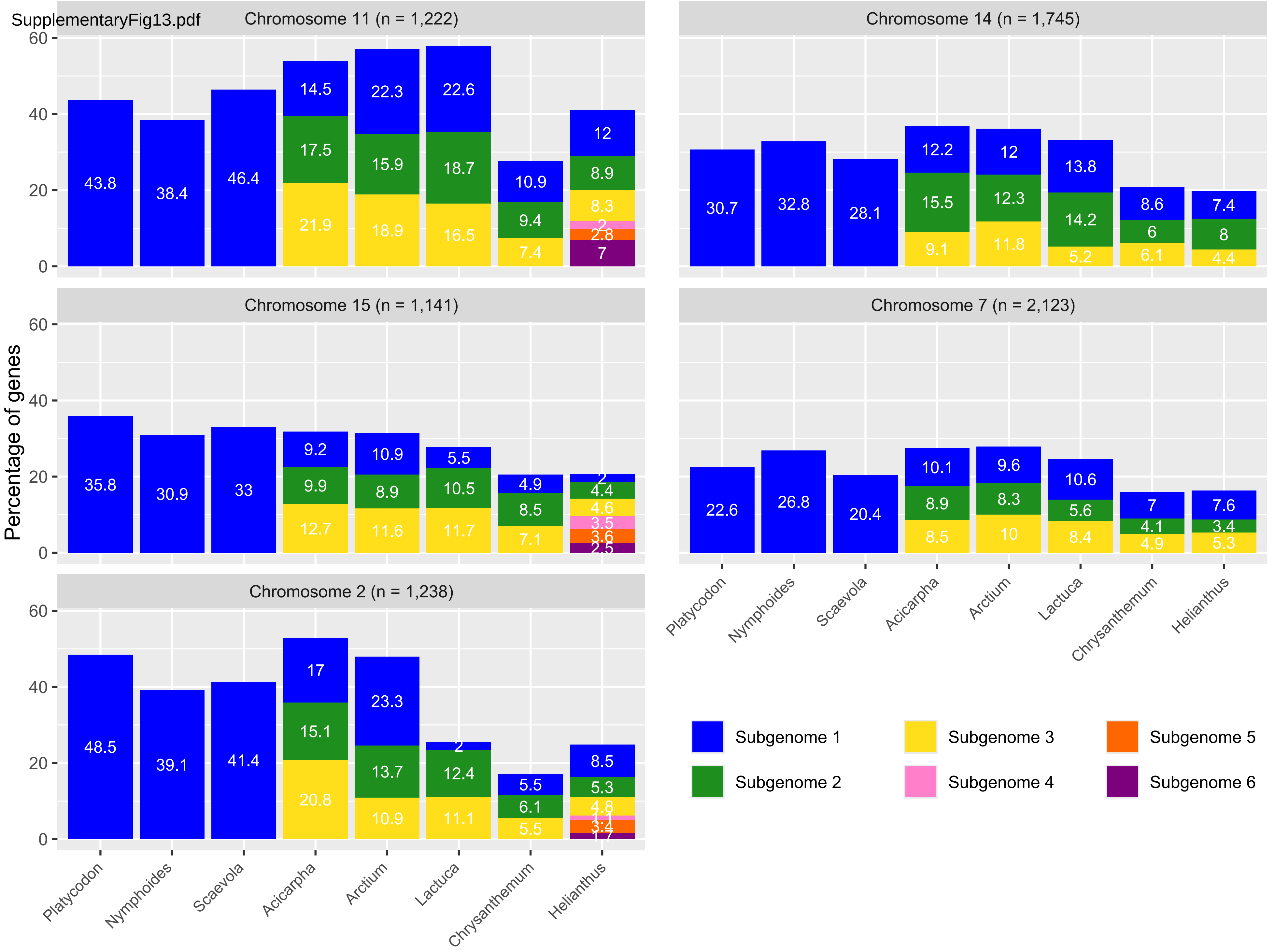


| | Subgenome | ***Platycodon*** | ***Nymphoides*** | ***Scaevola*** | ***Acicarpha*** | ***Arctium*** | ***Lactuca*** | ***Chrysanthemum*** | ***Helianthus*** |
|---|---|---|---|---|---|---|---|---|---|
| ***Chromosome 11*** *(n = 1,222)* | *1* | 535 | 469 | 567 | 177 | 273 | 276 | 133 | 147 |
| | *2* | --- | --- | --- | 214 | 194 | 228 | 115 | 109 |
| | *3* | --- | --- | --- | 268 | 231 | 202 | 90 | 101 |
| | *4* | --- | --- | --- | --- | --- | --- | --- | 24 |
| | *5* | --- | --- | --- | --- | --- | --- | --- | 34 |
| | *6* | --- | --- | --- | --- | --- | --- | --- | 85 |
| ***Chromosome 14*** *(n = 1,745)* | *1* | 535 | 572 | 491 | 213 | 210 | 240 | 150 | 130 |
| | *2* | --- | --- | --- | 271 | 214 | 248 | 104 | 139 |
| | *3* | --- | --- | --- | 159 | 206 | 90 | 106 | 77 |
| ***Chromosome 15*** *(n = 1,141)* | *1* | 409 | 353 | 377 | 105 | 124 | 63 | 56 | 23 |
| | *2* | --- | --- | --- | 113 | 102 | 120 | 97 | 50 |
| | *3* | --- | --- | --- | 145 | 132 | 133 | 81 | 52 |
| | *4* | --- | --- | --- | --- | --- | --- | --- | 40 |
| | *5* | --- | --- | --- | --- | --- | --- | --- | 41 |
| | *6* | --- | --- | --- | --- | --- | --- | --- | 29 |
| ***Chromosome 2*** *(n = 1,238)* | *1* | 600 | 484 | 512 | 210 | 288 | 25 | 68 | 105 |
| | *2* | --- | --- | --- | 187 | 169 | 153 | 75 | 66 |
| | *3* | --- | --- | --- | 257 | 135 | 138 | 68 | 60 |
| | *4* | --- | --- | --- | --- | --- | --- | --- | 14 |
| | *5* | --- | --- | --- | --- | --- | --- | --- | 42 |
| | *6* | --- | --- | --- | --- | --- | --- | --- | 21 |
| ***Chromosome 7*** *(n = 2,123)* | *1* | 480 | 569 | 434 | 215 | 203 | 226 | 148 | 162 |
| | *2* | --- | --- | --- | 188 | 177 | 118 | 87 | 73 |
| | *3* | --- | --- | --- | 180 | 213 | 178 | 104 | 113 |

SupplementaryFig14.pdf

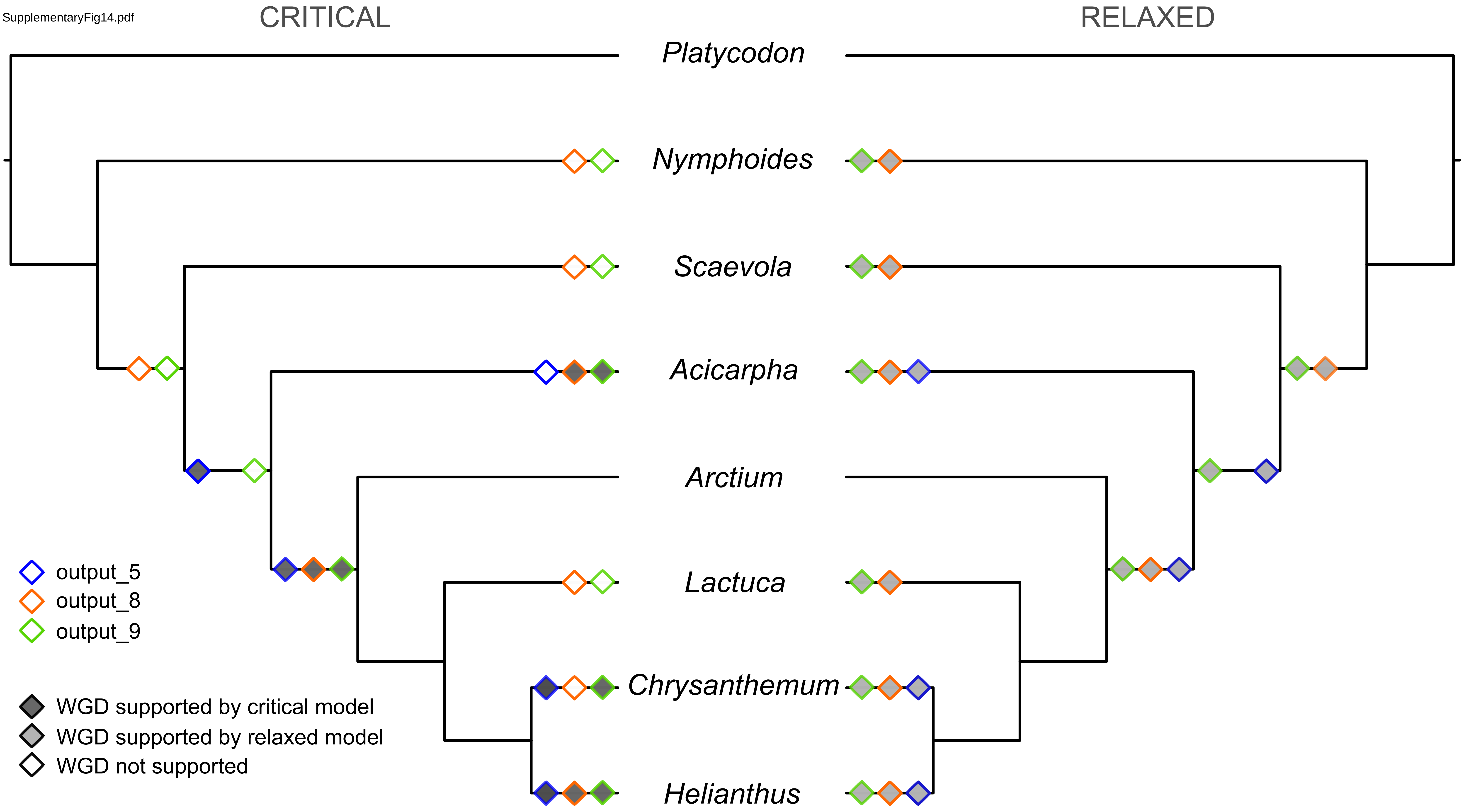

SupplementaryFig15.pdf

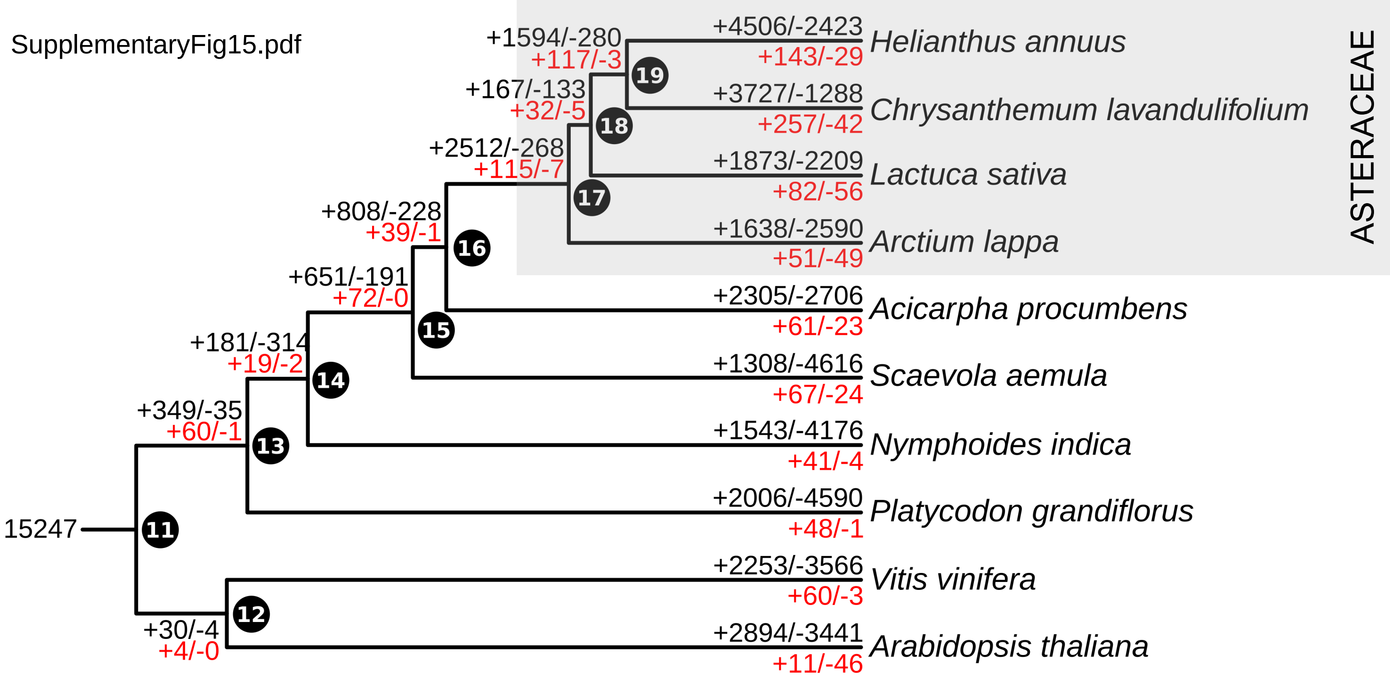

Supplementary Fig16.pdf

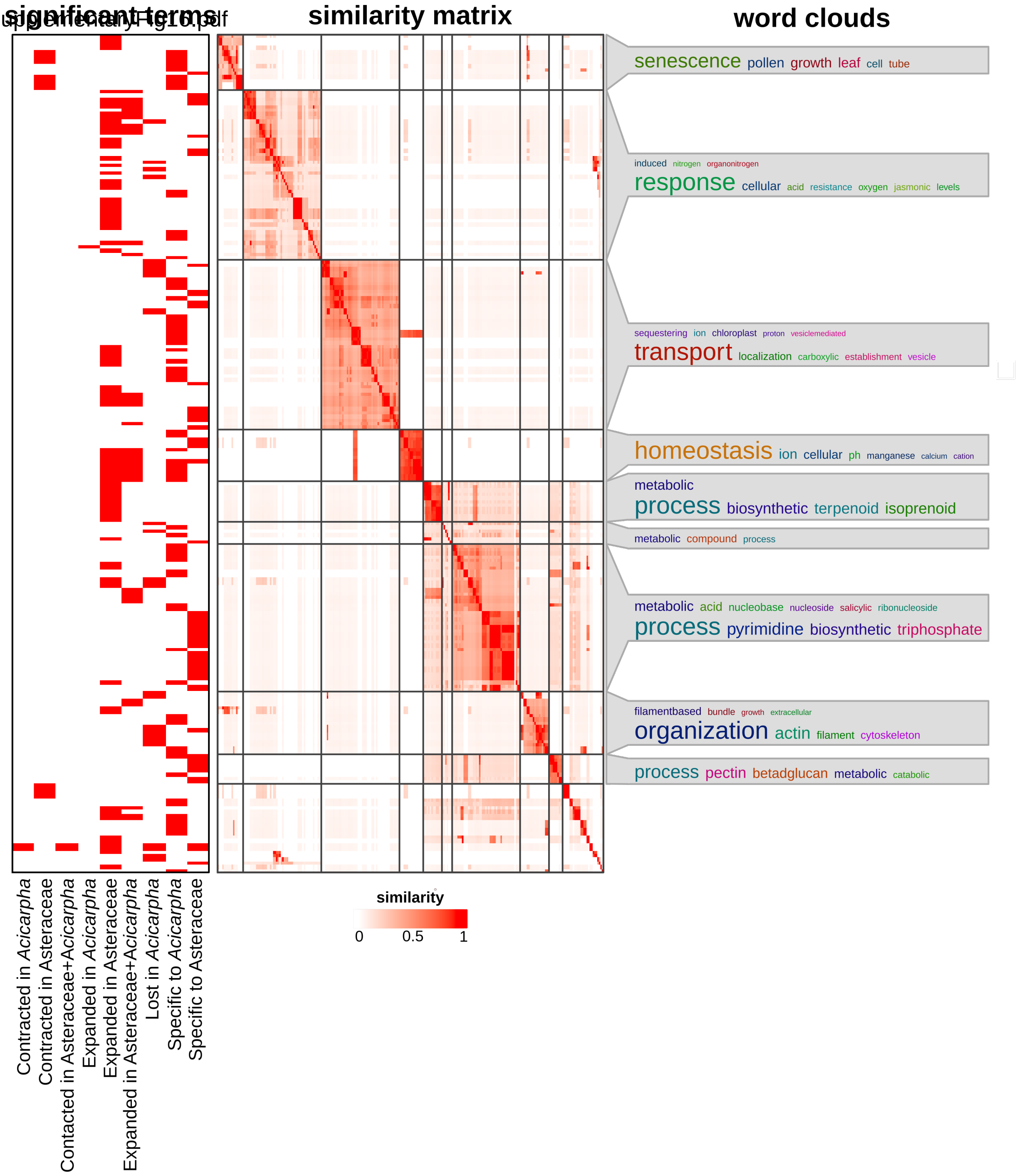

SupplementaryFig17.pdf

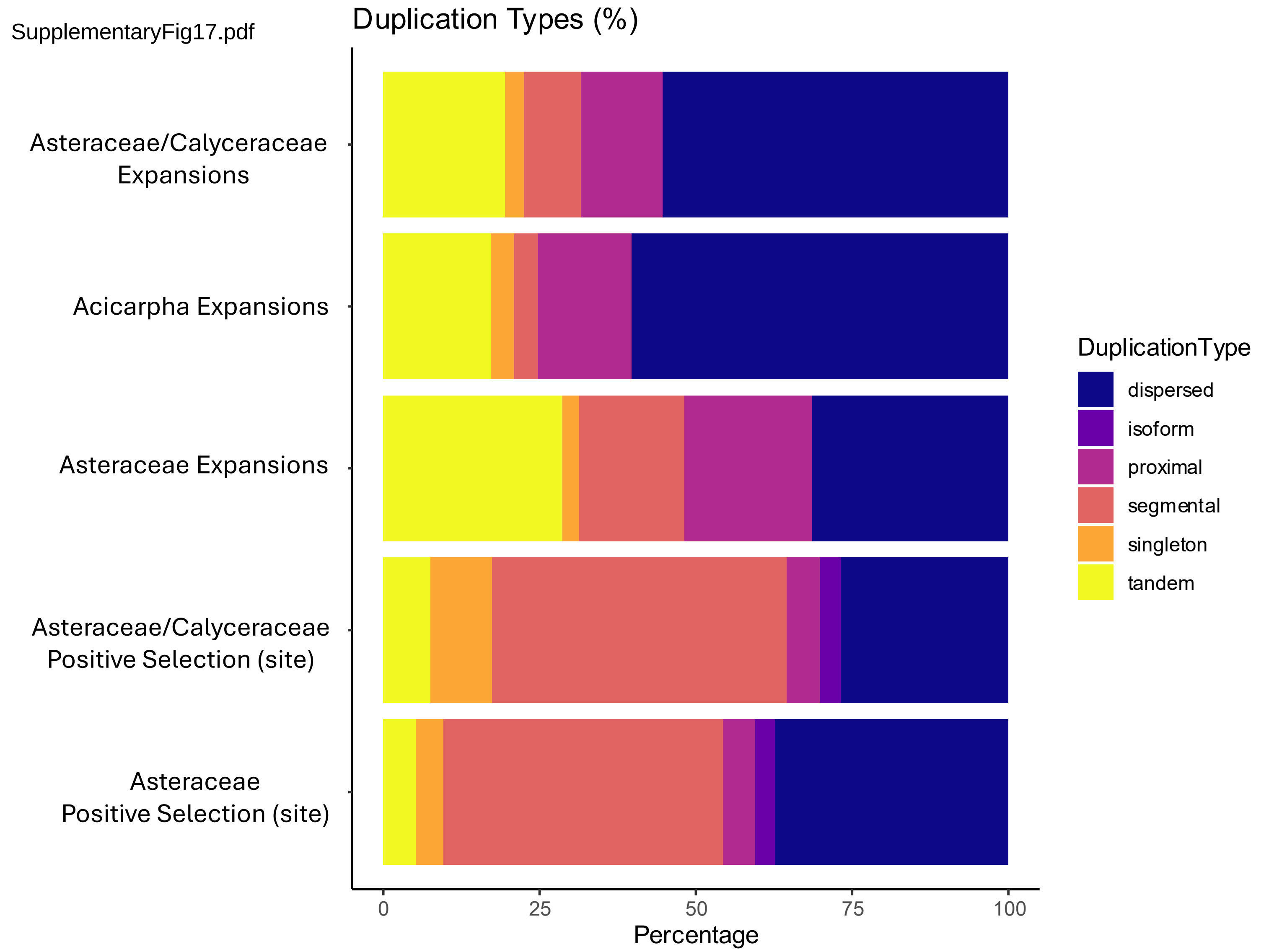

Orthogroup OG00000057 FAD2

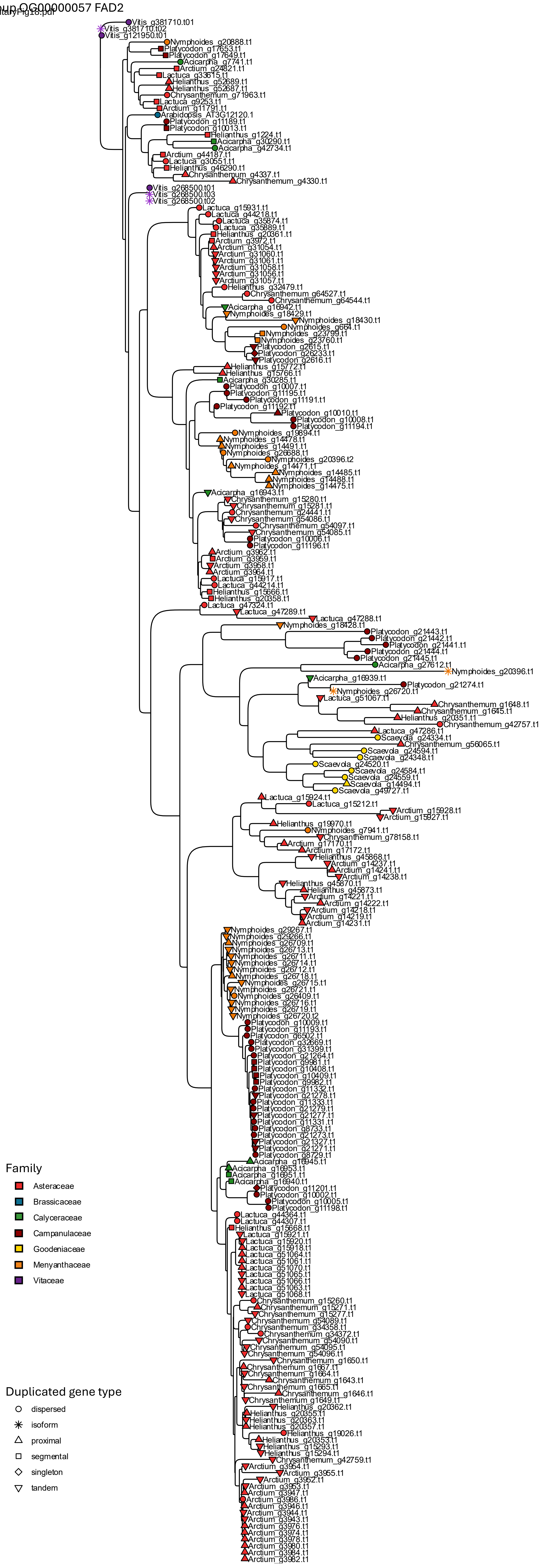

Orthogroup OG00000077 CYP1B

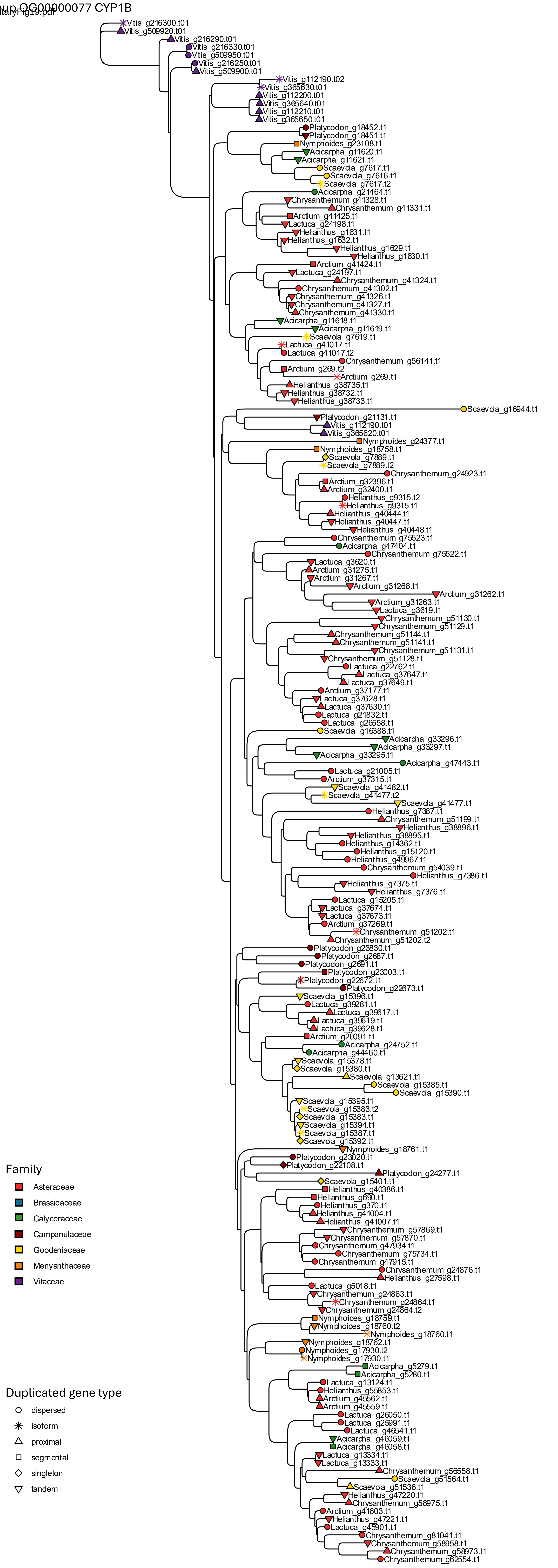

SupplementaryFig20.pdf

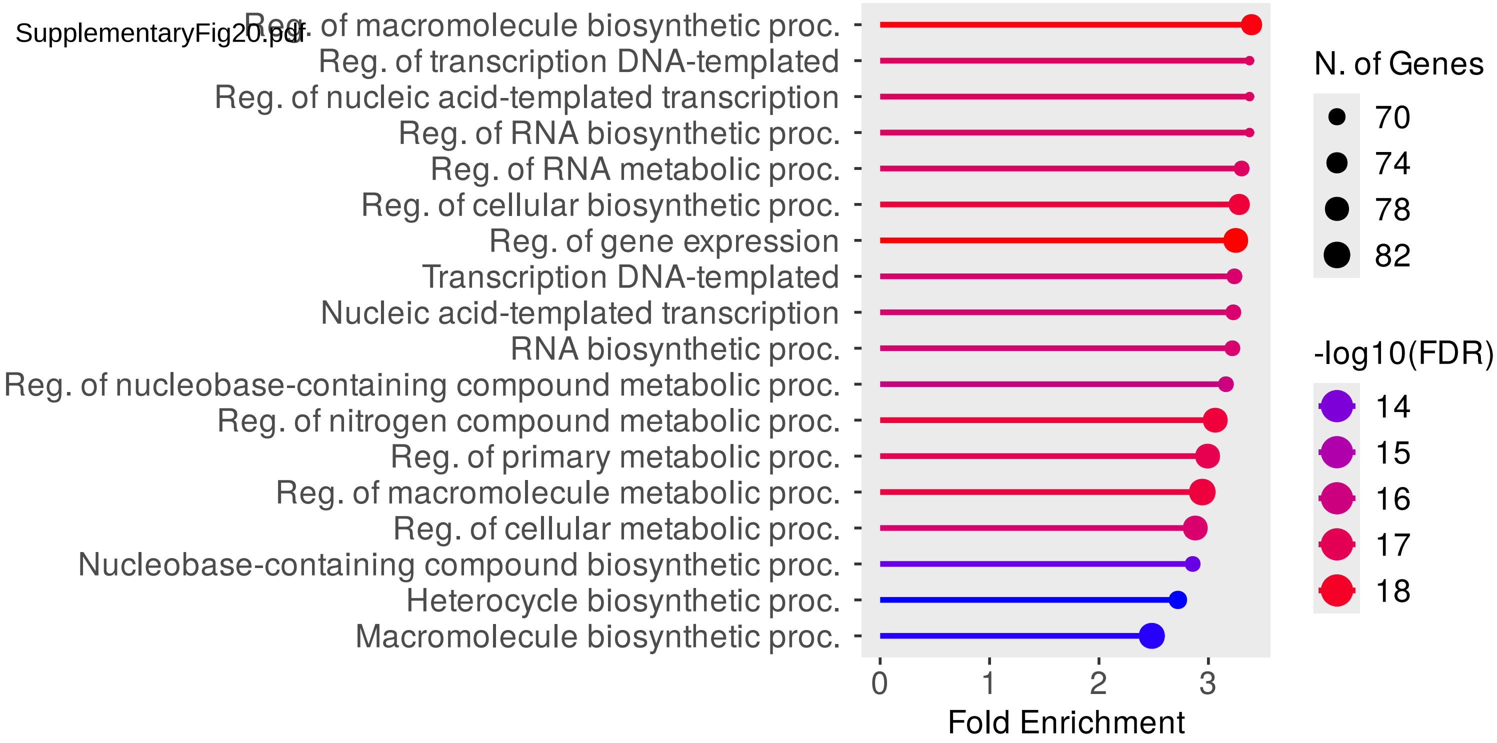

SupplementaryFig21.pdf

FLC-like extended tree

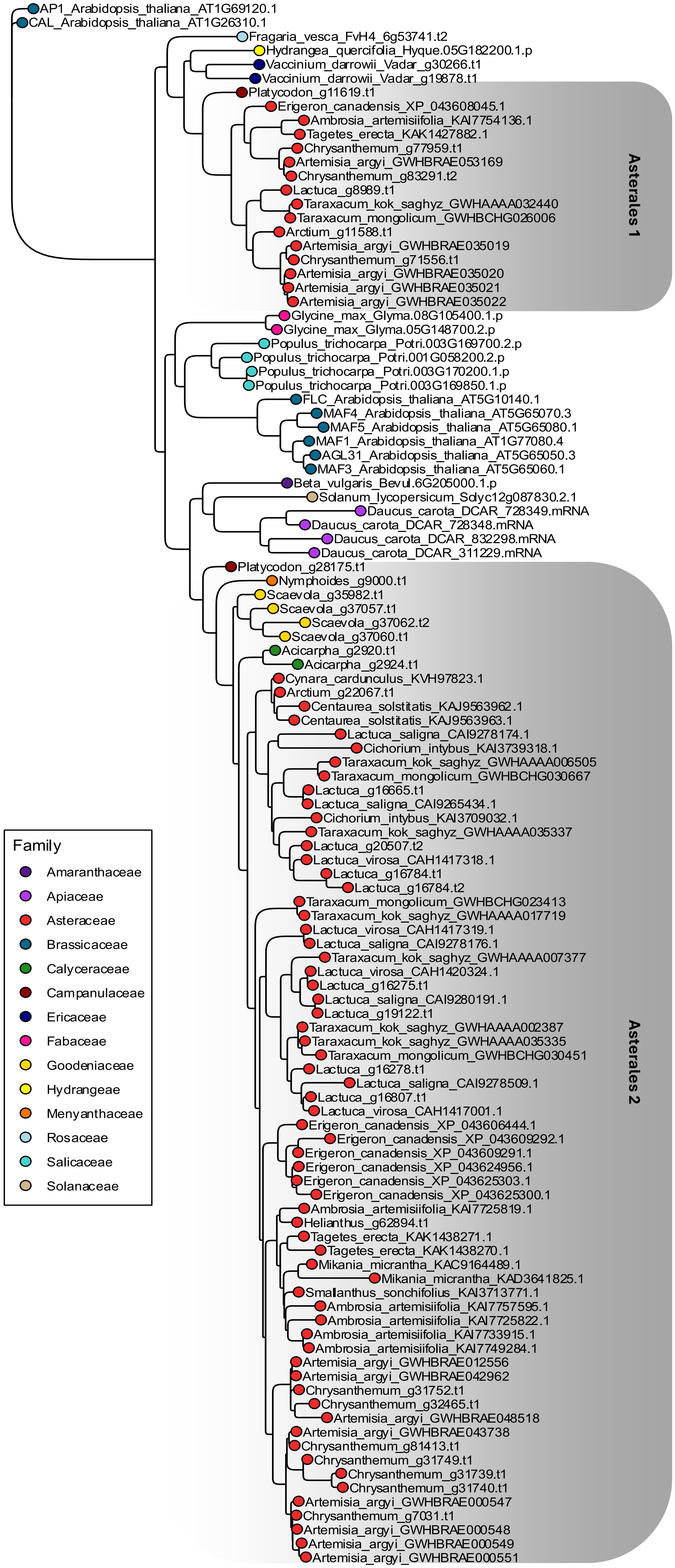

SupplementaryFig21.pdf

#### Orthogroup OG00000806 FLC

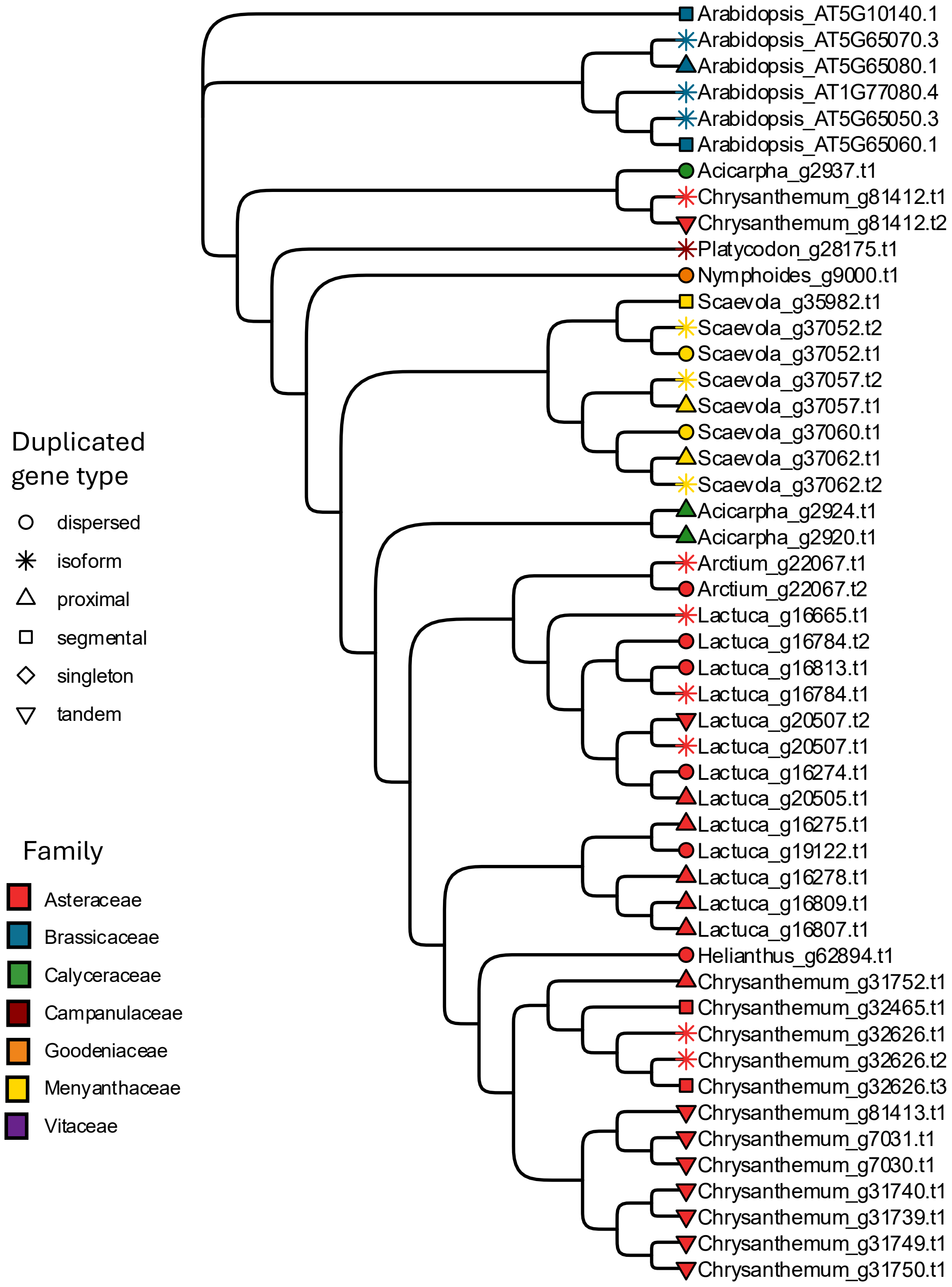

SupplementaryFig22.pdf

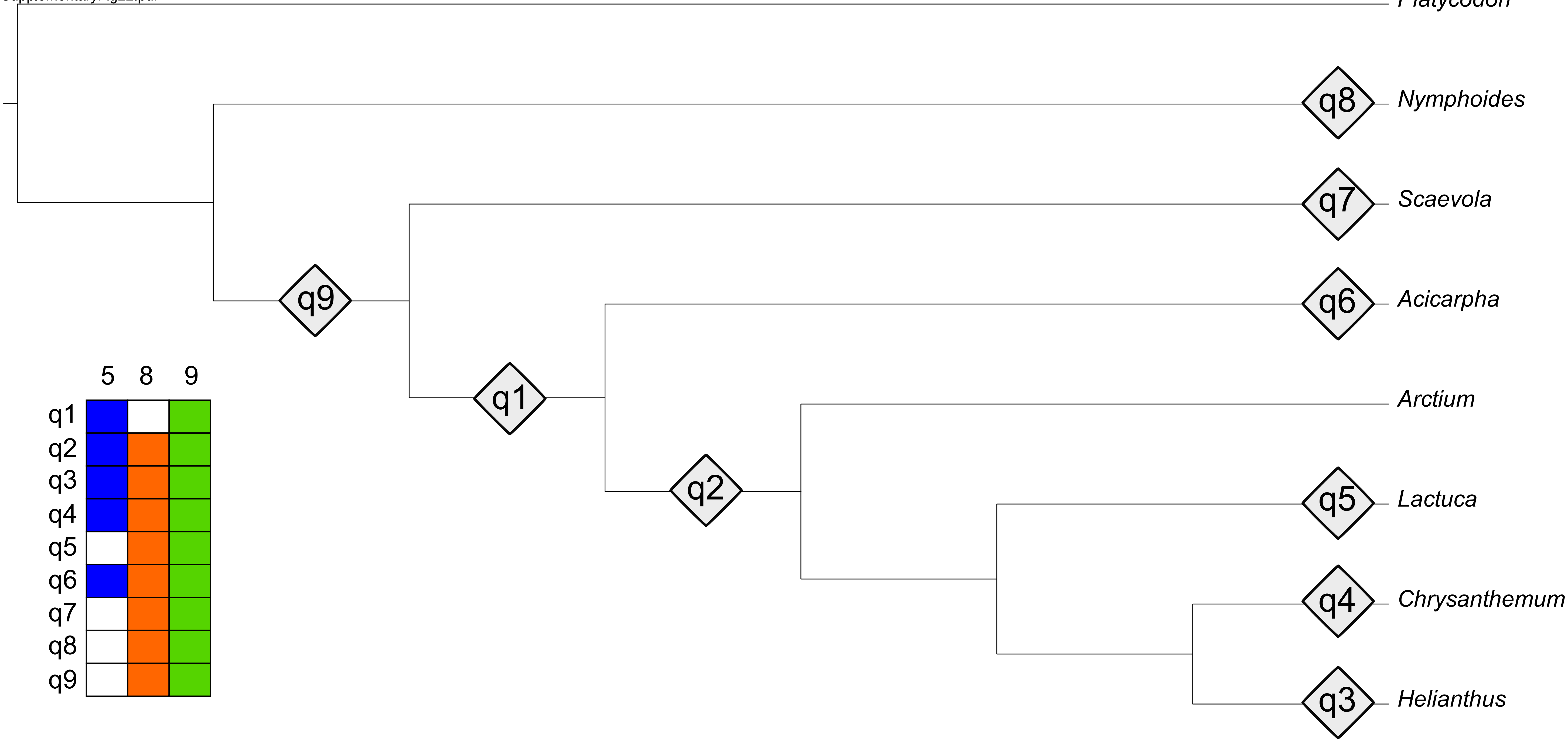

SupplementaryFig23.pdf

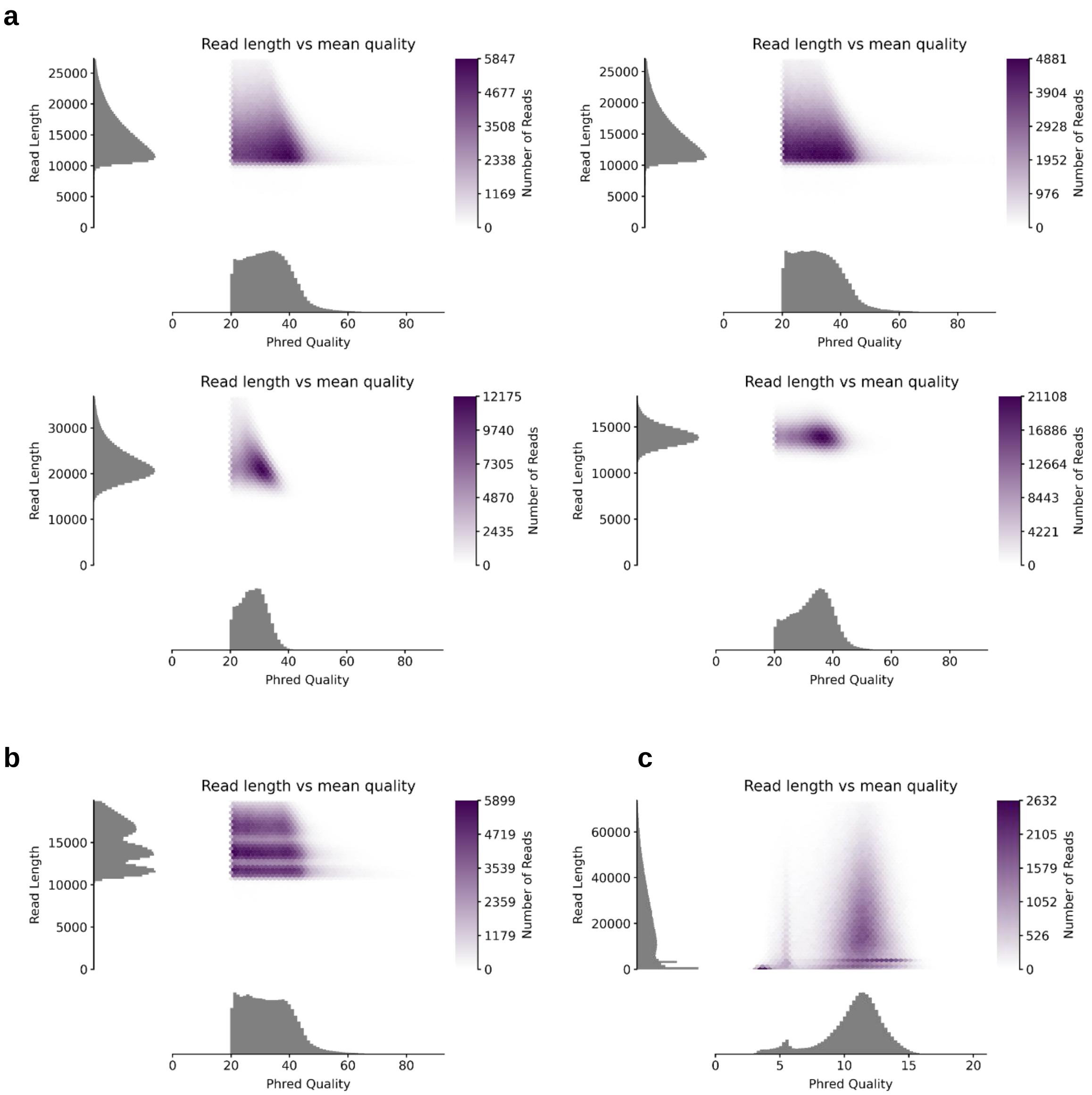

SupplementaryFig24.pdf

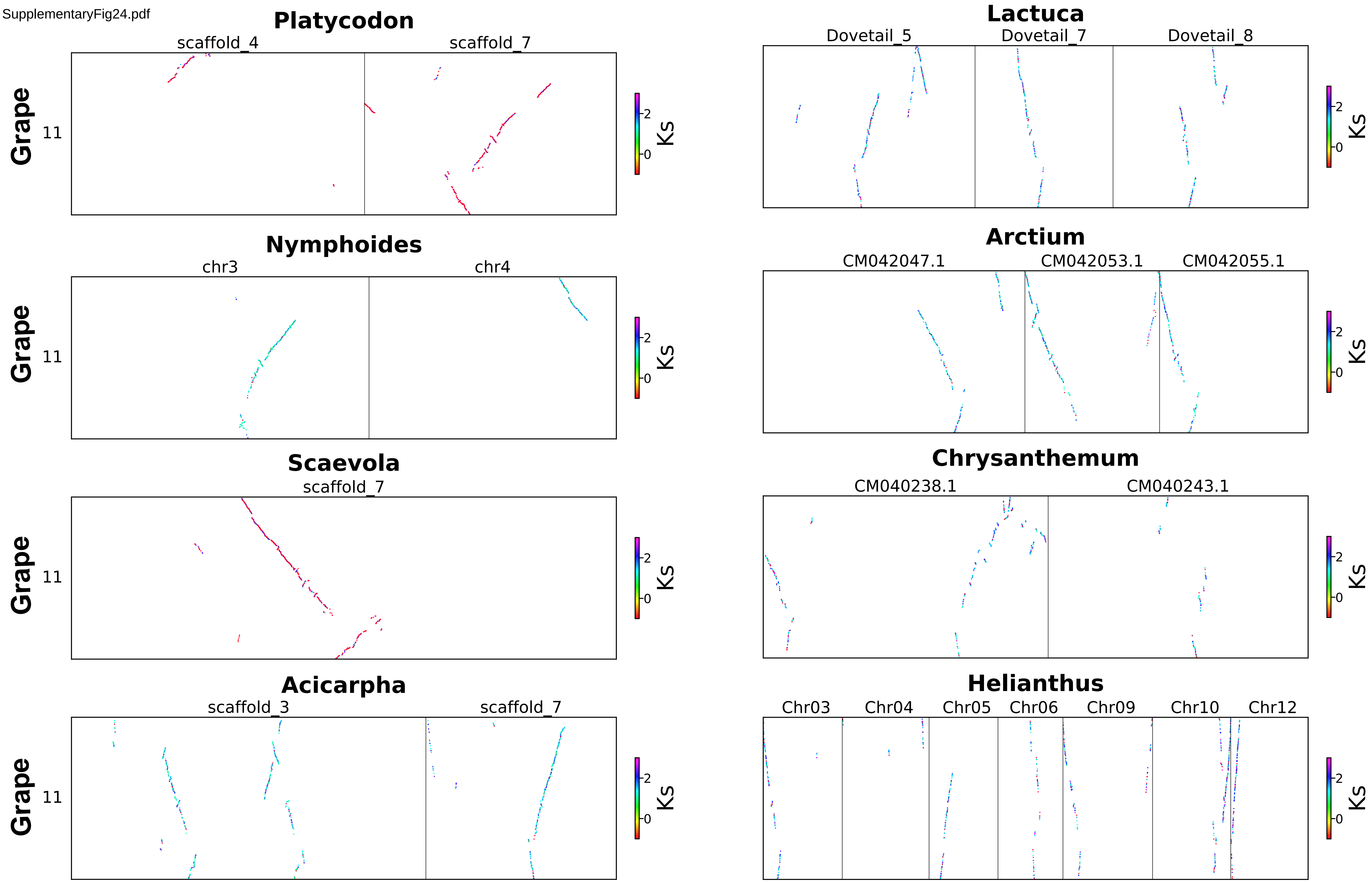

SupplementaryFig25.pdf

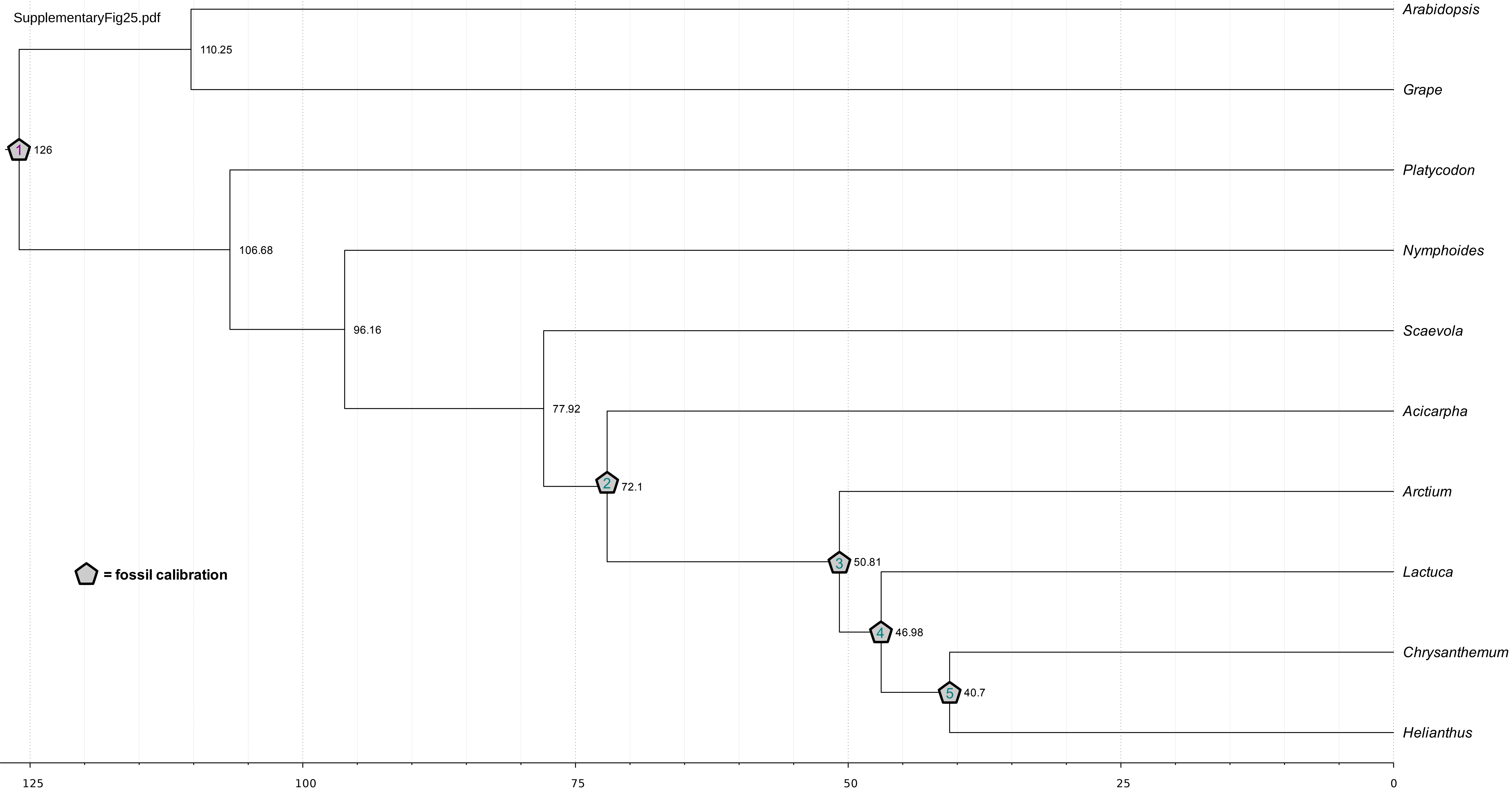

SupplementaryFig26.pdf

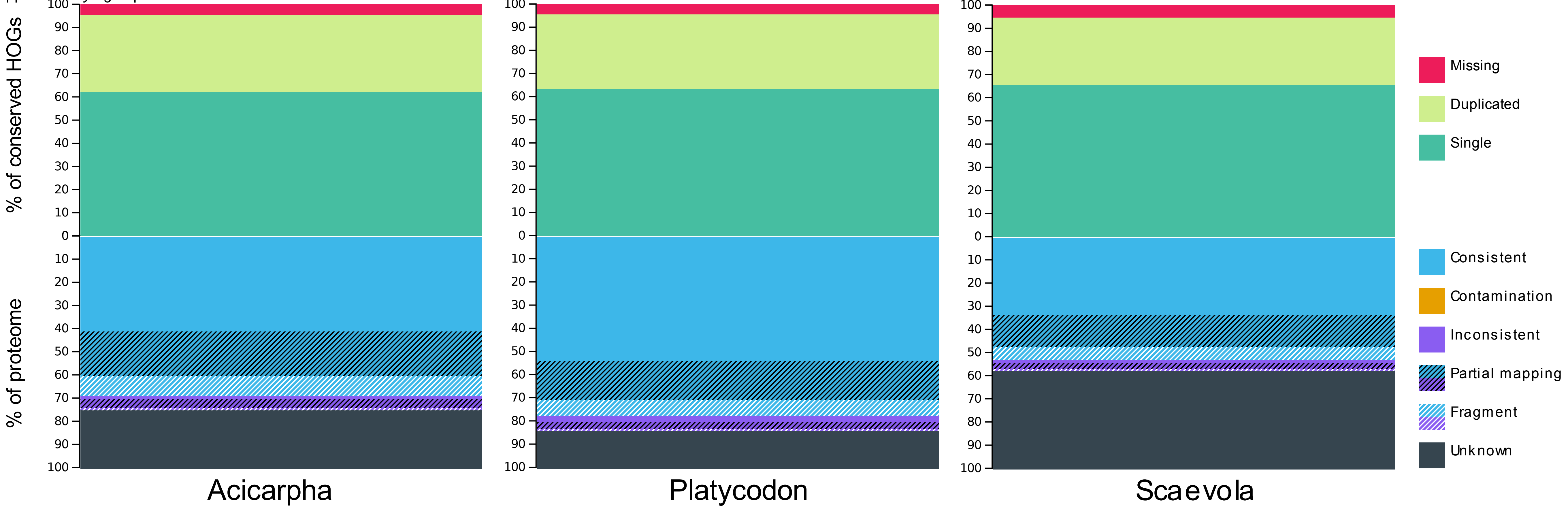

SupplementaryFig27.pdf

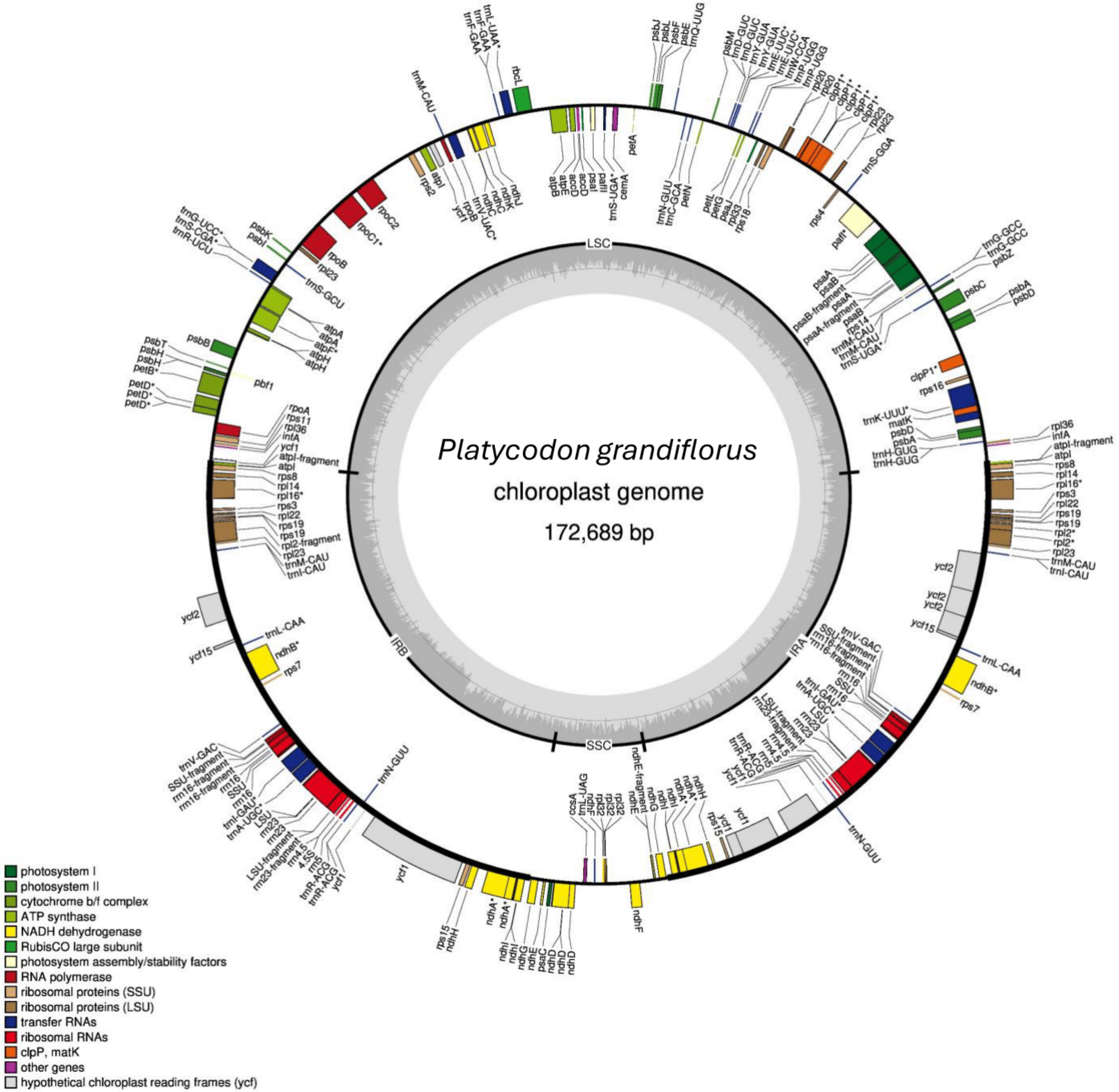


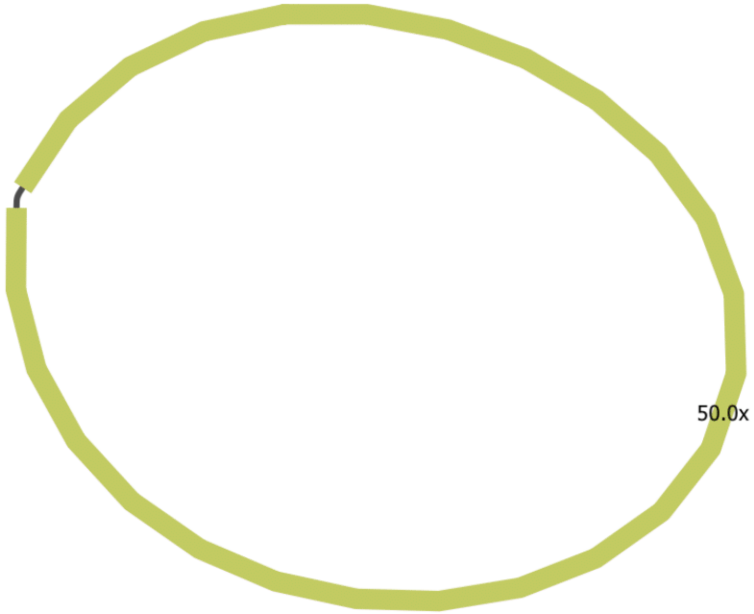

SupplementaryFig28.pdf

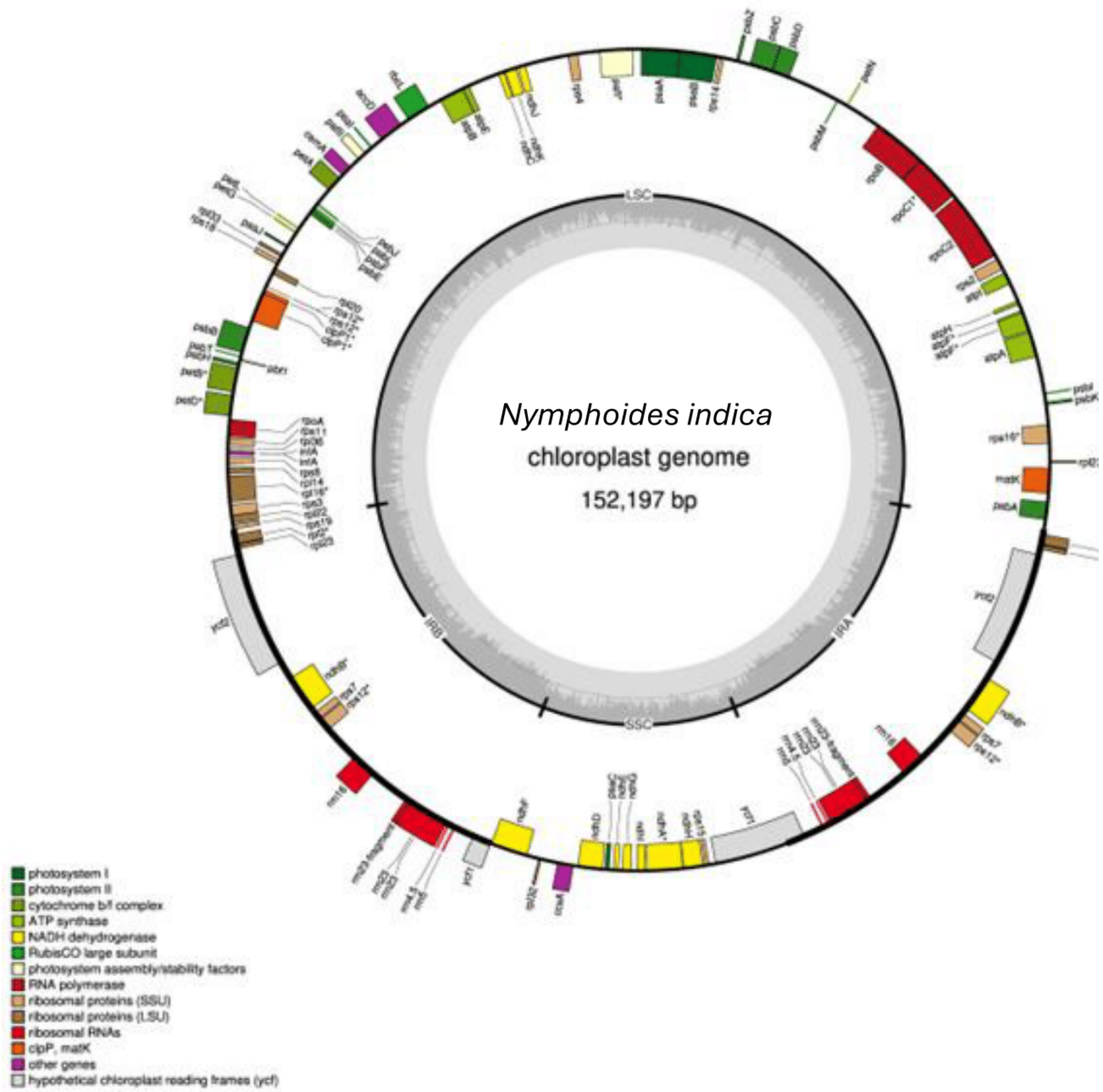


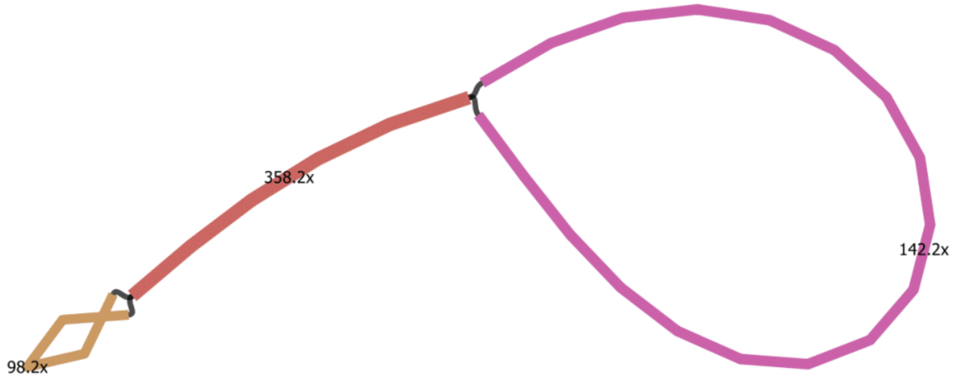

SupplementaryFig29.pdf

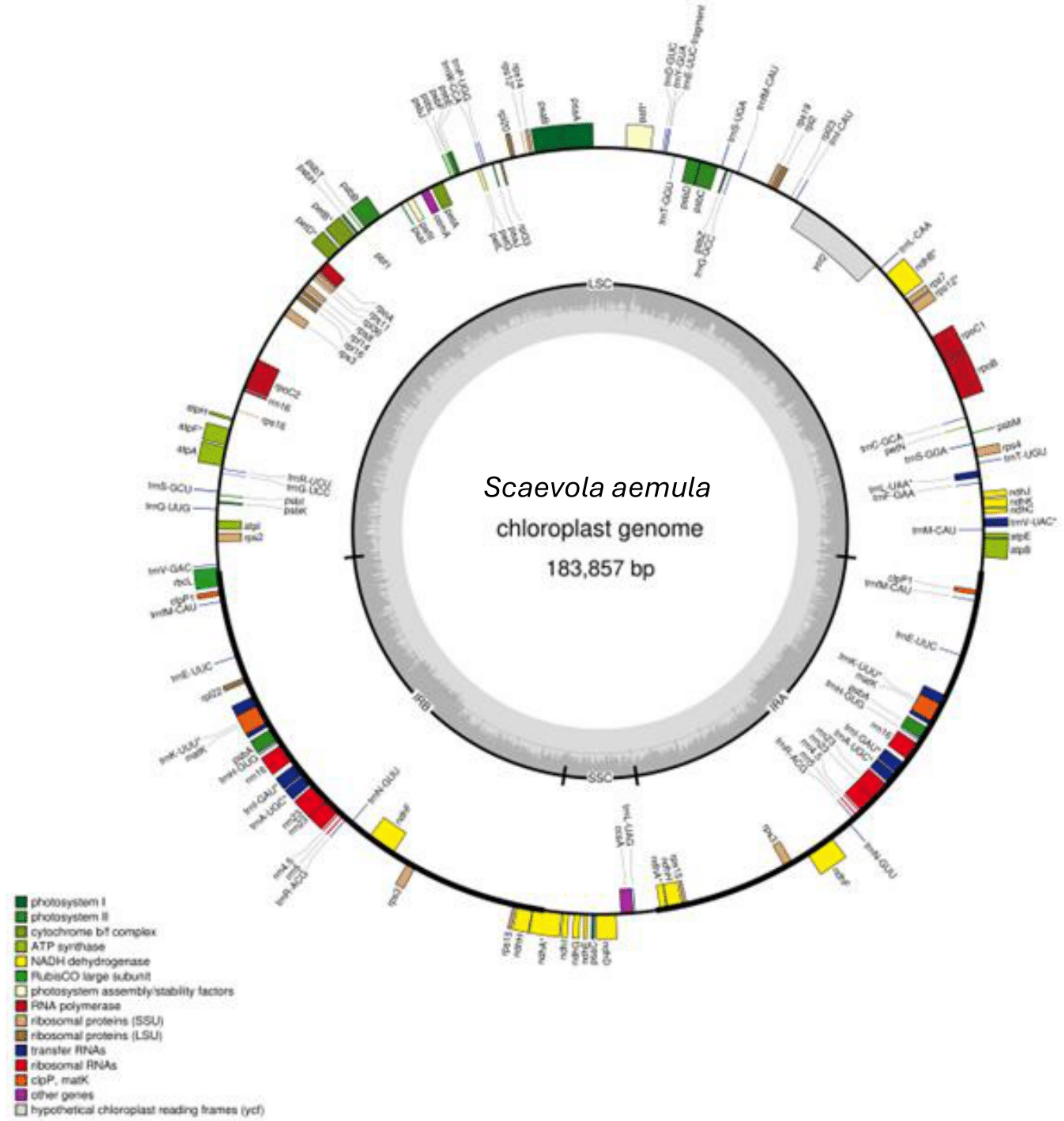


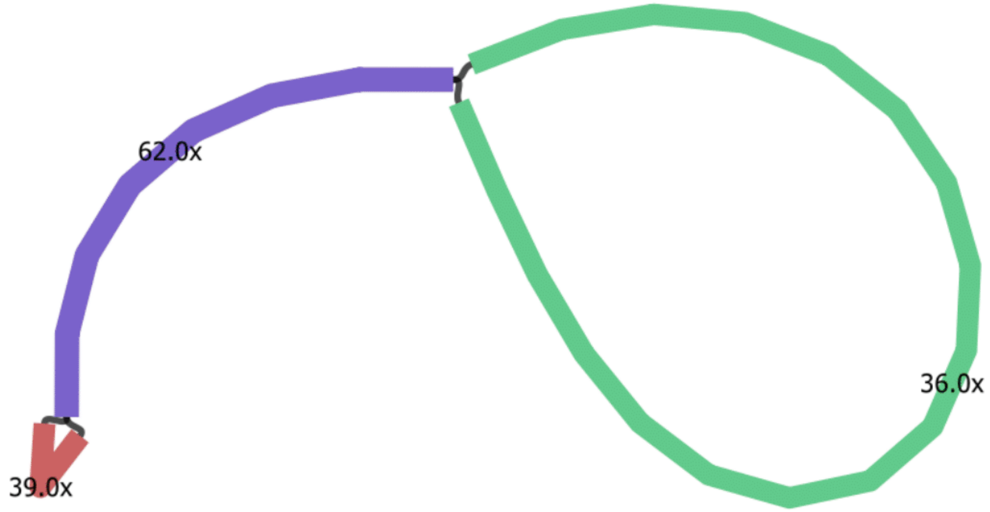

SupplementaryFig30.pdf

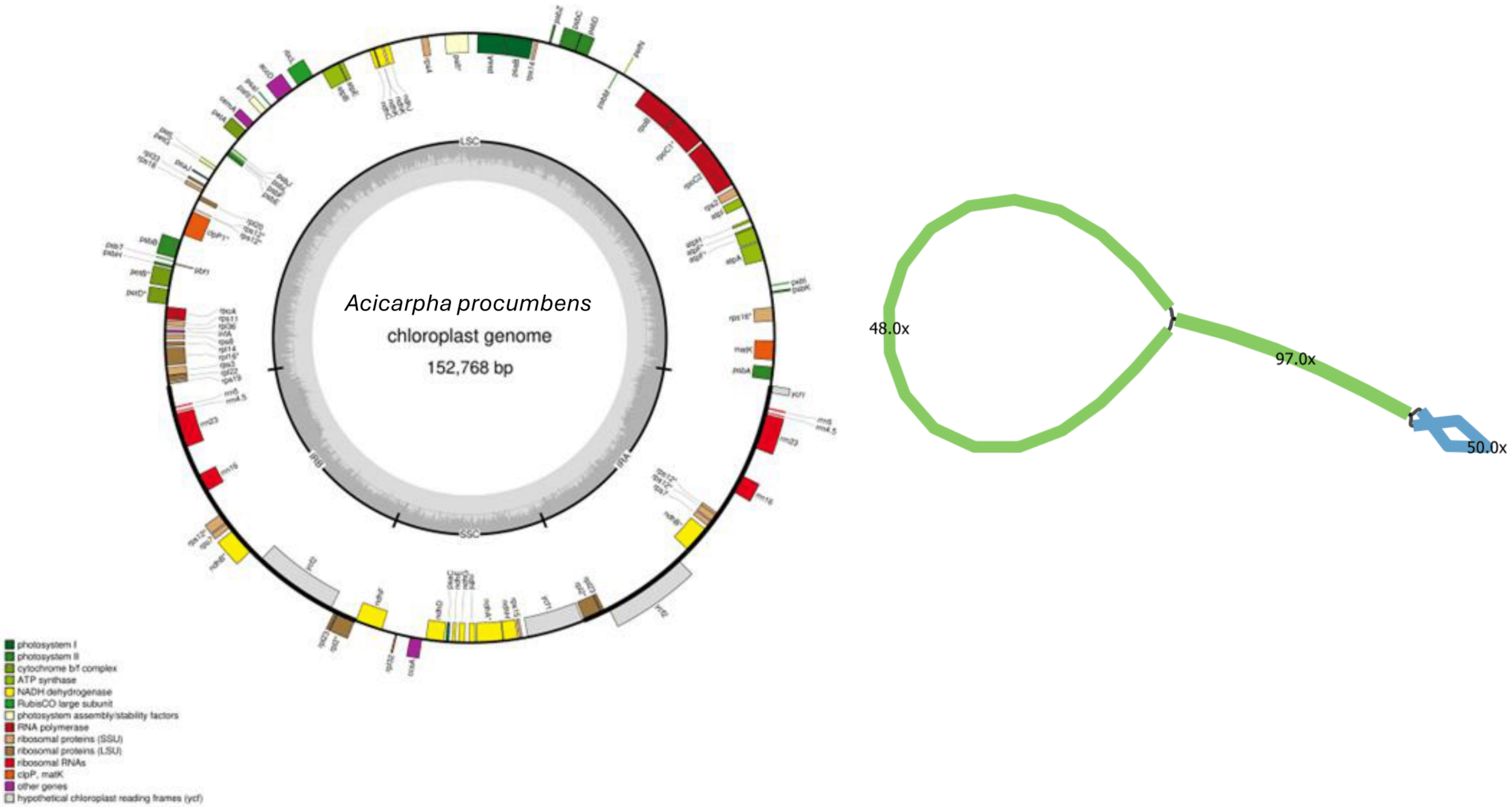

SupplementaryFig31.pdf

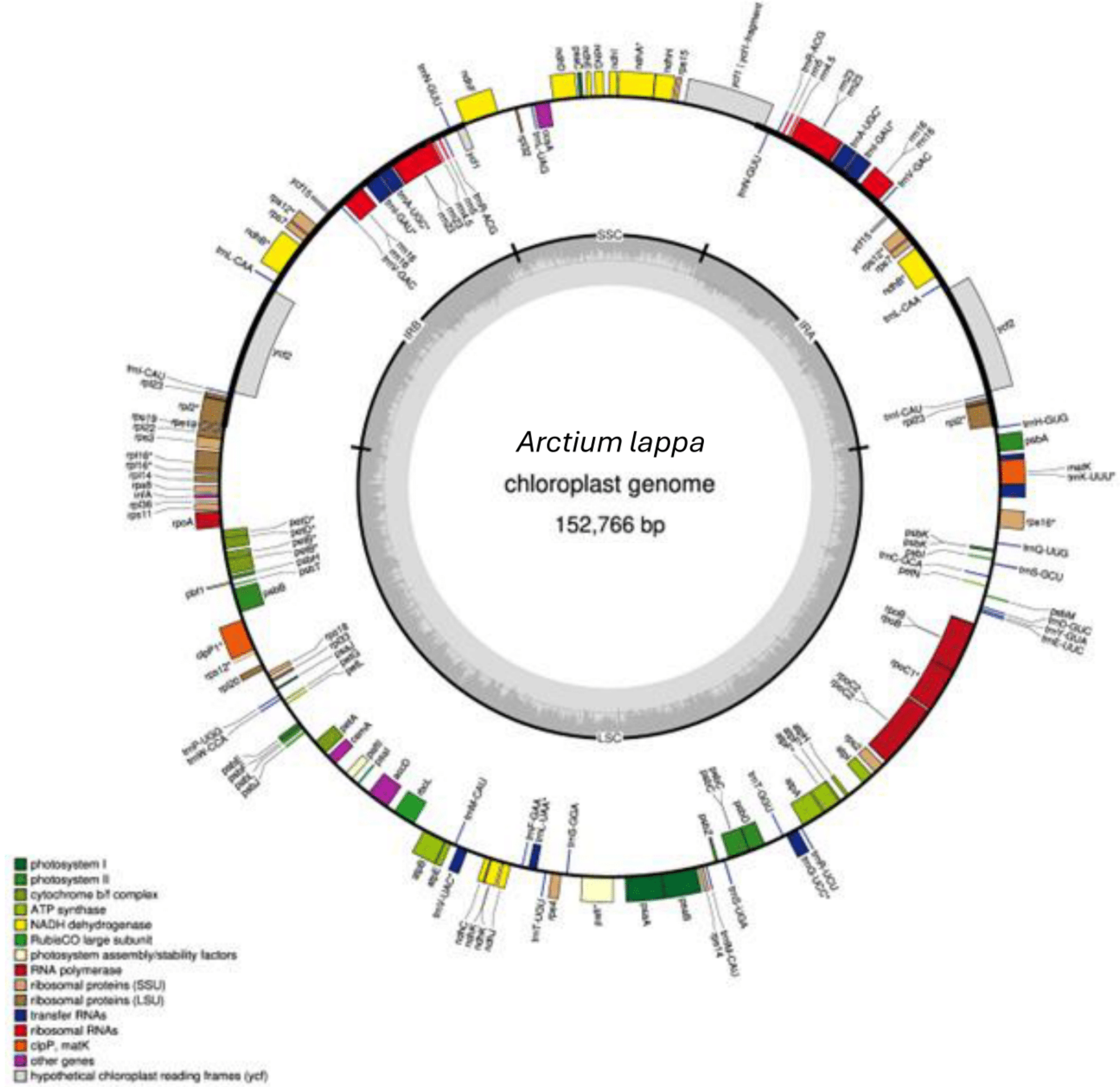


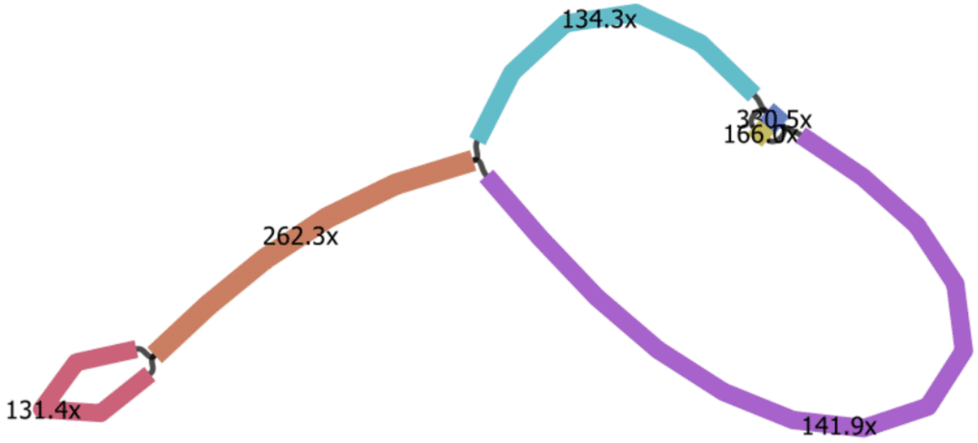

SupplementaryFig32.pdf

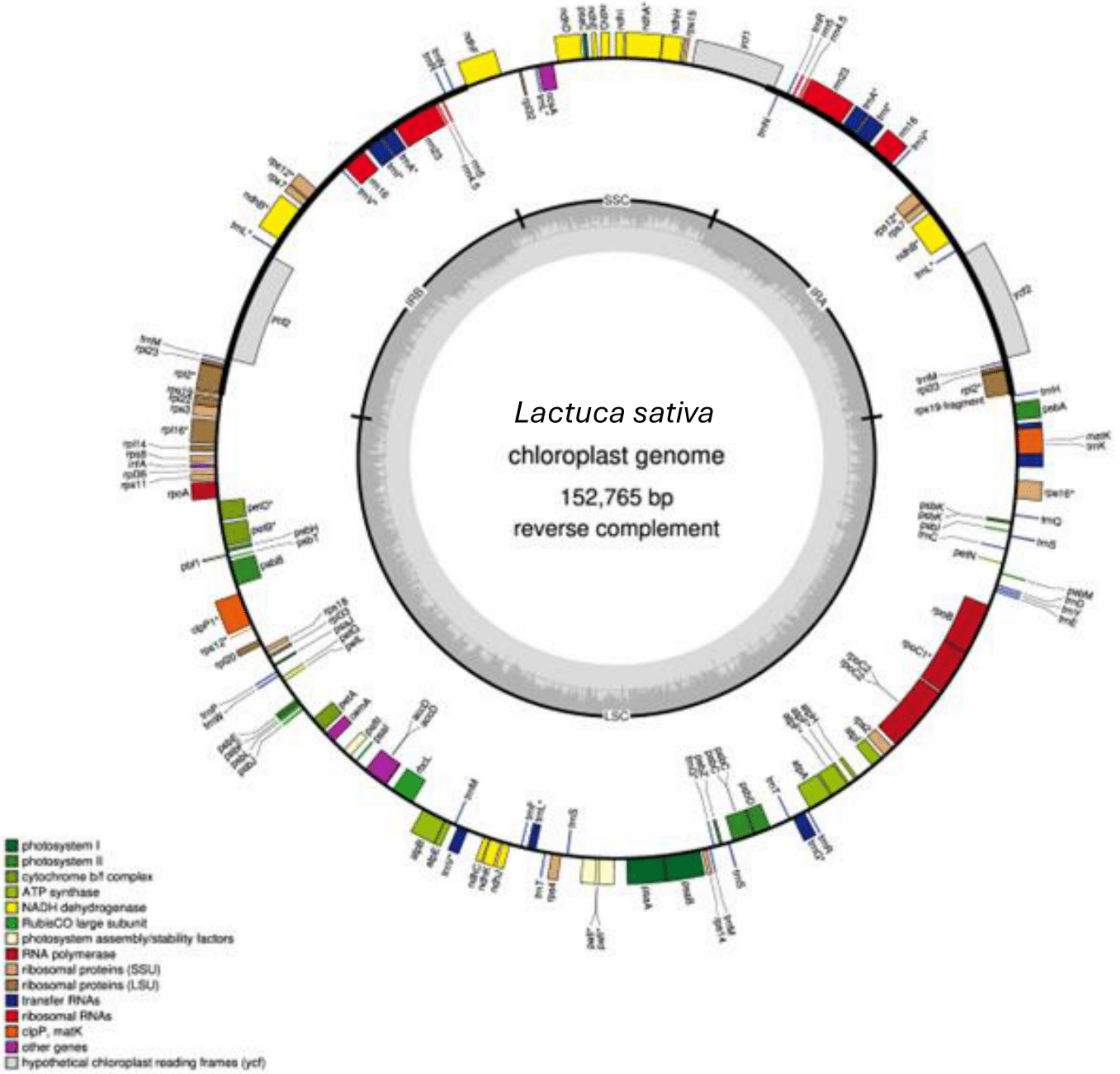


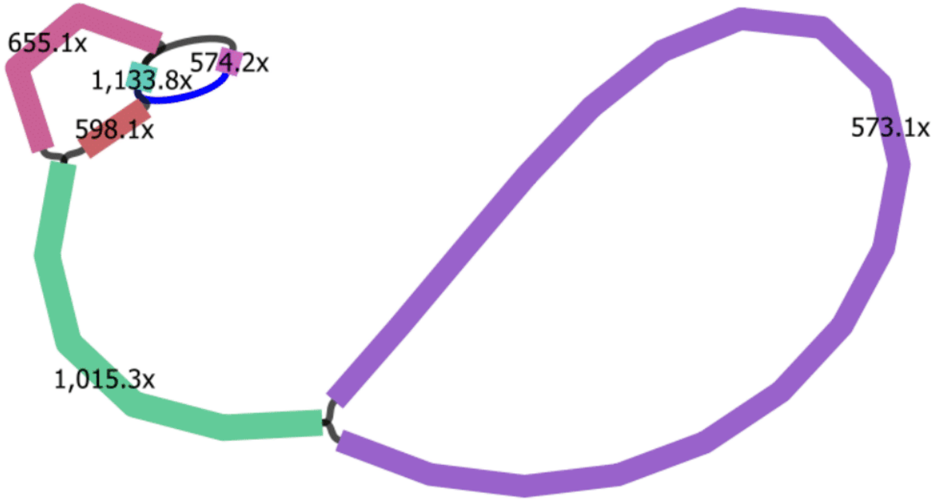

SupplementaryFig33.pdf

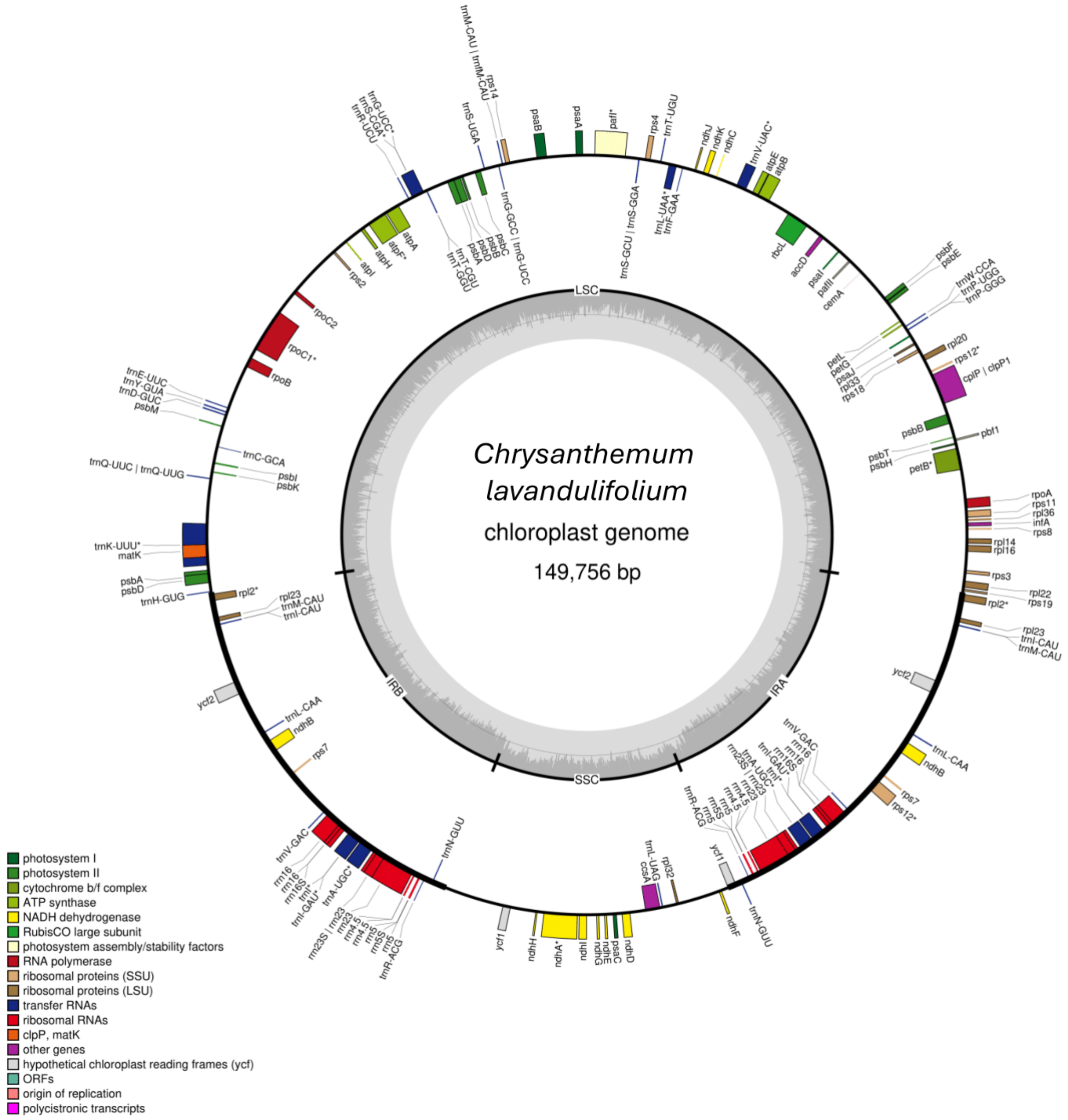


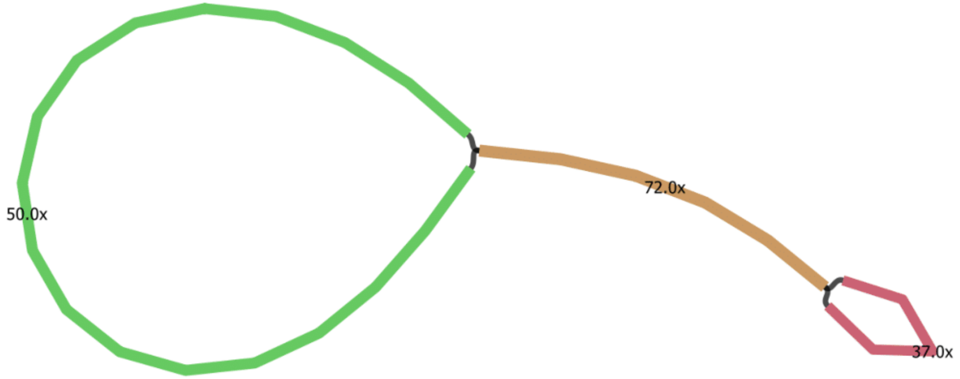

SupplementaryFig34.pdf

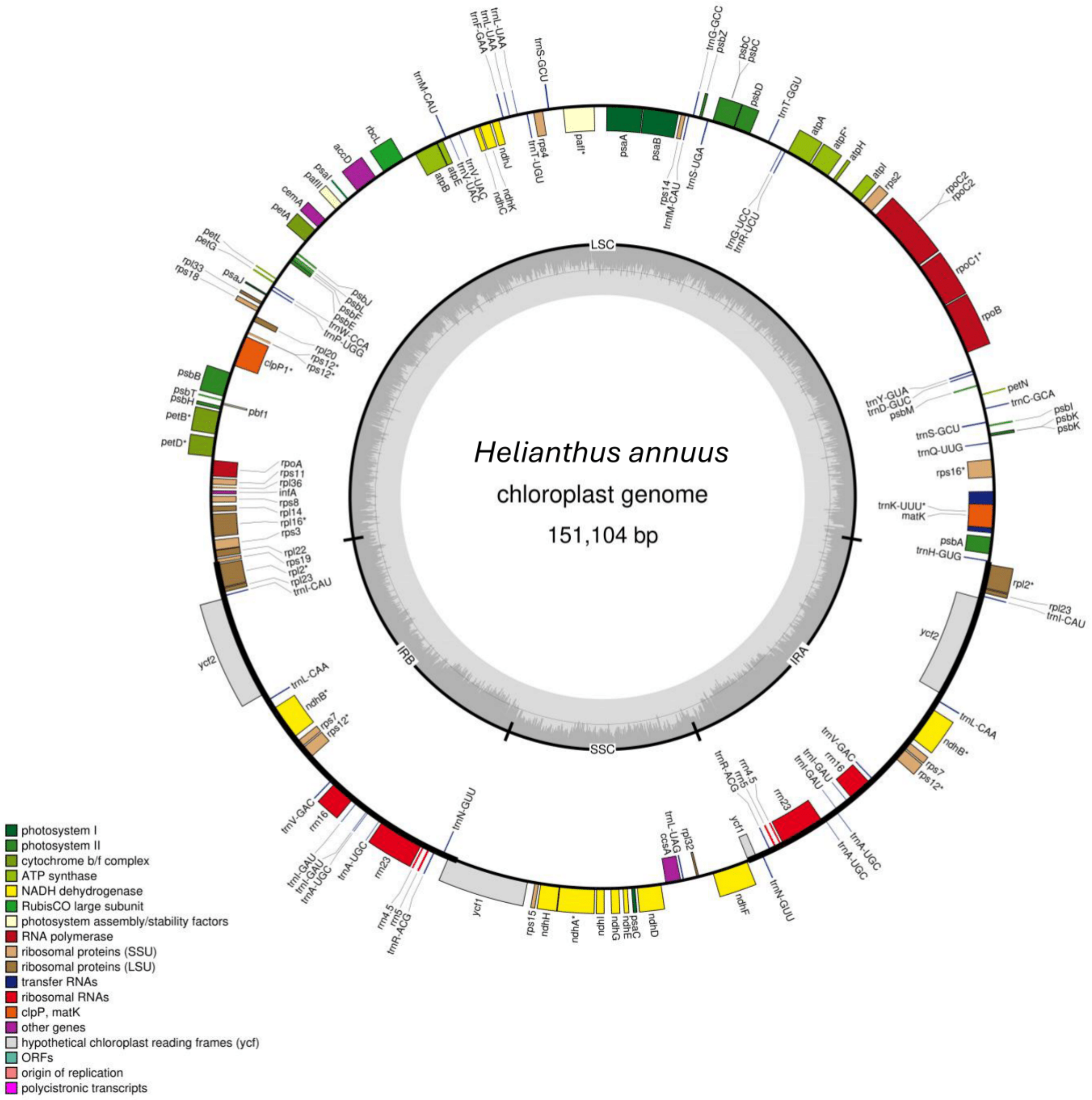

SupplementaryFig35.pdf

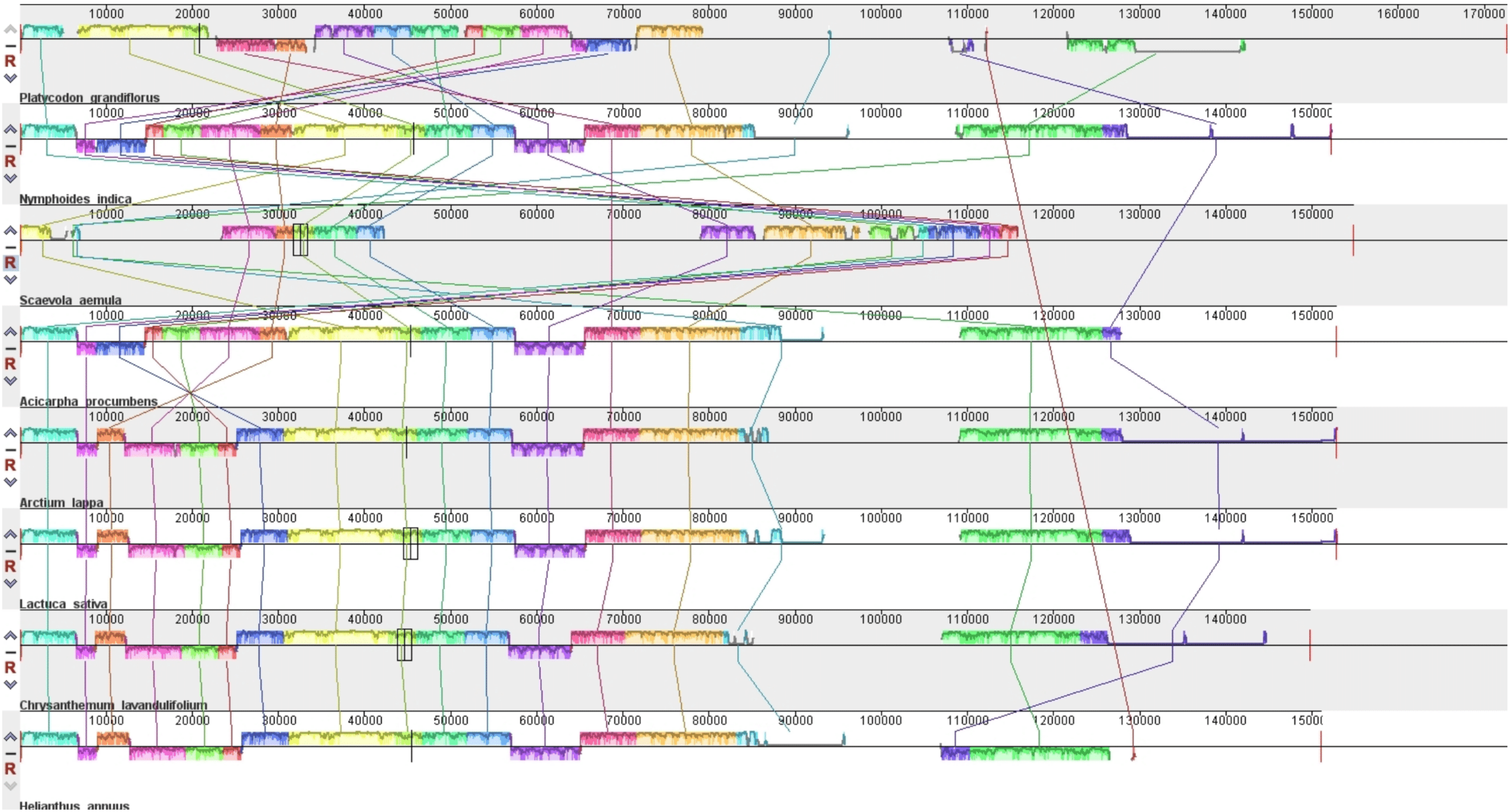

SupplementaryFig36.pdf

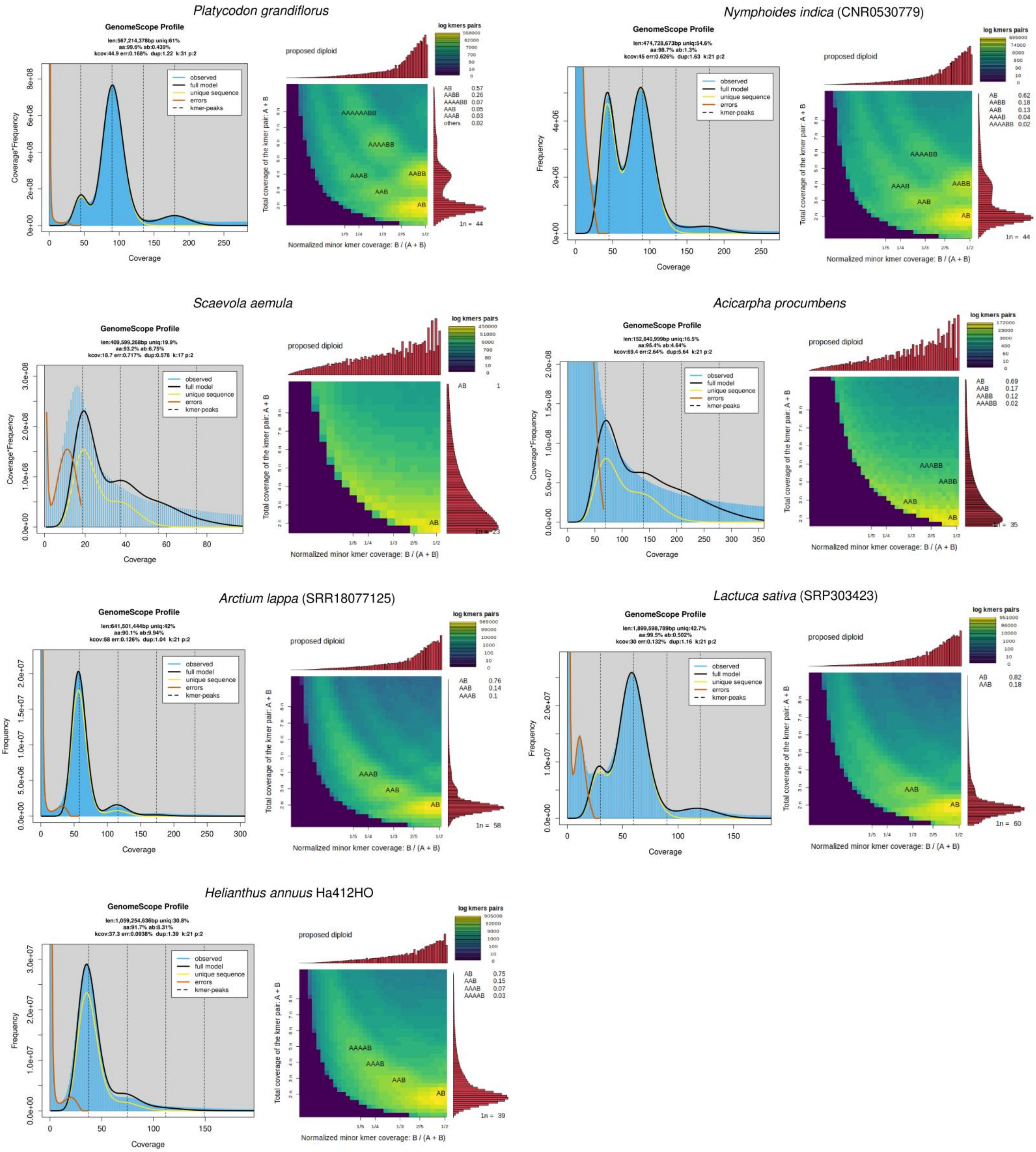

SupplementaryFig37.pdf

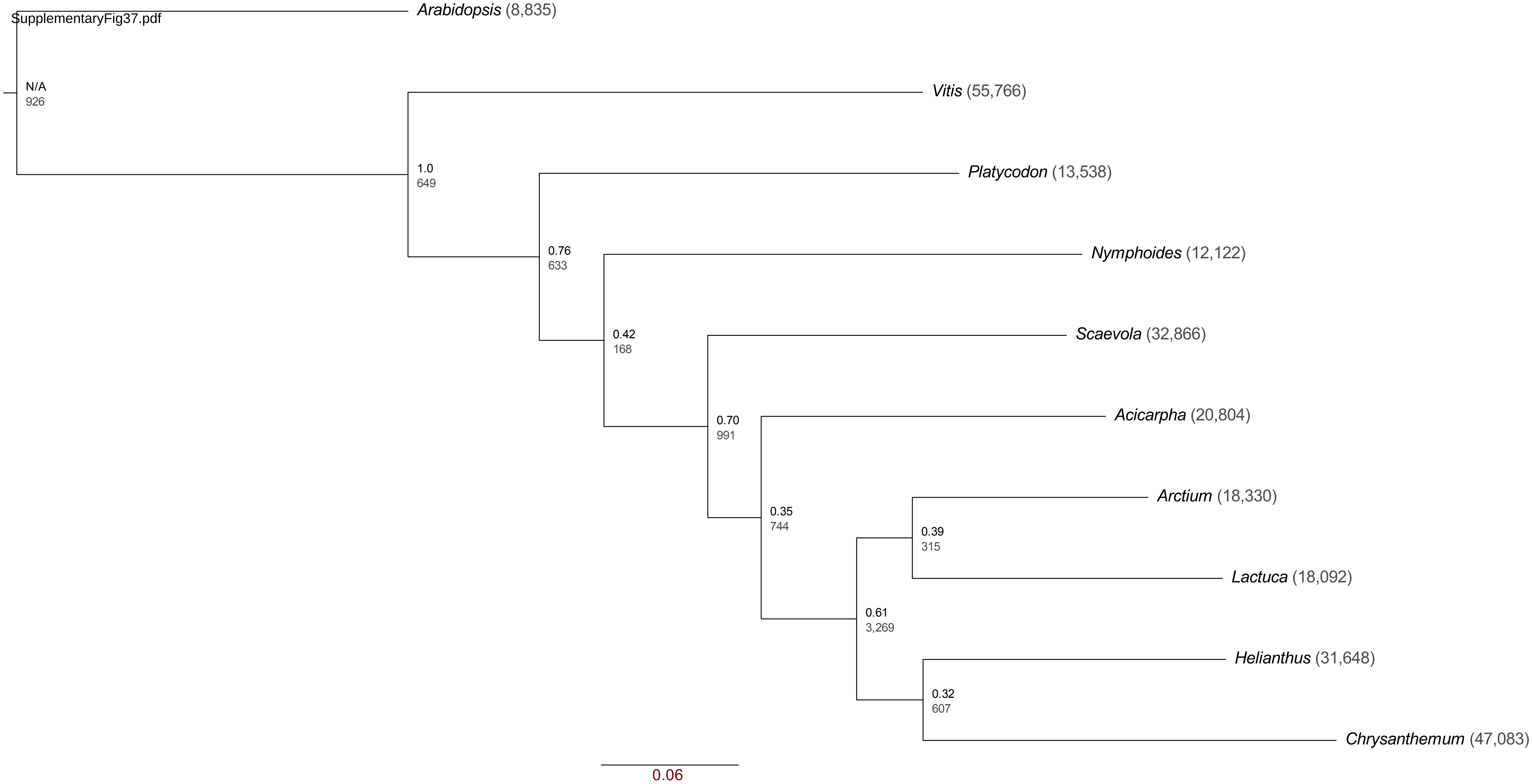

SupplementaryFig38.pdf

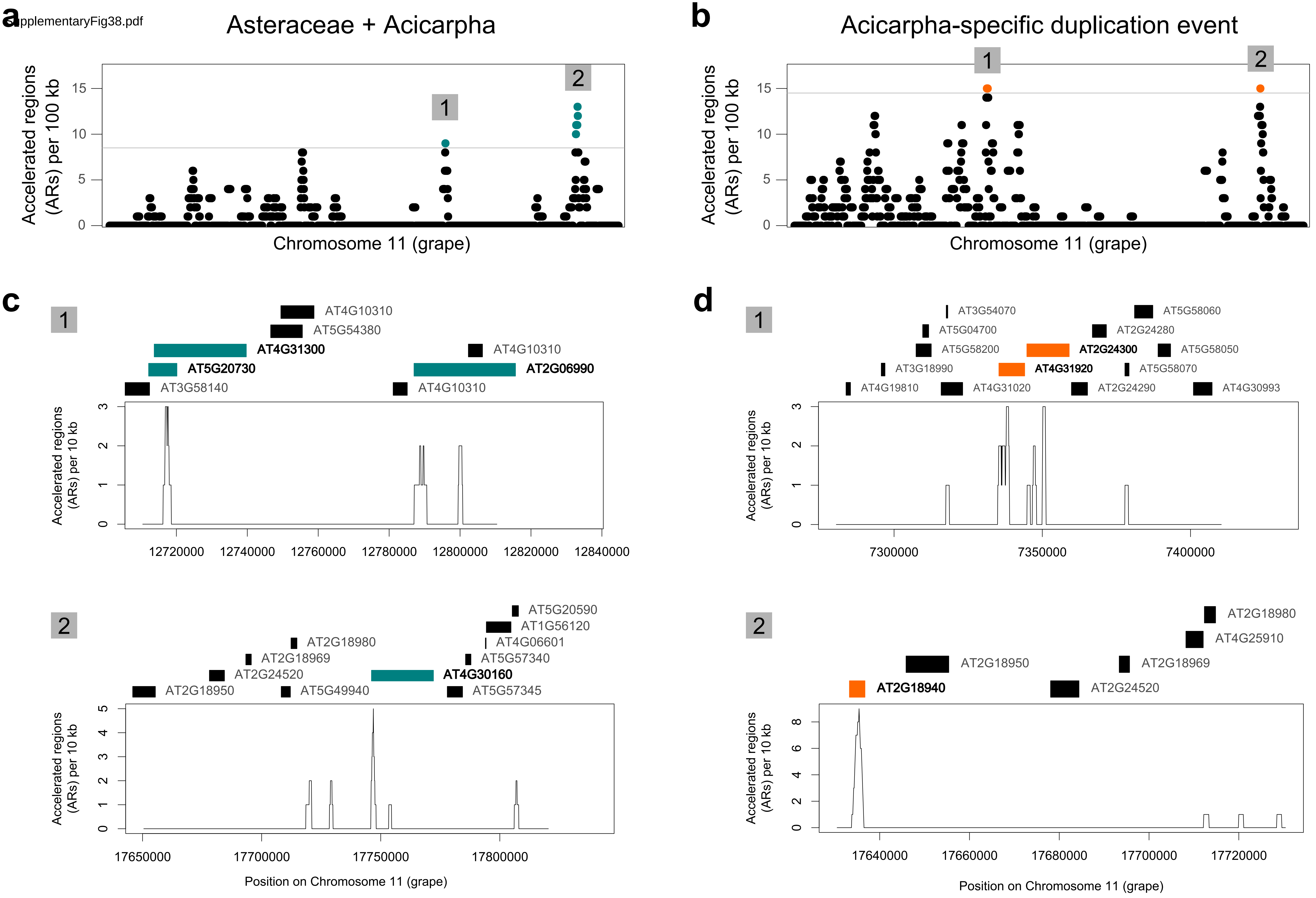
